# From local resynchronization to global pattern recovery in the zebrafish segmentation clock

**DOI:** 10.1101/2020.06.22.166215

**Authors:** Koichiro Uriu, Bo-Kai Liao, Andrew C. Oates, Luis G. Morelli

## Abstract

Rhythmic spatial gene expression patterns termed the segmentation clock regulate vertebrate body axis segmentation during embryogenesis. The integrity of these patterns requires local synchronization between neighboring cells by Delta-Notch signaling and its inhibition results in defective segment boundaries. The oscillating tissue deforms substantially throughout development, but whether such tissue-scale morphogenesis complements local synchronization during pattern generation and segment formation is not understood. Here, we investigate pattern recovery in the zebrafish segmentation clock by washing out a Notch inhibitor, allowing resynchronization at different developmental stages, and analyzing the recovery of normal segments. Although from previous work no defects are expected after recovery, we find that washing out at early stages causes a distinctive intermingling of normal and defective segments, suggesting unexpectedly large fluctuations of synchrony before complete recovery. To investigate this recovery behavior, we develop a new model of the segmentation clock combining key ingredients motivated by prior experimental observations: coupling between neighboring oscillators, a frequency profile, a gradient of cell mixing, tissue length change, and cell advection pattern. This model captures the experimental observation of intermingled normal and defective segments through the formation of persistent phase vortices of the genetic oscillators. Experimentally observed recovery patterns at different developmental stages are predicted by temporal changes of tissue-level properties, such as tissue length and cell advection pattern in the model. These results suggest that segmental pattern recovery occurs at two scales: local pattern formation and transport of these patterns through tissue morphogenesis, highlighting a generic mechanism of pattern dynamics within developing tissues.

**SIGNIFICANCE:** Interacting genetic oscillators can generate a coherent rhythm and a tissue-level pattern from an initially desynchronized state. Using experiment and theory we study resynchronization and pattern recovery of the zebrafish segmentation clock, which makes the embryonic body segments. Experimental perturbation of intercellular signaling with an inhibitor results in intermingled normal and defective segments. According to theory, this behavior may be caused by persistent local vortices scattered in the tissue during pattern recovery. Full pattern recovery follows dynamic global properties, such as tissue length and advection pattern, in contrast to other genetic oscillators in a static tissue such as circadian clocks. Our work highlights how dynamics of tissue level properties may couple to biochemical pattern formation in tissues and developing embryos.

## INTRODUCTION

Synchronization of genetic oscillations in tissues generates robust biological clocks. To attain synchrony, cells interact with each other locally and adjust their phase of oscillations. How local interactions between oscillators lead to the emergence of collective rhythms has been studied in static tissues and in dynamic tissues with local cell rearrangements, but how collective rhythms are influenced by the more complex deformations of entire tissues typical in embryogenesis remains challenging and is less well understood. A system to explore this is the synchronization of genetic oscillations during the segmentation of the vertebrate embryo’s body axis, a process termed somitogenesis. Cells in the unsegmented tissue, presomitic mesoderm (PSM) and tailbud, show collective rhythms of gene expression that set the timing of somite boundary formation and are referred to as the segmentation clock (1, 2). In the tailbud, spatially homogeneous oscillations can be observed. In the PSM, kinematic phase waves of gene expression move from posterior to anterior, indicating the presence of a spatial phase gradient along the axis (3-5). Importantly, this unsegmented oscillating tissue undergoes extensive deformations during the time of segment formation, with complex cellular rearrangements, flows and a changing global size and geometry (6-10). However, our current understanding of synchronization in the segmentation clock follows largely from considering a non-deforming tissue with constant size.

In the zebrafish segmentation clock, Her1 and Her7 proteins repress their own transcription, forming negative feedback loops (11-13). These negative feedback loops are considered to generate cell-autonomous rhythms of gene expression (14-16). Cells in the PSM interact with their neighbors via Delta-Notch signaling (17-20). It is thought that Her proteins repress the transcription of *deltaC* mRNA, causing oscillatory expression of DeltaC protein on the cell’s surface (17, 21). Binding of a Delta ligand to a Notch receptor expressed by neighboring cells leads to the cleavage and release of the Notch intracellular domain (NICD), which is translocated to the nucleus and modulates transcription of *her* mRNAs.

Several lines of evidence based on the desynchronization of the segmentation clock show that Delta-Notch signaling couples and thereby synchronizes neighboring genetic oscillators in the zebrafish PSM and tailbud. The first collective oscillation of the segmentation clock occurs immediately before the onset of gastrulation at 4.5 hours post fertilization (hpf), independently of Delta-Notch signaling (19, 22). Thereafter, cells from embryos deficient in Delta-Notch signaling gradually become desynchronized due to the presence of various sources of noise (17, 23). Single-cell imaging of a live Her1 reporter in the Delta-Notch mutant embryos *deltaC*/*beamter, deltaD*/*after eight* and *notch1a*/*deadly seven* during posterior trunk formation (∼15 hpf) shows that Her1 protein oscillation is desynchronized across the PSM cells (3). At the tissue level, Delta-Notch mutants form the anterior 4∼6 segments normally, followed by consecutive defective segments (24). These phenotypes are not caused by a direct failure of segment boundary formation (25), but have been explained in terms of the underlying desynchronization of the segmentation clock (18, 19).

This desynchronization hypothesis has been formalized as a theory based on coupled oscillators (19, 26). The theory postulates a critical value *Z*_*c*_ such that if the level of synchrony becomes lower than this critical value, a defective segment boundary is formed, Fig. 1A. In the absence of Delta-Notch signaling, the level of synchrony decays over time and eventually becomes lower than *Z*_*c*_. The time point at which the level of synchrony crosses *Z*_*c*_ for the first time is considered to set the anterior limit of defects (ALD), i.e. the anterior-most defective segment along the body axis. Indeed, theory based on this desynchronization hypothesis can quantitatively explain the ALD in Delta-Notch mutants (19).

**Figure 1.**
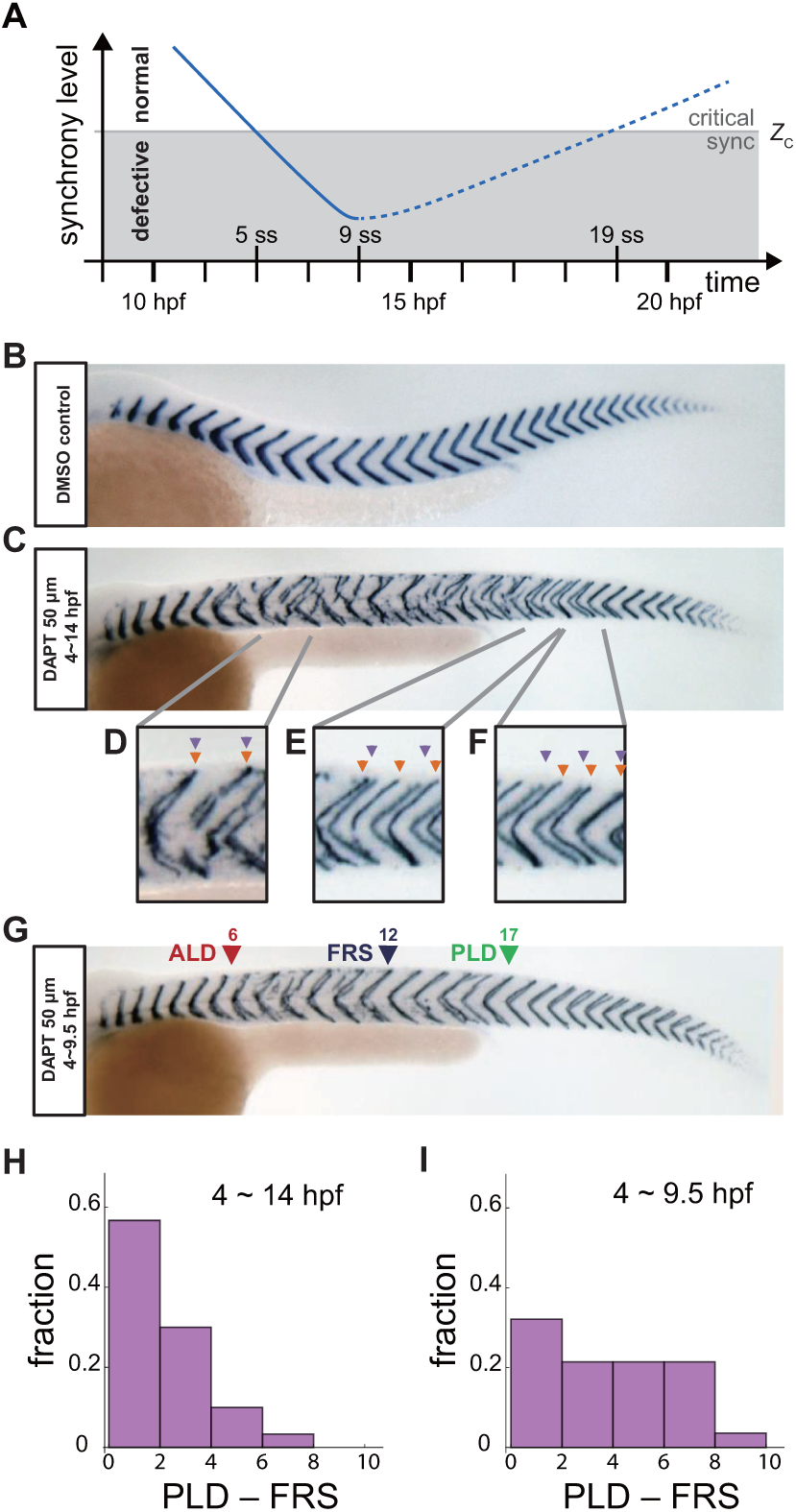
Segment boundary defects observed in late and early DAPT washout embryos. (A) Schematic time series of synchrony level during desynchronization and resynchronization. In the presence of DAPT, the synchrony level decreases due to the loss of Delta-Notch signaling (solid line). DAPT is washed out at 14 hours post fertilization (hpf; ∼9 somite stage; ss) in this panel and resynchronization starts from that time point (dotted line). If the synchrony level is higher (lower) than a critical value *Z*_*c*_, normal (defective) segments are formed. (B) Wild type control embryo treated with DMSO. (C) Embryo with late DAPT washout at 14 hpf (9 ss). Enlargements of (D) broken or fragmented boundaries, (E) incorrect number of boundaries and (F) left-right misaligned boundaries are shown below. (G) Embryo with early DAPT washout at 9.5 hpf (0 ss). Red, blue and green triangles indicate the anterior limit of defect (ALD), first recovered segment (FRS) and posterior limit of defect (PLD), respectively. (H), (I) Histograms of the difference between PLD and FRS (PLD – FRS) for embryos with DAPT washout at (H) late (14 hpf; *n* = 30) and early (9.5 hpf; *n* = 28) stages. Numbers of embryos examined in (H) and (I) were 15 and 14, respectively. FRS and PLD were measured separately between left and right sides of embryos.

The desynchronization hypothesis implies that the oscillators could be resynchronized by restoring Delta-Notch signaling, with the expectation that resynchronization of the segmentation clock requires several oscillation cycles for the level of synchrony to smoothly increase and surpass the threshold *Z*_*c*_, giving rise to a transition from defective to recovered normal segments. Due to the constitutive lack of coupling, Delta-Notch mutants cannot be used to analyze resynchronization dynamics. A powerful tool for this purpose is the Notch signal inhibitor DAPT, which inhibits the cleavage and release of NICD, blocking cell coupling. Importantly, DAPT can be washed out to recover cell coupling and this allows cells to resynchronize their genetic oscillations, Fig. 1A (19, 25-27). Previous experimental studies showed that after late washout of DAPT at the 9 somite stage, embryos start making normal segments again after several oscillation cycles (19, 26, 27). The position of the first recovered normal segment after DAPT washout represents the time at which the level of synchrony surpasses *Z*_*c*_ for the first time (26). In previous studies, almost all segments formed normally after the first fully recovered segment (19, 26), consistent with a monotonic increase of the level of synchrony in the vicinity of *Z*_*c*_, Fig. 1A, as expected from the desynchronization hypothesis. Despite this success, it remains fundamentally unclear how tissue-scale gene expression patterns underlying segment recovery reorganize from local intercellular interactions and whether they are affected by tissue size and shape changes that occur during development.

Here we analyze resynchronization dynamics of the zebrafish segmentation clock at different developmental times using both experimental and theoretical techniques. In contrast to late washout mentioned above, we find that washing out DAPT at earlier developmental stages causes a region of scattered segment defects, where normal and defective segments are intermingled. This striking phenotype was not anticipated by previous models (19, 28). To investigate the processes in the segmentation clock that yield this distinctive recovery behavior, here we develop a new model of the segmentation clock that encompasses two scales, describing resynchronization and pattern recovery in terms of local interactions between cells in the tissue, as well as global properties of the tissue. In concert, we develop observables that allow pattern dynamics to be quantified and compared between simulation and experiment, the vorticity index and local order parameter. Despite its simplicity, this model can describe the intermingling of normal and defective segments. Numerical simulations indicate that persistent phase vortices appear during resynchronization, resulting in scattered and intermingled segment defects along the axis. The length of the PSM and tailbud and advection pattern influence the recovery process via the transport of phase vortices from the posterior to the anterior of the PSM. Moreover, by including temporal changes to tissue length, advection pattern and coupling strength, the model makes predictions about the pattern of resynchronization at both early and late stages that we confirm experimentally in the embryo.

## RESULTS

### Scattered defective segments in resynchronization assay

To investigate the processes involved in resynchronization of the segmentation clock, we used a resynchronization assay based on washing out the Notch signaling inhibitor DAPT at different developmental stages. In this assay, zebrafish embryos were placed in DAPT for a defined duration, during which time the segmentation clock desynchronized and defective segments were formed, then washed extensively to allow Delta-Notch signaling activity to resume. Subsequently, the segmentation clock gradually resynchronized and normal segments were made.

Throughout this study, we administered DAPT at 4 hpf, a developmental stage before the oscillating genes of the segmentation clock were expressed. This was a treatment duration of at least 5 hours, sufficient to obtain defects on both left and right sides of the embryo for subsequent resynchronization analysis (25). A record of the resulting spatiotemporal pattern of somitogenesis was visualized after its completion by whole-mount *in situ* hybridization for the myotome segment boundary marker gene *xirp2a* (*xin actin-binding repeat containing 2a*) in ∼36 hpf embryos, Fig. 1B, C. We analyzed these staining patterns by scoring boundaries as either normal or defective using established criteria (19, 26), with the exception that we scored the left and right embryonic sides separately. Examples of defects observed with DAPT treatment included fragmented, mis-spaced or mis-aligned boundaries, Fig. 1D-F. To prevent incorrect identification of misaligned boundaries in embryos with bent axes, or in tilted samples, we confirmed that the boundaries outside of the defective region were well aligned between left and right sides. To assign the ordinal segment number to defective boundaries when boundaries were severely fragmented, we used the contralateral side or counted either dorsal or ventral boundary ends, which were often clearer, to estimate their axial position.

As described in the introduction, the location of the transition from normal to defective segments resulting from desynchronization is termed the anterior limit of defects (ALD), given by the first segment along the embryo’s axis that shows a defective boundary, Fig. S1A. After removing DAPT, resynchronization begins and normal segments form eventually. This transition has been recorded by the first recovered segment along the axis (FRS) (26) and then the posterior limit of defects (PLD), the most posterior segment along the axis with a boundary defect, Fig. S1A (19). Note that because segments form rhythmically and sequentially along the body axis, FRS and PLD label both an axial position and the developmental time of segment formation.

In late washout experiments, we observed that a normal segment boundary sometimes formed shortly after ALD even when DAPT was still present, possibly due to desynchronization fluctuations. In previous reports, the definition of the FRS avoided counting these early defects because washout was done late, after full desynchronization. However, when DAPT was washed out early, before ALD occurred, we could not discriminate whether a normal segment formed due to desynchronization fluctuations or as a consequence of resynchronization. The frequency of defects kept growing during an early phase in all conditions until reaching a plateau around segment 9, Fig. S1B, suggesting that the desynchronization phase lasted until segment 9, at least. Hence, in this study we defined FRS as the first recovered segment after segment 9, when the desynchronization phase was over.

We first analyzed the recovery of normal segments when DAPT was washed out at 14 hpf (*t*_*wash-out*_ = ∼9 somite-stage (ss)), as in previous studies, Fig. 1C. Several defective segment boundaries were formed after washout, suggesting that the level of synchrony was still lower than the critical value for normal segment formation during that time interval. However, embryos recovered a normal segment boundary after some time, indicating that the level of synchrony surpassed the threshold, Fig. 1C. With this late washout time, we often observed contiguous defective segments before FRS, suggesting that cells in the PSM were completely desynchronized by a DAPT pulse of this length. In addition, PLD coincided closely with FRS, with the distribution of PLD – FRS peaking at lowest values, Figs. 1H, S1A, F suggesting that once the level of synchrony surpassed the critical value *Z*_*c*_, it remained larger than *Z*_*c*_, as expected. This observation can be interpreted using the desynchronization hypothesis to indicate that the level of synchrony increases monotonically over time, resulting in the formation of consecutive normal segments after the FRS, Fig. 1A.

Importantly, however, when we washed DAPT out at an earlier time *t*_*wash-out*_ = 9.5 hpf (∼ 0 ss), the majority of embryos had an interval along the axis where normal and defective segments were intermingled between FRS and PLD, with the difference between PLD and FRS distributed more uniformly, Figs. 1G, I, S1A, C. This result suggests that the segmentation clock has a level of synchrony close to *Z*_*c*_ in this intermingled region, and persistent fluctuations in synchrony level lead to intermittent boundary formation.

### Physical model of the PSM

According to the desynchronization hypothesis, intermingling of normal and defective segment boundaries suggests a fluctuation of the phase order parameter around its critical value for proper segment formation. How could such large and potentially long-lasting fluctuations of the phase order occur? The desynchronization hypothesis (18) was first formalized in a mean-field theory describing synchronization dynamics from global interactions (19). Later, synchronization in the tailbud was analyzed with a theory with local interactions and neighbor exchange by cell mobility (28, 29). However, a critical prediction of these theories is that once a population of oscillators is synchronized, a large fluctuation of synchrony level is not expected (30-32). Instead of such global phase order fluctuations, other hypotheses for the intermingled defects are the emergence of localized disorder, or the existence of local phase order with a mis-orientation to the global pattern. To explore these potential behaviors, we develop a physical model of the PSM and tailbud that includes the local processes of phase coupling and physical forces between neighboring oscillators, and also the tissue-level properties of a frequency profile, oscillator arrest front, tissue length, a gradient of cell mixing, and the advection pattern of the PSM, Figs. 2, S2 and Table S1.

**Figure 2.**
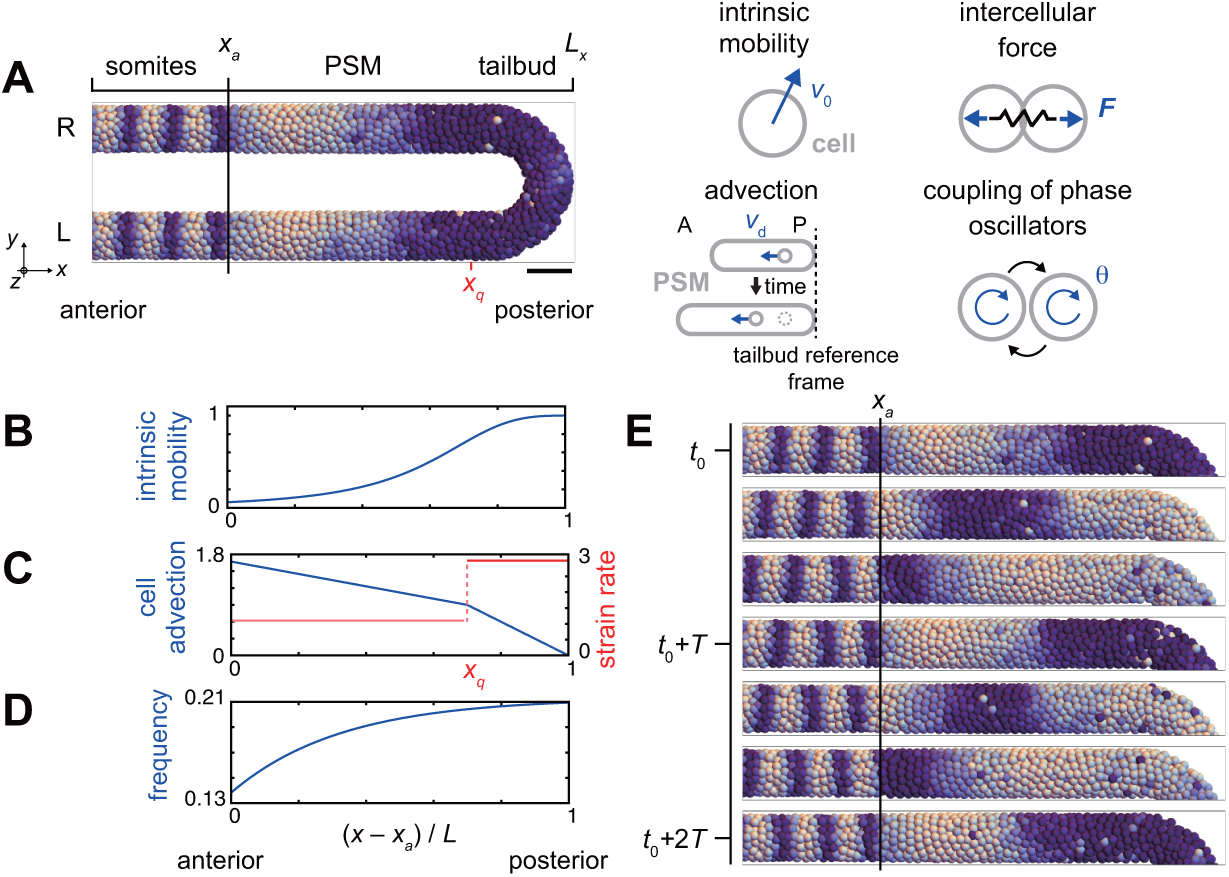
Physical model of the PSM and tailbud. (A) U-shape geometry of the PSM and tailbud (left), and schematics of key ingredients in the model (right). Each sphere represents PSM cells. The color indicates (1 + sin *θ*_*i*_)/2 where *θ*_*i*_ is the phase of oscillation for cell *i*. R: right. L: left. Scale bar: 50 μm. (B) Intrinsic cell mobility gradient, (C) cell advection speed, and (D) autonomous frequency gradient along the anterior-posterior axis of the PSM and tailbud. In (C) the absolute value of the spatial derivative of advection speed, referred to as strain rate, is indicated by the red line. *L* is the length of the PSM *L* = *L*_x_ – *x*_*a*_. (E) Kinematic phase waves moving from the posterior to anterior PSM in a simulation. Snapshots of the right PSM are shown. See also Movie 1. *t*_0_ = 302 min is a reference time point. *T* = 30 min is the period of oscillation at the posterior tip of the tailbud. Parameter values for simulations are listed in Table S1.

The model describes the PSM and tailbud as a U-shaped domain in a 3D space, Figs. 2A, S2, see Materials and Methods in Supporting Information (SI) for details. We set the posterior tip of the tailbud as a reference point. Cells are represented as spheres in the 3D space and subjected to physical forces between neighboring cells and from tissue boundaries (28). In accordance with previous experimental studies, we consider the spatial gradient of intrinsic cell mobility across the PSM and tailbud, with highest mobility in the posterior, Fig. 2B (8, 9, 28). Axis extension as observed in the lab is here described, from the reference point of the posterior tip of the tailbud, as cell advection from posterior to anterior, Fig. 2A, C (7, 33, 34). The value of the spatial derivative of the advection velocity at each position effectively represents the local strain rate, Fig. 2C.

Genetic oscillation in each PSM cell is described as a phase oscillator with noise terms. The phase oscillators are coupled with their neighbors, representing intercellular interaction with Delta-Notch signaling. We define cells to be neighbors when their distance is shorter than a cell diameter. We also consider a frequency profile along the anterior-posterior axis of the PSM, as observed in zebrafish embryos, to create kinematic phase waves, Fig. 2D (4, 5, 7). The frequency of oscillation is highest at the tip of the tailbud and gradually decreases toward the anterior PSM. When a simulation was started from a uniformly synchronized initial condition, the frequency profile generated kinematic phase waves due to growing phase differences between the anterior and posterior PSM, Fig. 2E and Movie 1.

To model the formed segments, we arrest the oscillation when cells leave the PSM from its anterior end *x*_*a*_, Fig. 2A, E. The arrested phase stripes in the region *x* < *x*_*a*_ are representative of segment boundaries, and the segment length is the wavelength of this arrested phase pattern. Although the determination of the segment boundary *in vivo* is a complex process (35, 36), for simplicity we consider that these phase stripes correspond to the boundaries of the resulting morphological segments and the chevron-shaped myotomes that are detected in our experiments. Cells in the formed segment region *x* < *x*_*a*_ continue to be advected anteriorly at the same speed as the anterior end of the PSM, due to axial elongation, and their relative positions are fixed. While small heterogeneities in cell density and cell division may exist in the tissue (9, 10, 28, 37), as a simplifying assumption we keep global cell density constant over time. Consequently, we add new cells with random phase values (17) at random positions in the PSM and tailbud to balance the cells exiting through the anterior end of the PSM.

### Intermingled defective segments result from spatially heterogeneous resynchronization in the PSM

Using this physical model, we analyzed resynchronization dynamics in simulations. As an initial condition, we described the state of the PSM and tailbud immediately after DAPT washout by assigning random initial phases to cells, top panel in Fig. 3A, as an extended treatment of saturating dose of DAPT was expected to cause such complete randomization of oscillator phases (3). In this desynchronized state, normal segment boundaries - as defined by ordered phase stripes – did not form, matching the experimental appearance of embryos with persistent loss of Delta-Notch signaling. For simplicity, we start our analysis with constant tissue parameters. Below, we will introduce temporal changes to the parameters.

**Figure 3.**
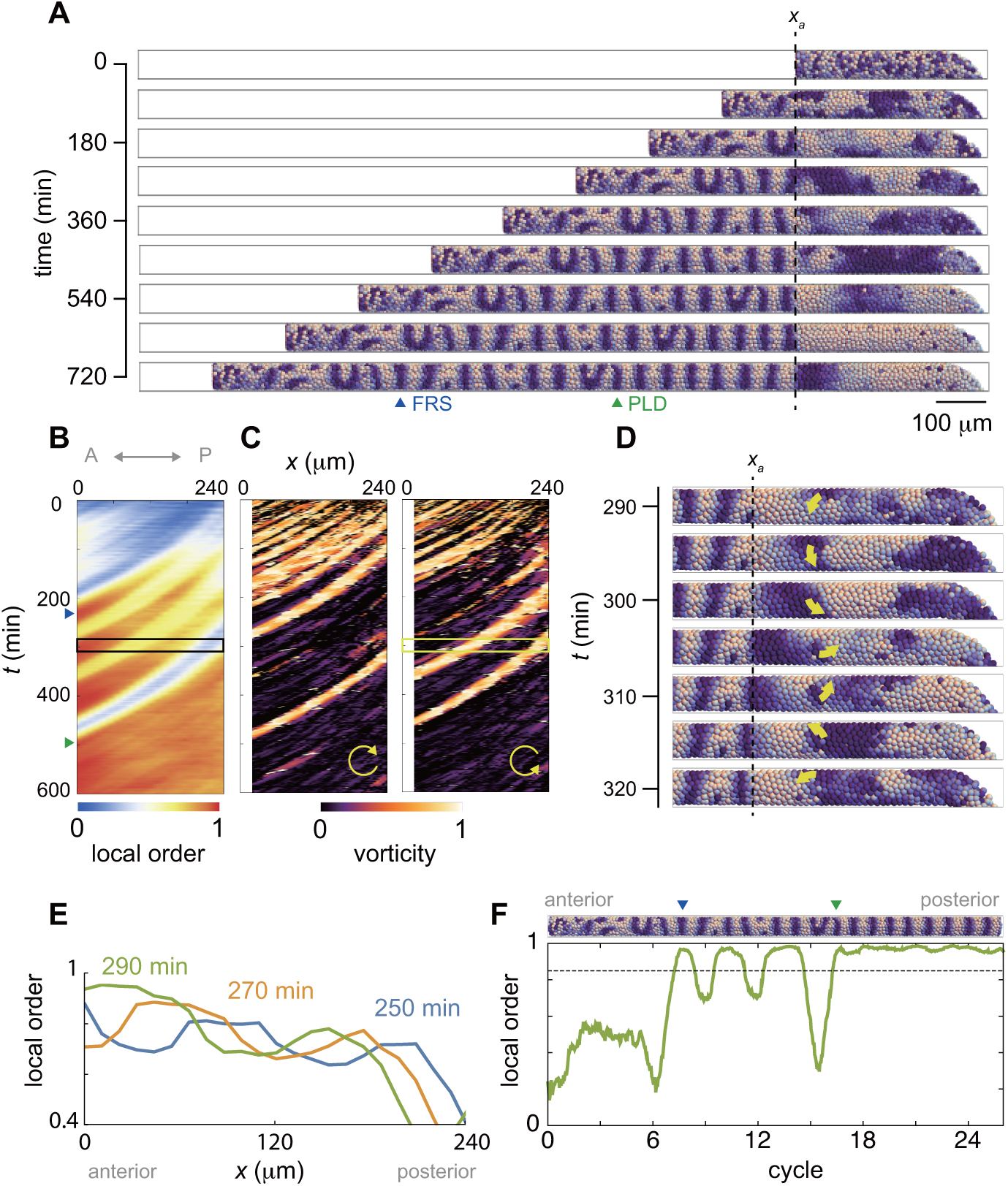
Resynchronization simulations with constant tissue parameters. (A) Snapshots of a resynchronization simulation. Color indicates (1 + sin *θ*_*i*_)/2 where *θ*_*i*_ is the phase of oscillation for cell *i*. The black dotted vertical line indicates the position of the anterior end of the PSM *x*_*a*_ = 0. Tissue parameters are constant over time. See also Movie 2. (B)-(F) Analysis of local phase order and vortex formation in the simulation shown in (A). (B) Kymograph of local phase order parameter of the right PSM shown in (A). (C) Kymographs of phase vorticity for (left) clockwise and (right) counter clockwise rotations. The phase patterns within the black and yellow boxed space-time domains in (B) and (C) are shown in (D). (D) Snapshots of a phase vortex. The yellow arrows indicate the direction of rotation. (E) Local phase order parameters along the anterior-posterior axis of the PSM at different time points. (F) Time series of local phase order parameter at the anterior end of the right PSM *x*_*a*_. The horizontal broken line indicates the threshold *Z*_*c*_ = 0.85 for determining normal and defective segments in simulations. The resultant stripe pattern is on top. In (A), (B) and (F), the blue and green triangles mark FRS and PLD, respectively. Parameter values for simulations are listed in Table S1.

Figure 3A shows snapshots of a synchrony recovery simulation, see also Movie 2. To characterize resynchronization dynamics, we computed local phase order at each position along the anterior-posterior axis, Figs. 3B, S3. Due to local coupling, cells first formed local phase synchronization, Fig. 3A, B and Movie 2. During the first stage of this synchrony recovery, the kymograph of local phase order shows three locally synchronized domains that extended their size due to an increase in number of cells at the same time as they were advected in an anterior direction, Fig. 3B, E. When the size of the most anterior domain with local phase order above a threshold of 0.85 (*Z*_*c*_) exceeded one segment length at its arrival in the anterior end of the PSM, a first recovered segment (FRS) was formed in a simulation, blue triangles in Fig. 3A, B, F. However, because local interactions drove resynchronization in a spatially heterogeneous manner, domains where phase order was lower than *Z*_*c*_ could exist more posterior to such a well-synchronized domain, Fig. 3B, E. Subsequent arrival of the less synchronized domains caused fluctuation of the local order parameter in the anterior end of the PSM, which resulted in defective segment formation after FRS, Fig. 3A, F. Note that patterns of synchronized domains in the left and right sides of the PSM were not well correlated during this time interval, in contrast to the synchronized state. Thus, the numerical simulation suggests that a sequence of synchronized and less synchronized domains along the PSM results in an intermingling of normal and defective segments along the axis. This intermingling matches the experimental observations.

These less-synchronized domains typically formed persistent phase patterns that rotated along an axis as a vortex, as illustrated in the simulation, Fig. 3D and Movie 2. To characterize these patterns, we introduced an index referred to as vorticity, Fig. S4 and Materials and Methods in SI. The kymograph of the vorticity indicates that the less synchronized domains in the kymograph of phase order were caused by the phase vortices, Fig. 3A-C. When a vortex was brought to the anterior end of the PSM through cell advection, it was converted into a defective segment boundary and delayed the PLD, Fig. 3B, C. Thus, the formation of persistent local phase patterns with a mis-orientation to the global pattern in the posterior PSM caused defective segment boundaries after the FRS, providing an explanation for the early washout experiments.

### Dependence of FRS and PLD on each tissue parameter

These results show that the model captures the intermingling of normal and defective boundaries frequently observed in the early washout experiments, but can the model also capture the distribution of FRS and PLD observed in the late washout experiments, thereby joining these observations into a coherent picture of resynchronization across developmental stages?

The finding that transport of local phase patterns across the tissue can influence segment recovery suggests that in addition to local intercellular interactions, tissue level parameters may also be important. Previous experimental studies showed that the PSM length becomes shorter as segments are added (5, 6). Convergent extension by cells in the anterior part of the tissue contributes to advection pattern in the PSM at early developmental stages (38). At later developmental stages, cellular flows from the tissues dorsal to the tailbud change the advection pattern (8-10). These complex rearrangements are represented in our model in a simplified manner by the advection profile. Thus, several lines of experimental data support changes in the PSM length and its advection pattern, and other properties may also vary during development. We therefore studied how the FRS and PLD depend on each of the tissue parameters in the physical model. We begin by shifting a given single parameter to a new constant value, while leaving the others unchanged for the simulation, Table 1 and Figs. S5-S12, and return to the time-dependent cases in the next section. Simulations were started from complete random initial phases as in Fig. 3. We computed FRS and PLD over 100 different realizations of simulations and the results are summarized in Table 1. See Materials and Methods in SI for the quantification of FRS and PLD in simulations.

**Table 1.** Dependence of FRS and PLD on each tissue parameter.

| Influence | Tissue parameters |
| --- | --- |
| change in only PLD | PSM length, cell mixing, advection pattern |
| change in only FRS | none |
| change in both FRS and PLD | coupling strength, PSM radius (tube radius) |
| no or weak effect | frequency profile, tailbud size (torus radius),<br>phase noise intensity |

These results were obtained with simulations where all the tissue parameters were constant over time.

We found that the speed of cell mixing at the tailbud, PSM length *L* and the cell advection pattern in the PSM did not affect FRS, whereas they strongly affected PLD, Figs. S5-S7. Faster cell mixing in the tailbud reduced PLD, Fig. S5, because it prevented the formation of persistent phase vortices at the tailbud. The PSM length and cell advection pattern determined the time taken for a phase vortex to reach the anterior end of the PSM. Cell advection underlies the transport of phase vortices, Figs. 2C, S7. Longer PSM increased PLD because a phase vortex needed more time to arrive at the anterior end by the cell advection, Fig. S6. If cell advection was slower at the posterior part of the PSM, PLD became larger because vortices stayed longer at the posterior part, Fig. S7B. In contrast, if cell advection was faster at the posterior part, PLD became smaller due to faster transport of phase vortices to the anterior part, Fig. S7B. Thus, these parameters are important for understanding the new phenotype, because each can increase the difference between FRS and PLD, producing the intermingled defects observed experimentally.

In contrast, the PSM radius *r* and the coupling strength between neighboring oscillators *κ*_0_ influenced both FRS and PLD, Figs. S8, S9. Smaller PSM radius *r* decreased FRS and PLD, Fig. S8. Smaller cell number at each position in the PSM with a smaller radius *r* allowed local coupling to more rapidly generate a synchronized domain as large as the tissue diameter, leading to a normal segment. Larger values of *κ*_0_ reduced FRS and PLD, Fig. S9. A larger coupling strength reduced the time for local order to form, including vortex patterns. As a result, the last-formed vortex departed the posterior PSM earlier, decreasing PLD.

Coupling keeps phase differences between neighboring oscillators in check. There are two sources of local phase fluctuations in the model: (i) the noise term in individual phase dynamics and (ii) the addition of new cells with random phase values. Desynchronization simulations, where the coupling between cells was absent, demonstrated that the addition of new cells alone was enough to disrupt the wave pattern and compromise the integrity of segment boundaries, Fig. S10. Additionally, a larger phase noise intensity further contributed to a faster decay of the pattern. However, in resynchronization simulations with coupling, both FRS and PLD only weakly depended on the noise intensity, Fig. S10.

Finally, we found that the shape of frequency profile and the torus size for the tailbud *R* did not influence either FRS or PLD, Figs. S11, S12. Note that there was no parameter that influenced only FRS, Table 1. In summary, FRS was determined by parameters that influence local synchronization of oscillators. PLD, on the other hand, was influenced by parameters that control either local synchronization or advection of phase vortices across the PSM.

### Prediction of PLD from DAPT washout timing, PSM shortening and changing advection pattern

In the previous section we considered constant values of parameters defining coupling and tissue properties. However, as noted above, some features like PSM length vary during development on timescales that may be relevant for resynchronization (5-7). To further investigate whether the early and late segmentation phenotypes shown in Fig. 1 could result from a common underlying set of processes, we introduced the washout process into the model, and examined the effect of different washout times in simulations in which tissue properties changed over the course of the simulation.

To model differences in timing of DAPT washout, we started with coupling strength *κ*(*t*) = 0 for *t* < *t*_*wash-out*_ and then switched on coupling at *t* = *t*_*wash-out*_. We assigned random phases to the oscillators in the model as an initial condition, assuming that all DAPT treatments completely desynchronized oscillators, as above. Hence, the phase disordered state lasted until *t* = *t*_*wash-out*_ and resynchronization begun at that time. We performed 100 realizations of simulations for each washout time *t*_*wash-out*_, and recorded the developmental time taken from washout to observation of FRS and PLD, termed the time to FRS (FRS – *t*_*wash-out*_) and time to PLD (PLD – *t*_*wash-out*_).

In the absence of tissue shortening or a changing cell advection pattern, the times to FRS and PLD were not affected by washout time, as expected, Fig. 4A. We analyzed the consequence of PSM shortening on PLD while keeping all the other parameters constant over time, Figs. 4B, S13 and Movie 3. For simplicity, we assumed that the PSM length decreased linearly over time in the simulation, Fig. S13A. PSM shortening decreased the time to PLD for later washout times, Fig. 4B, because in a shorter PSM at later somite stages phase vortices reached the anterior end of the PSM more quickly, Fig. S13B. With higher speed of PSM shortening, the time to PLD after washout became shorter, Fig. S13C-F. As expected from the independence of FRS on constant PSM length, PSM shortening did not affect FRS, Figs. 4B, S6.

**Figure 4.**
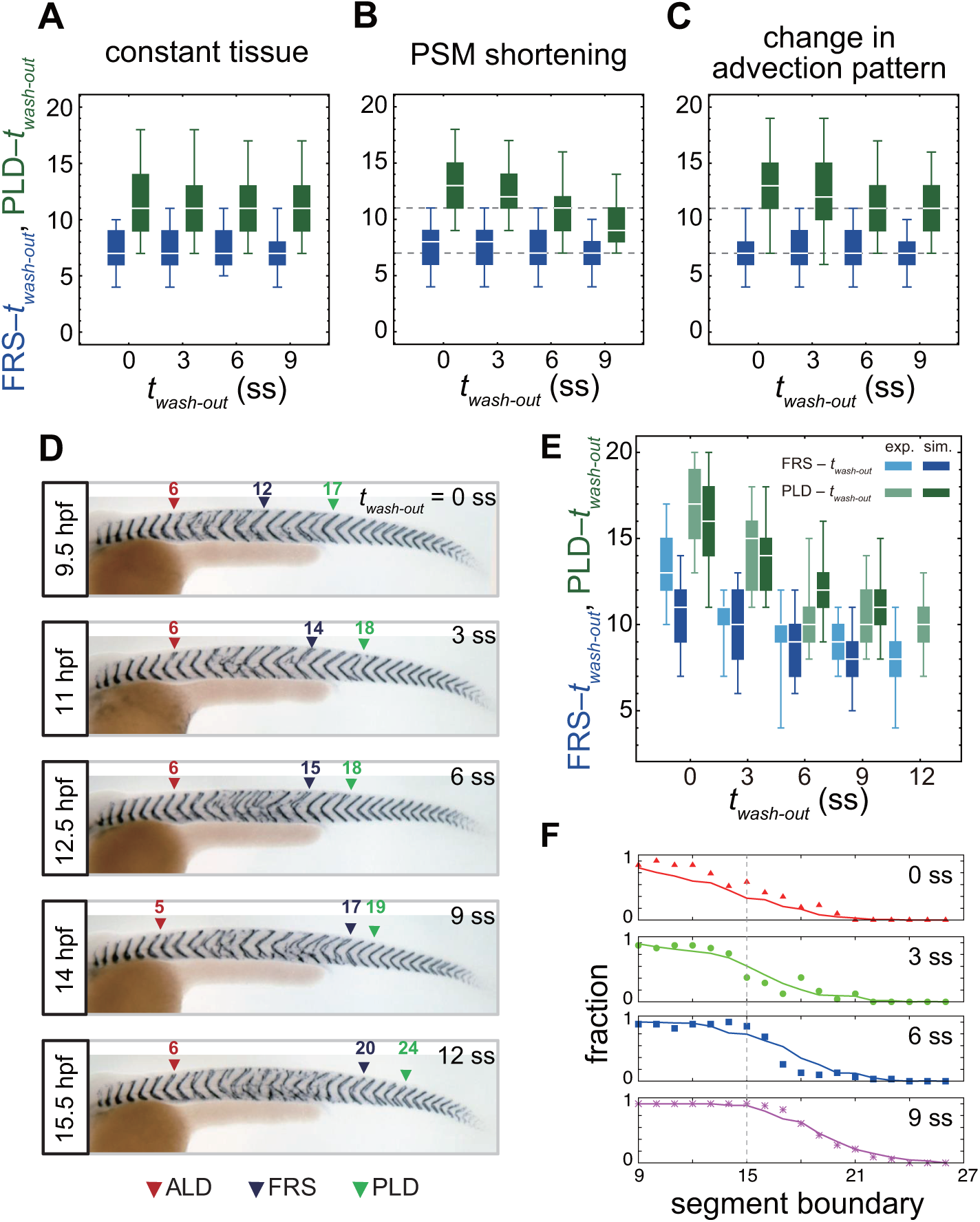
Gradual transition from early to late washout boundary phenotypes captured by the physical model. (A)-(C) Dependence of times to FRS and PLD on DAPT washout time for different conditions in simulations. (A) Constant tissue where all the tissue parameters remain unchanged during a simulation. (B) PSM length becomes shorter with time. All the other parameters are constant. See also Movie 3. (C) Cell advection pattern changes at 9 somite stage (ss). Before 9 ss, the strain rate is larger in the anterior than posterior PSM. After 9 ss, the strain rate becomes larger in the posterior PSM. See also Movie 4. All the other parameters are constant. The box-whisker plots indicate 5, 25, 75 and 95 percentiles. The white bars mark the median. In (B) and (C), the gray dotted lines mark the medians of FRS and PLD in the constant tissue shown in (A). (D) Whole-mount *in situ* hybridization for the myotome segment boundary marker gene *xirp2a* in ∼36 hours post fertilization (hpf) embryos. DAPT washout time is 9.5 hpf (0 ss; *n* = 28), 11 hpf (3 ss; *n* = 22), 12.5 hpf (6 ss; *n* = 28), 14 hpf (9 ss; *n* = 30) and 15.5 hpf (12 ss; *n* = 26) from top to bottom. Red, blue and green triangles indicate the ALD, FRS and PLD, respectively. (E) Dependence of times to FRS and PLD on DAPT washout time. Light blue and green box-whisker plots indicate 5, 25, 75 and 95 percentiles for embryonic experimental data (exp.). Dark blue and green box-whisker plots indicate those for simulation data (sim.). The white bars mark the median. The PSM shortening, change in cell advection pattern and increase in the coupling strength are combined in the model, see also Movies 5, 6. The lack of information about the formation of final segments in embryos precludes simulations for the latest washout (12 ss), see the text. (F) Spatial distribution of segment boundary defects. Symbols indicate embryonic experimental data and lines indicate simulation data. Grey dashed vertical line across panels is a guide to the eye. In (A)-(C), (E), results of 100 realizations of simulations with each washout timing are plotted. Parameter values for numerical simulations are listed in Tables S1 and S2.

We next analyzed the effect of a change in cell advection pattern on PLD, keeping all the other parameters constant over time, Figs. 4C, S14 and Movie 4. In the model we represented the change in the advection pattern in a simplified way such that at earlier somite stages (*t* < *t*_*g*_) the local strain rate was larger in the anterior region of the PSM, whereas the strain rate became larger in the posterior region at later stages (*t* > *t*_*g*_), Fig. S14A. We found that such a change in advection pattern increased time to PLD for earlier DAPT washout, Fig. 4C. If the change in advection pattern occurred at later developmental stages, time to PLD for later washouts were also increased, Fig. S14D-G. As described above, the cell advection pattern in the PSM underlies the transport of phase vortices. When advection did not occur in the posterior PSM at earlier somite-stages, the movement of phase vortices relative to the tailbud was slowed in that region, delaying PLD for earlier washout timing, Fig. S14A-C. As expected from the independence of FRS on constant advection patterns, a change in pattern did not affect FRS, Figs. 4C, S14.

Taken together, these results predict that the changes in tissue length and cell advection pattern observed in the embryo over developmental time may have an impact on resynchronization dynamics, reflecting in the decrease in difference between FRS and PLD. Thus, changing tissue properties may explain the difference between early and late washout phenotypes.

### Embryonic segment recovery depends on timing of DAPT washout

To test the theoretical predictions about the influence of tissue-level changes on the time from washout to FRS and PLD over developmental stages, we next performed DAPT washout experiments, as illustrated in Fig. 4D, in which DAPT was removed at different times (*t*_*wash-out*_) between 9.5 and 15.5 hpf. We visualized the resulting distribution of segment defects along the axis on both left and right-hand sides of the embryo and identified the FRS and PLD, arrowheads in Fig. 4D.

We found that FRS and PLD both increased with later washout times, Fig. S15. In accordance with the prediction of the model, the time to PLD (PLD – *t*_*wash-out*_) decreased gradually over developmental time, Fig. 4E. In contrast, the experimentally observed gradual decrease in time to FRS (FRS – *t*_*wash-out*_) in Fig. 4E was not expected from the simulations, Fig. 4B, C. After washout, it took ∼ 13 segments to observe FRS for an earlier washout time, whereas it took 8 segments for a later washout time. Embryos yielded more scattered, non-continuous defects with earlier than with later washouts, Figs. 4D, S1C-G.

Combined, these results revealed the transition between early and late washout segmentation phenotypes. The observed decrease in the time to PLD was qualitatively consistent with the theoretical predictions. However, the expectation that FRS would be independent of washout timing, based on its insensitivity to global tissue properties of PSM shortening and changing cell advection patterns in the model, was not observed experimentally. We therefore hypothesized that FRS might instead be affected by mechanisms that determine the level of local synchronization.

### Prediction of FRS from DAPT washout timing and increasing coupling strength

Local synchronization is thought to be driven by local intercellular interactions. The intensity of such local interactions is described in the theory by the coupling strength *κ*_0_ between neighboring oscillators. As discussed previously, coupling strength can strongly influence FRS, Table 1 and Fig. S9. Although FRS can also be influenced by the PSM radius, which becomes smaller with developmental stage, the effect of change in the PSM radius was weaker than the coupling strength within the biologically plausible range, Figs. S8, S16. Therefore, we tested whether a changing coupling strength could describe the dependence of time to FRS on *t*_*wash-out*_ observed in experiments.

For simplicity we assumed that, in the absence of any perturbation, the coupling strength increased as a linear function of time in the simulation, Fig. S17A. An increase in coupling strength over developmental stages in the embryo could be caused by an increase in the abundance or activity of Delta and Notch proteins in cells (21, 26, 39, 40), an increase in the contact surface area between neighboring cells, or some other mechanism.

A temporal increase in coupling strength in the simulation, allowing it to double by ∼15 ss, led to a decrease in the time to FRS that reproduced the experimental results, Fig. S17B. The time to PLD also decreased with *t*_*wash-out*_. However, the magnitude of reduction in time to PLD was similar to that in time to FRS, meaning that this effect would not be expected to contribute to the observed experimental reduction in the difference between PLD and FRS. These results indicate that the increase in the coupling strength over somite stages alone is sufficient to realize the dependence of time to FRS on *t*_*wash-out*_, whereas the global tissue parameters contribute to the behavior of PLD observed in experiments.

### Embryonic segmentation defect patterns are captured by PSM shortening, change in cell advection pattern and an increase in coupling strength

Finally, we simulated a resynchronizing PSM with all the three effects described above combined: the changes in PSM length and cell advection pattern over time were imposed by existing experimental data, the change in coupling strength was motivated by the results in the previous section, and all other parameters remained unchanged, Figs. 4E, S18 and Movies 5, 6.

The beginning and the end of somitogenesis are special. The segmentation clock becomes active and rhythmic before somitogenesis starts, during epiboly (19). At this stage the embryo undergoes dramatic morphological changes that we do not describe with the current model, which instead describes the PSM shape from 0 ss onwards. For the end of somitogenesis, there is a lack of quantitative information about the formation of the final segments that precludes constraining the theory at this late stage. Therefore, we simulated DAPT washout times from the beginning of somitogenesis 0 ss (*t*_*wash-out*_ = 9.5 hpf) until 9 ss (*t*_*wash-out*_ = 14 hpf), where the model could be well parametrized and provided a fair description of tissue shape changes.

The requirement for many realizations to compute FRS and PLD precluded application of standard fitting procedures for determining parameter values in the model. Instead, we used parameter values close to those observed in embryos. We found a decrease in time to PLD with the magnitude of the decrease greater than time to FRS, as observed in the experimental data, Fig. 4E. Inclusion of the PSM shortening and change in cell advection pattern in simulations recapitulated the experimental observation that the difference between PLD and FRS became smaller with later washout time. PSM shortening decreased time to PLD, without affecting FRS, thereby reducing the difference between PLD and FRS for later washout time, Fig. 4B. The slower cell advection in the posterior PSM at earlier time in simulations delayed PLD without affecting FRS, in principle enlarging the difference between PLD and FRS for earlier washout time, Fig. 4C.

In summary, a change in the coupling strength was sufficient to reproduce the behavior of FRS. Combined effects of the PSM shortening and cell advection pattern were the dominant factors that generated the behaviors of PLD in simulations. Thus, the physical model quantitatively reproduced the behaviors of FRS and PLD, suggesting that the physiologically plausible changes in these tissue parameters may underlie behaviors observed in the experiment.

### Prediction of segment defect distribution

We showed that the model could capture the onset of segment boundary recovery and its completion, quantified by FRS and PLD, respectively. However, segment recovery is a complex gradual process reflected in intermingled segment defects. Therefore, we further tested whether the model captured this gradual recovery process between its onset and completion with data that were not used to develop it: the spatial distribution of segment boundary defects along the embryonic body axis.

We used the same parameters listed in Table S2 that we established to describe FRS and PLD, Fig. 4E. Since simulations started from completely random initial phases, the initial fraction of defective segments was one. The fraction of defective segments decreased from one to zero along the body axis after DAPT washout, shifting posteriorly for later *t*_*wash-out*_, Fig. 4F. We then compared the simulated axial distribution of defective segments with embryonic DAPT washout experiments. We restricted the comparison to the washout phase of the experiment. We counted the number of defective segments along the axis in embryos and defined the fraction of defective boundaries at each segment position, Fig. 4F. The distributions of defective segments were similar between left and right sides of embryos, Fig. S19A. After washout, the fraction of defective segment boundaries gradually decreased, and eventually it became zero, suggesting that synchronization was fully recovered at that time. As DAPT was washed out at increasingly later times, defective segment boundaries continued to more posterior locations, in agreement with simulations, Fig. 4F.

From this distribution, we could compute ALD, FRS and PLD using probability theory, see Fig. S19 and SI. This distribution also explained the ratio of single defects, where either the left or right segment was defective at a segment locus, to double defects, where both left and right segments were defective, Fig. S20. This agreement between experimental data and probability theory for the fraction of single defects suggests that recovery occurred independently between left and right PSMs. In summary, the physical model predicted the segment defect distribution, providing a thorough description of synchrony recovery.

## DISCUSSION

The segmentation clock produces dynamic patterns that determine the formation of vertebrate segments. This multicellular clock, consisting of thousands of cells that make the PSM and tailbud, produces a kinematic wave pattern. The integrity of this wave pattern relies on local synchronization of the oscillators mediated by Delta-Notch cell-cell signaling. Our current view of synchrony is largely informed by desynchronization experiments in which cells were uncoupled by interfering with Delta-Notch signaling, resulting in a loss of wave patterns. In contrast, resynchronization, where oscillators re-establish coherent rhythms from a desynchronized state, can be used to probe how tissue-scale collective patterns arise from local interactions during morphogenesis.

In this study, we applied a combination of experiments and theory to explore the recovery of normal body segments during the resynchronization of oscillators. Our surprising experimental discovery was regions of normal and defective segments intermingled along the body axis following DAPT washout. Since we seek to capture pattern recovery at multiple scales, our new physical model describes cellular oscillations with an effective phase variable together with local intercellular interactions (28, 34), as well as larger scale mechanics such as cell movements (28, 41) and tissue shape changes (7). The model qualitatively explains the formation of these intermingled defective segments by the emergence of persistent phase vortices in the posterior PSM and their advection through the tissue to the anterior. The vortices arise from the local coupling of desynchronized oscillators and the advection is a consequence of axis elongation. As vortices arrive at the anterior end, their mis-oriented local phase patterns result in segment defects, before global pattern recovery is achieved. Importantly, our model achieves a quantitative description of the experimental phenotype by additionally incorporating global features of the PSM and tailbud that are observed in the embryo, such as tissue length change, a change in the advection pattern and a gradient of cell mixing. Further, we introduced observables such as an index of off-lattice vorticity, and a local order parameter to distinguish normal and defective segments for comparison between simulations and experiments. Simulations of the model confirm that stronger local coupling leads to faster resynchronization and recovery of the gene expression pattern, as expected from previous work (26). The time-dependent increase in effective coupling strength used in the simulation is plausible from existing data of Delta and Notch gene expression, for example, but remains an expectation of the current work. Thus, this formulation has allowed a quantitative comparison to experimental data and a phenomenological understanding of pattern recovery.

Vortices could arise in other models of the segmentation clock with similar local interactions (28, 34, 41, 42). Indeed, vortex formation is a common feature of systems that can be described with locally coupled polar variables, in both excitable and oscillatory media (43). These include biological systems such as cAMP patterns in aggregating populations of *Dictyostelium* cells (44) and spiral patterns in heart tissue (45), planar cell polarity (46), chemical systems like the BZ reaction (45, 47, 48) and in physical systems generally (49, 50). Further characterization of the vortex dynamics in our model remains an interesting topic for future work.

Recent experiments using cells isolated from mouse tailbuds and reaggregated to form so-called emergent PSM have shown the emergence of striking wave patterns that depend on Notch signaling, suggesting that local interactions can drive large-scale dynamical features (51, 52). The relationship between such features and normal versus defective segment boundary formation remains to be explored. Since phase vortices arise naturally in systems of locally coupled oscillators starting from disordered initial conditions, Movie 7, we predict that these structures will form also in mammalian PSM tissue culture systems (51-54). The framework of our model with different geometries will facilitate analysis of dynamics in these and other collective cellular systems with both local interactions and tissue-level deformations.

In simulations, the kinematics of phase vortices across the PSM is determined by global tissue properties, including the PSM length and its cell advection pattern. In this way, the recovery of a gene expression wave pattern across the PSM and subsequent normal boundary formation is set both by the timescales of local synchronization through intercellular interactions, and also by those of morphological processes. In addition, this result implies that the quantification of global tissue parameters will be necessary to obtain quantitative agreement between theory and experiment. Temporal changes in PSM length have been measured in various species (5, 6, 55), and cell advection patterns across the PSM have been investigated (8-10, 56). Involvement of these global tissue properties discriminates the resynchronization of the segmentation clock from its desynchronization, which is dominated by local cellular properties such as noise in clock gene expression (18, 19, 23, 57). Furthermore, morphological tissue development distinguishes the synchronization of the segmentation clock from that of other biological oscillator systems in spatially static tissues, such as the suprachiasmatic nucleus in the mammalian circadian clock (58).

Numerical simulations of the physical model showed that the first recovered segment FRS and the posterior limit of defects PLD contain different information about resynchronization. From FRS, we estimated properties of local synchronization, such as the coupling strength between genetic oscillators. In contrast, PLD is influenced by global tissue properties, such as tissue length and cell advection pattern, that determine the transport of persistent mis-oriented phase patterns. Hence, it is important to choose these measures appropriately depending on the question addressed. For example, recent studies suggested that the level of cell mixing observed in the tailbud promotes synchronization of genetic oscillation (28). This effect appears in PLD, but not in FRS, because the elevated mixing in the tailbud prevents formation of the local phase patterns at the posterior PSM. On the other hand, FRS can be a better measure for the coupling strength (26), because it is less affected by the other tissue parameters than PLD.

In conclusion, our study of resynchronization has revealed how the segmentation clock can be influenced by global developmental processes such as tissue length change, cell advection pattern and cell movement, as well as local coupling strength between cells. We propose that the distinctive intermingling of well-formed and defective segments seen during recovery arises from persistent phase vortices. Such transient patterns occurring during the resynchronization of biological oscillators would be difficult to recognize in snapshots of the segmentation clock due to the restricted system geometry, the discrete character of cellular tissue, and the difficulty of determining phase from a single time point, and consequently may have been overlooked in previous experiments. However, they should soon be within reach of techniques for perturbation of cell coupling combined with live imaging over the long durations required to observe them in the embryo (59, 60). Our findings suggest that segmental pattern recovery occurs at two scales: local pattern formation and transport of these patterns through tissue deformation. Other developing systems such as *Dictyostelium* colony aggregation (44), and tissue elongation in the *Drosophila* pupal wing (61) also feature an interplay between locally driven interactions and global morphological changes, pointing to a common principle of pattern dynamics within developing tissues.

## ACKNOWLEDGEMENTS

We thank the fish facility staff of the MPI-CBG, Dresden and Arianne Bercowsky, Gabriela Petrungaro, Sundar Naganathan, and Olivier Venzin for comments on the manuscript. Research was sponsored by JSPS KAKENHI grant number 17H05762, 19H04955, 19H04772 to KU; Ministry of Science and Technology, Taiwan (MOST 108-2311-B-019-001-MY3) to B-KL; SNSF Project funding division III (31003A_176037) and Wellcome Trust Senior Research Fellowship in Basic Biomedical Science (WT098025MA) to AO; European Research Council Starting Independent Research Grant (ERC-2007-StG: 207634) and the Francis Crick Institute to B-KL and AO; ANPCyT PICT 2012 1954, PICT 2013 1301, PICT 2017 3753 to LGM, FOCEM-Mercosur (COF 03/11) to IBioBA; and JSPS Short Term Grant S17064 to KU and LGM.

---

**Figure S1.**
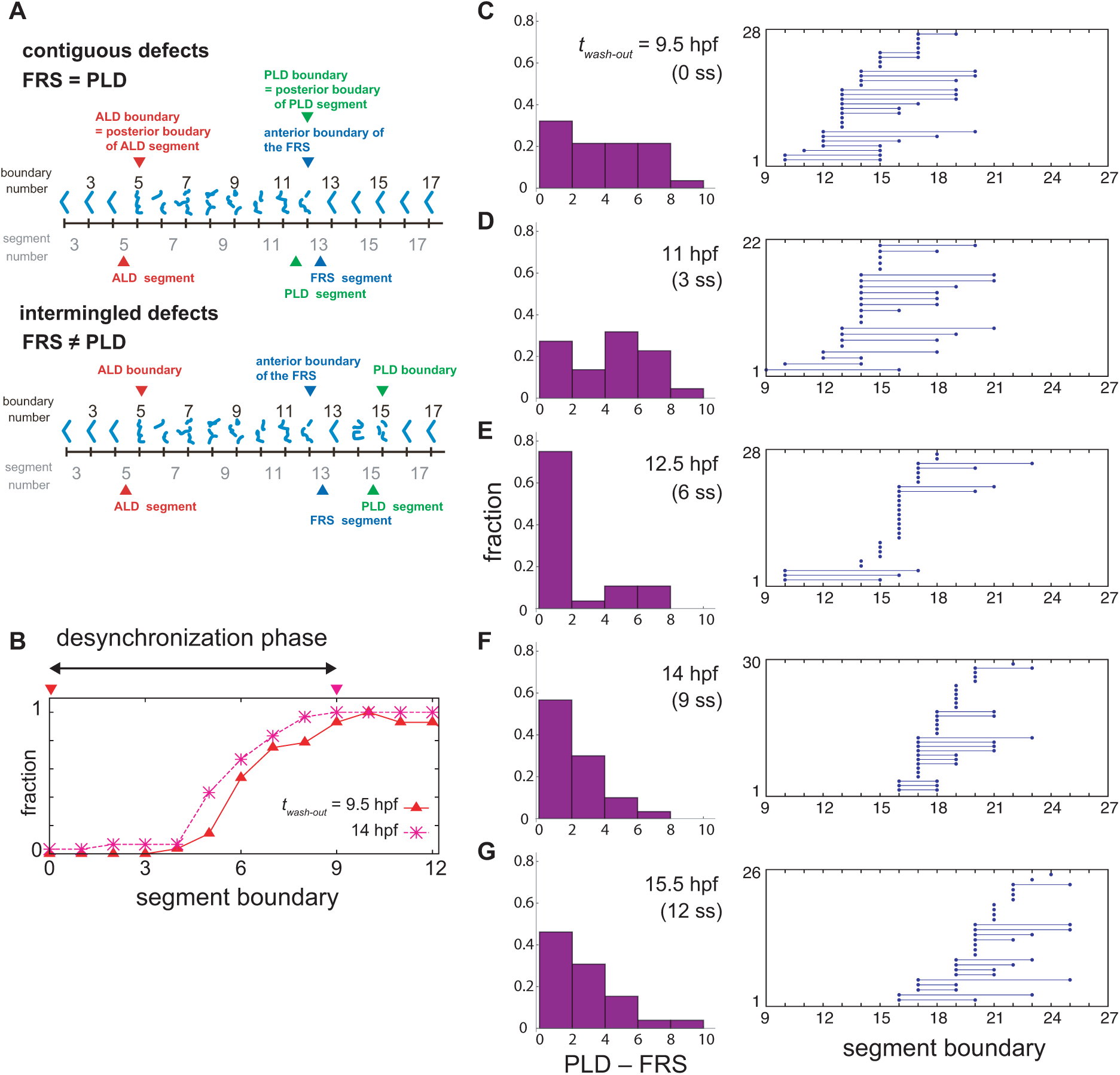
Difference between FRS and PLD in experimental data. (A) Segment defects and definitions of the anterior limit of defects (ALD), first recovered segment (FRS) and posterior limit of defects (PLD). Blue chevrons indicate segment boundaries. Fragmented or broken chevrons represent defective boundaries. Only one embryonic side is illustrated. In this study, embryos were scored one side at a time. Distributions of defective segments may be different between left and right sides of embryos. ALD and PLD is defined with the posterior boundary of the first and last defective segments, respectively. FRS is defined with the anterior boundary of the first normal segment. (B) Spatial distribution of defective segment boundaries for DAPT washout at 9.5 hours post fertilization (hpf) (0 somite-stage, ss: triangles, *n*=28) and at 14 hpf (9 ss: crosses, *n*=30). The horizontal axis is segment boundary number. The vertical axis is the fraction of defective segment boundaries over embryos. Because of the increase in the fraction of defective segments, the desynchronization phase is considered to continue until segment 9. The two inverted triangles indicate DAPT washout times. (C)-(G) Left: Histograms of the difference between FRS and PLD (PLD – FRS). When the first normal segment boundary is found immediately after the last defective boundary, FRS and PLD coincide, and PLD – FRS is zero, see (A) and Materials and Methods. Right: FRS and PLD for each embryonic left-right side. The left circles indicate FRS. The right circles connected with the left circles by lines indicate PLD of the same embryonic side. Single circles indicate that FRS and PLD are the same. The number of examined embryonic left-right sides is shown in the vertical axis. DAPT washout at (C) 9.5 hpf, (D) 11 hpf, (E) 12.5 hpf, (F) 14 hpf, and (G) 15.5 hpf.

**Figure S2.**
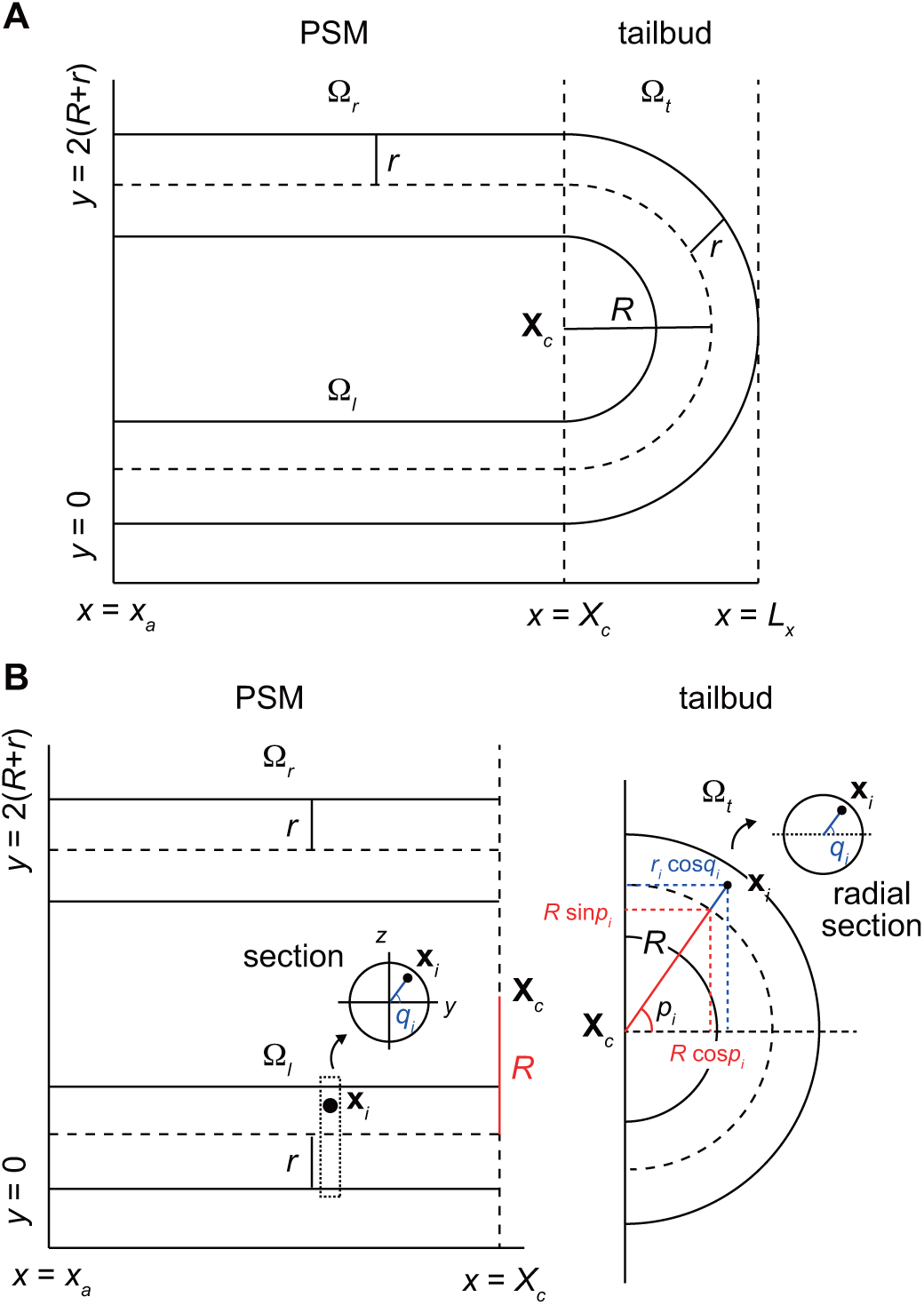
3D geometry of the PSM and tailbud in the physical model. (A) Two tubes and a half torus represent the PSM and tailbud, respectively with anterior to the left and posterior to the right. The *z*-axis is perpendicular to the paper. (B) Position of cell *i* can be expressed with a radial distance *r*_*i*_ from the center of core curve of torus or that of tubes, and two angles *p*_*i*_ and *q*_*i*_.

**Figure S3.**
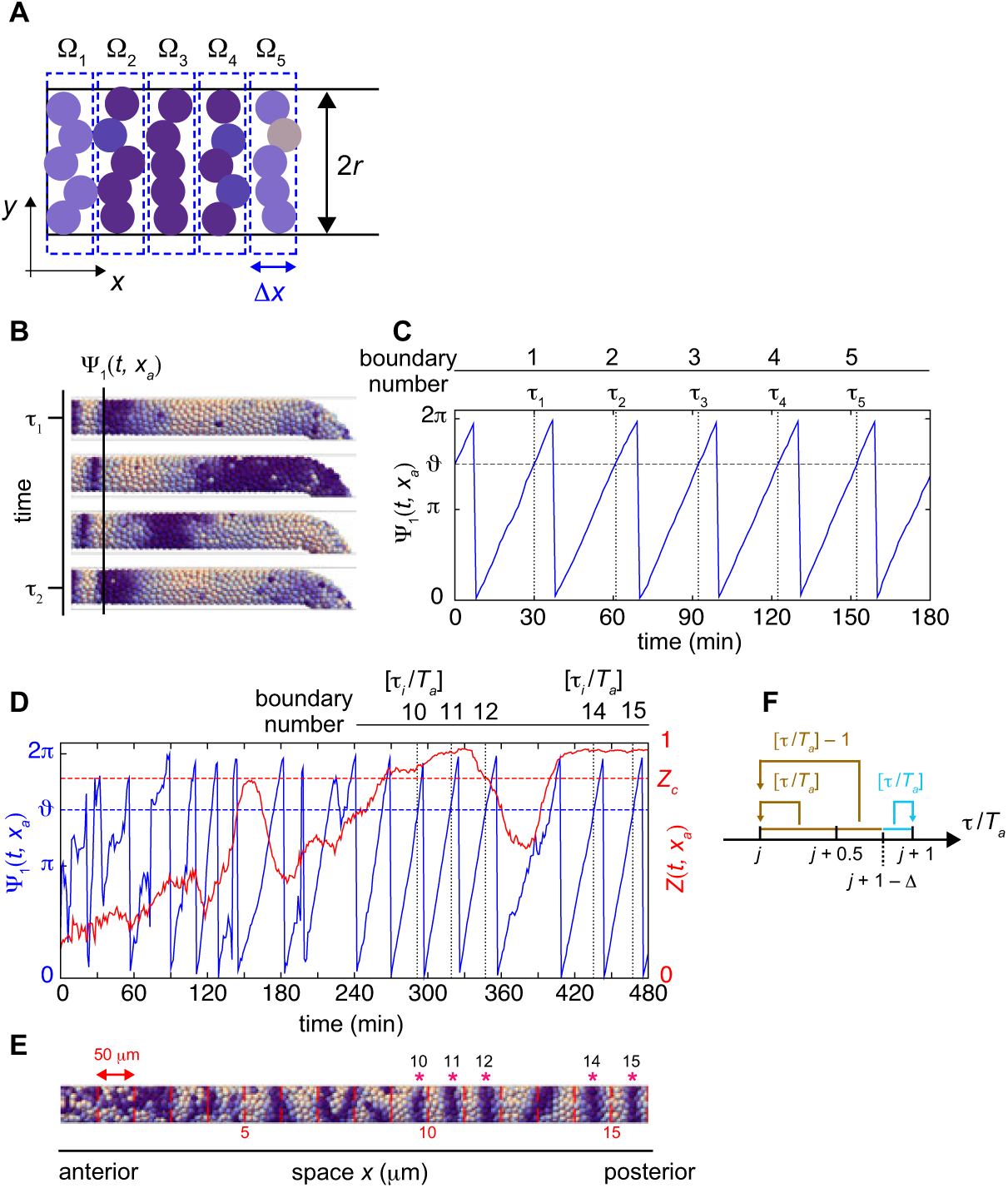
Definitions of local phase order and a normal segment boundary in simulations. (A) Calculation of local phase order. Kuramoto phase order parameter *Z*_*m*_ in each domain Ω_*m*_ (*m* = 1, 2, .5) are first calculated. The average over these domains is defined as a local phase order. Colored circles indicate cells. The width of the domain Δ*x* is equal to the cell diameter *d*_*c*_, Δ*x* = *d*_*c*_ and 5Δ*x* is approximately one segment size. (B) Detection of a segment boundary position by the mean phase at the position *x*_*a*_. (C) Time series of the mean phase at the position *x*_*a*_. When the value of mean phase becomes *ϑ*, a segment boundary is considered to be formed. *τ*_*i*_ indicates times at which segment boundaries are formed. In (B) and (C), a simulation was started from a synchronized initial condition for illustration. (D) Time series of the mean phase (blue) and local phase order (red) at the position *x*_*a*_. The red dotted line indicates the threshold for the local phase order to determine normal boundaries. *τ*_*i*_ indicates time when the value of the mean phase surpasses *ϑ*. *T*_*a*_ is the period of oscillation at the position *x*_*a*_. (E) Formed segment boundaries. The asterisks indicate normal segment boundaries. The red vertical lines indicate expected segment boundary positions. (F) Assignment of segment boundary number. See Materials and Methods for details.

**Figure S4.**
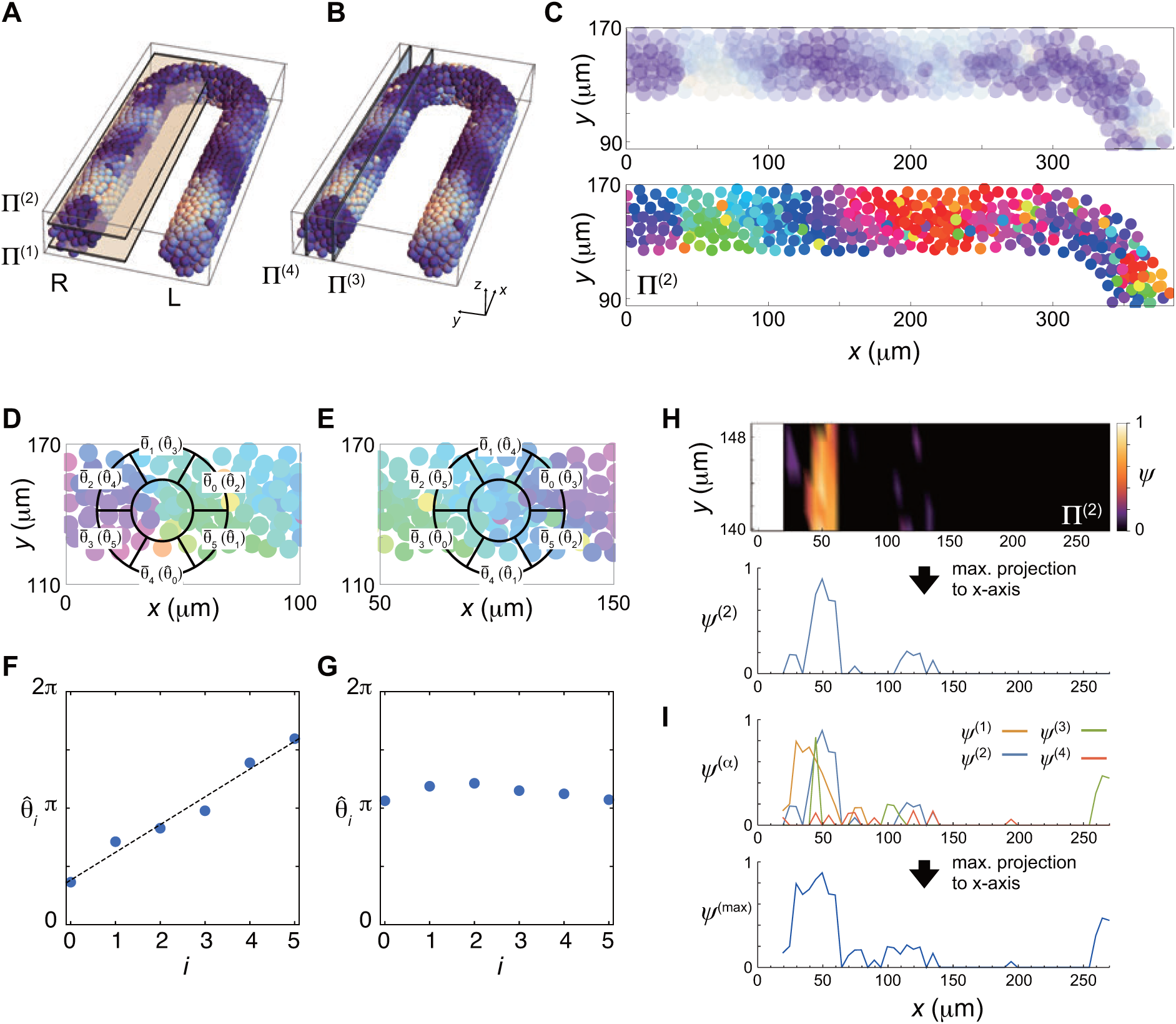
Calculation of vorticity. (A), (B) Planes Π^(*i*)^ for vorticity calculation. (C) Phase distribution in Π^(2)^ of the right PSM. The values of (top) (1 + sin *θ*_*i*_)/2 and (bottom) *θ*_*i*_ are color coded. (D), (E) Average phase 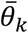 of the subdomain *V*_*k*_ of a ring located at a lattice point. 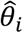 is the permutation of 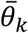. A phase vortex is present in the region shown in (D), whereas there is no vortex in (E). (F) Linear increase of the values of 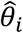 along the perimeter of the ring shown in (D). The black line indicates a linear fit to the data points. The vorticity is defined as 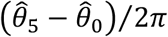. (G) If there is no phase vortex as shown in (E), 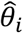 does not increase linearly. (H) (top) Spatial distribution of the vorticity in Π^(2)^. (bottom) The vorticity is projected to *x*-axis as *ψ*^(2)^. (I) Maximum projection of *ψ*^(*i*)^ obtained in each plain Π^(*i*)^ to *x*-axis.

**Figure S5.**
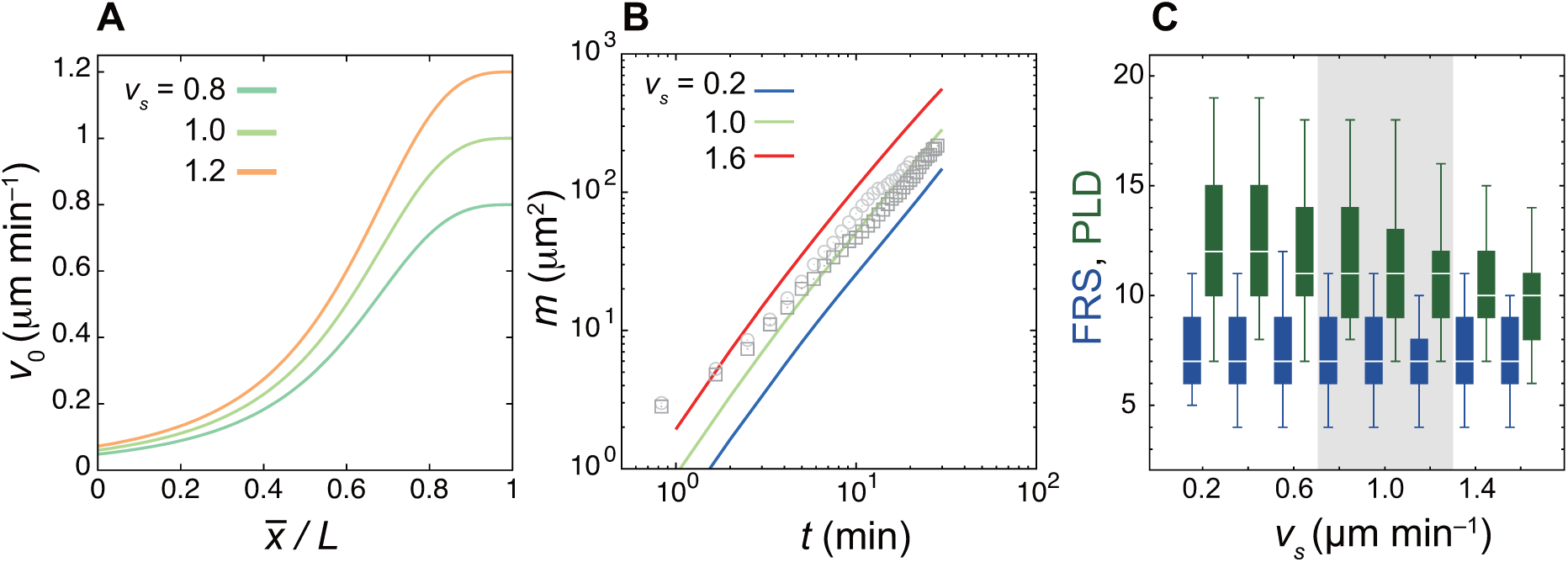
Faster cell mixing reduces PLD whereas it does not influence FRS. (A) Shapes of the speed of intrinsic mobility gradient *v*_0_(*x*) in Eq. (3) in the Materials and Methods for different values of the maximum speed at the tailbud *v*_*s*_. The origin of the horizontal axis is the anterior end of the PSM 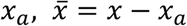. (B) Time evolution of the mean squared difference of displacement vector (MSDD) for different values of *v*_*s*_. The grey circles and squares indicate the MSDDs of two 17 somite-stage embryos quantified previously (Uriu et al. 2017, ref. 28 in the main text) as references. (C) Dependence of FRS (blue) and PLD (green) on *v*_*s*_. The box-whisker plot indicates (0.05, 0.25, 0.75, 0.95) quantiles of FRS and PLD for 100 realizations of simulations. The white bars indicate median. The shaded interval of *v*_*s*_ is the range within which simulated MSDD is close to those from embryonic tissues shown in (B).

**Figure S6.**
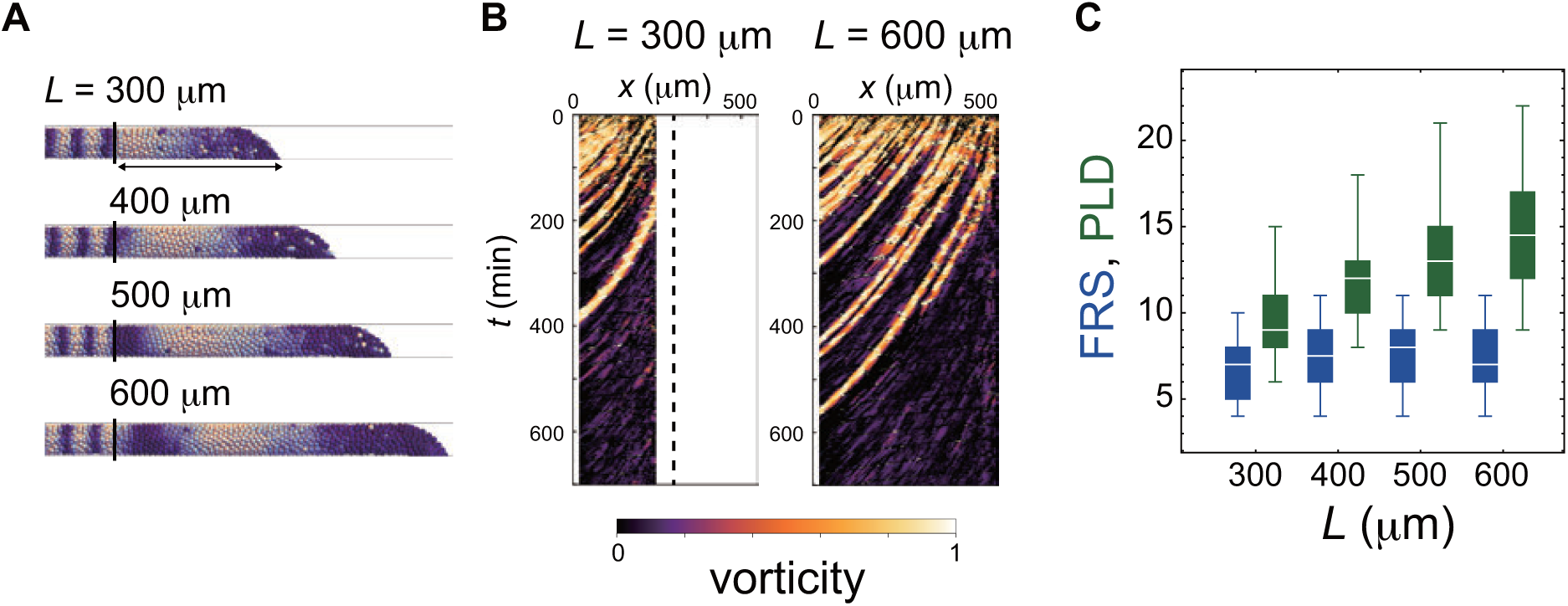
Dependence of FRS and PLD on the PSM length in simulations. (A) PSMs with different lengths. (B) Kymographs of vorticity for the different PSM lengths (left: 300 μm and right: 600 μm). Vorticity of the right PSM is shown. Trajectories of counter-clockwise vortices for PSM length *L* = 300 μm and *L* = 600 μm are presented. (C) Dependence of FRS (blue) and PLD (green) on the PSM length *L*. The box-whisker plot indicates (0.05, 0.25, 0.75, 0.95) quantiles of FRS and PLD for 100 realizations of simulations. The white bars indicate median.

**Figure S7.**
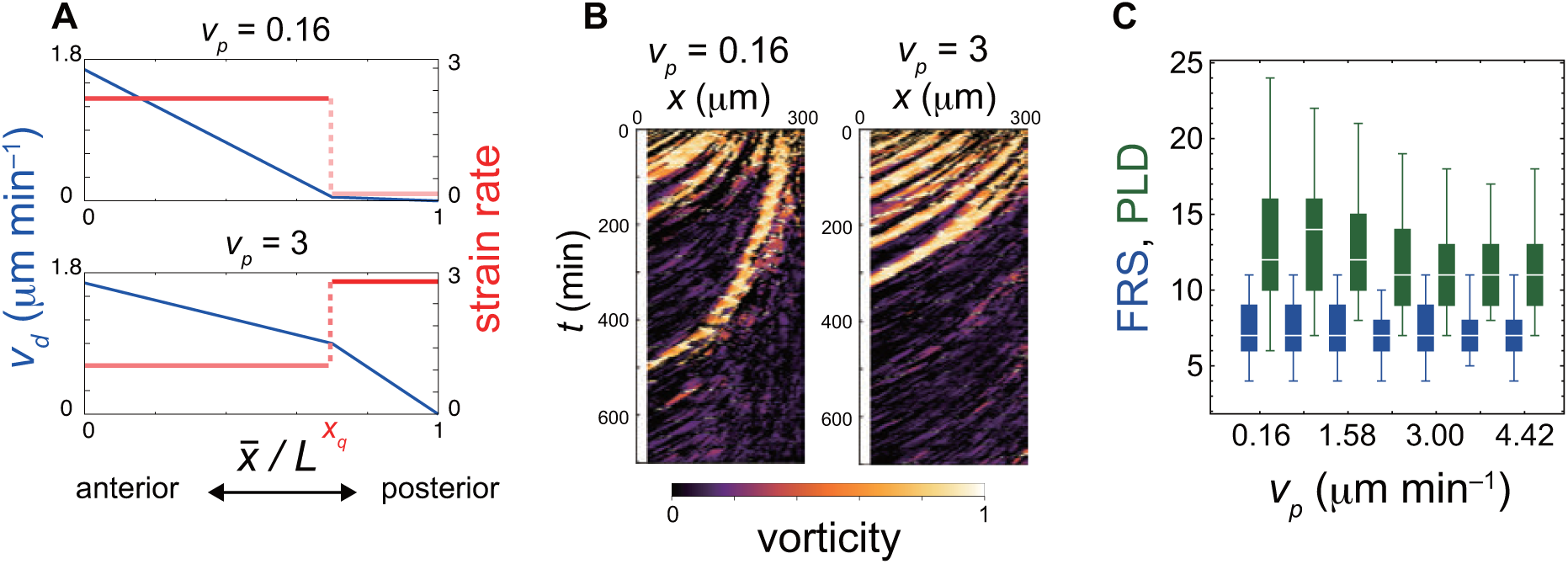
Dependence of FRS and PLD on PSM advection pattern in simulations. (A) Advection speed *v*_*d*_ (blue) as a function of normalized position, 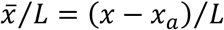. *v*_*p*_ is the strain rate that is the magnitude of spatial derivative of advection speed (red) at the posterior part of the PSM 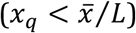. A smaller *v*_*p*_ (top: *v*_*p*_ = 0.16) represents a lower strain rate at the posterior part than the anterior part of the PSM. A larger *v*_*p*_ (bottom: *v*_*p*_ = 3) represents a larger strain rate in the posterior part of the PSM. (B) Kymographs of vorticity for (left) *v*_*p*_ = 0.16 and (right) *v*_*p*_ = 3. Vorticity across the right PSM is shown. Trajectories of counter-clockwise vortices are presented. (C) Dependence of FRS (blue) and PLD (green) on *v*_*p*_. The box-whisker plot indicates (0.05, 0.25, 0.75, 0.95) quantiles of FRS and PLD for 100 realizations of simulations. The white bars indicate median.

**Figure S8.**
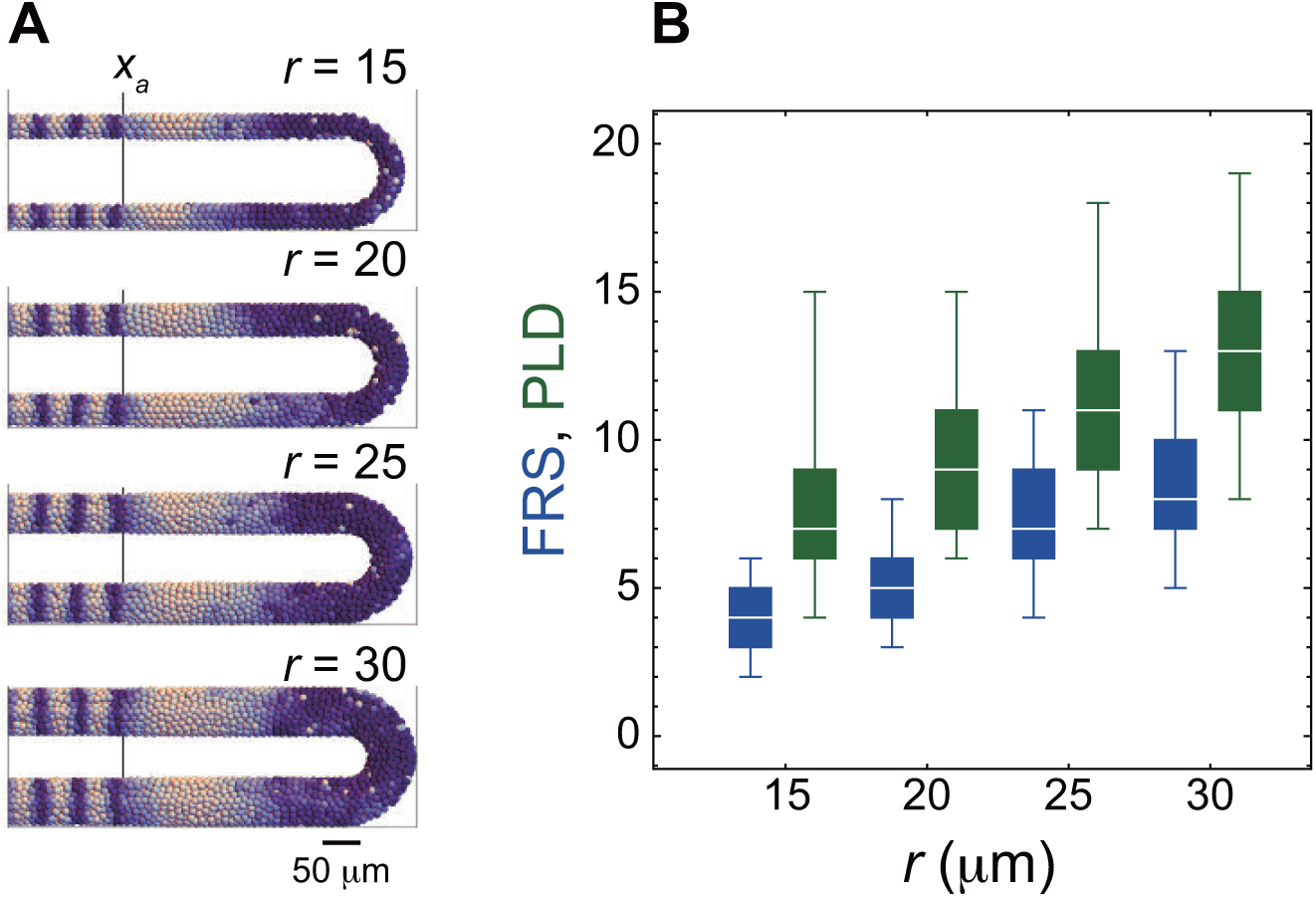
PSM radius *r* influences both FRS and PLD. (A) Snap shots of the spatial phase patterns in the PSM with different values of *r*. The black vertical lines indicate the anterior end of the PSM *x*_*a*_. (B) Dependence of FRS (blue) and PLD (green) on *r*. The box-whisker plot indicates (0.05, 0.25, 0.75, 0.95) quantiles of FRS and PLD for 100 realizations of simulations. The white bars indicate median.

**Figure S9.**
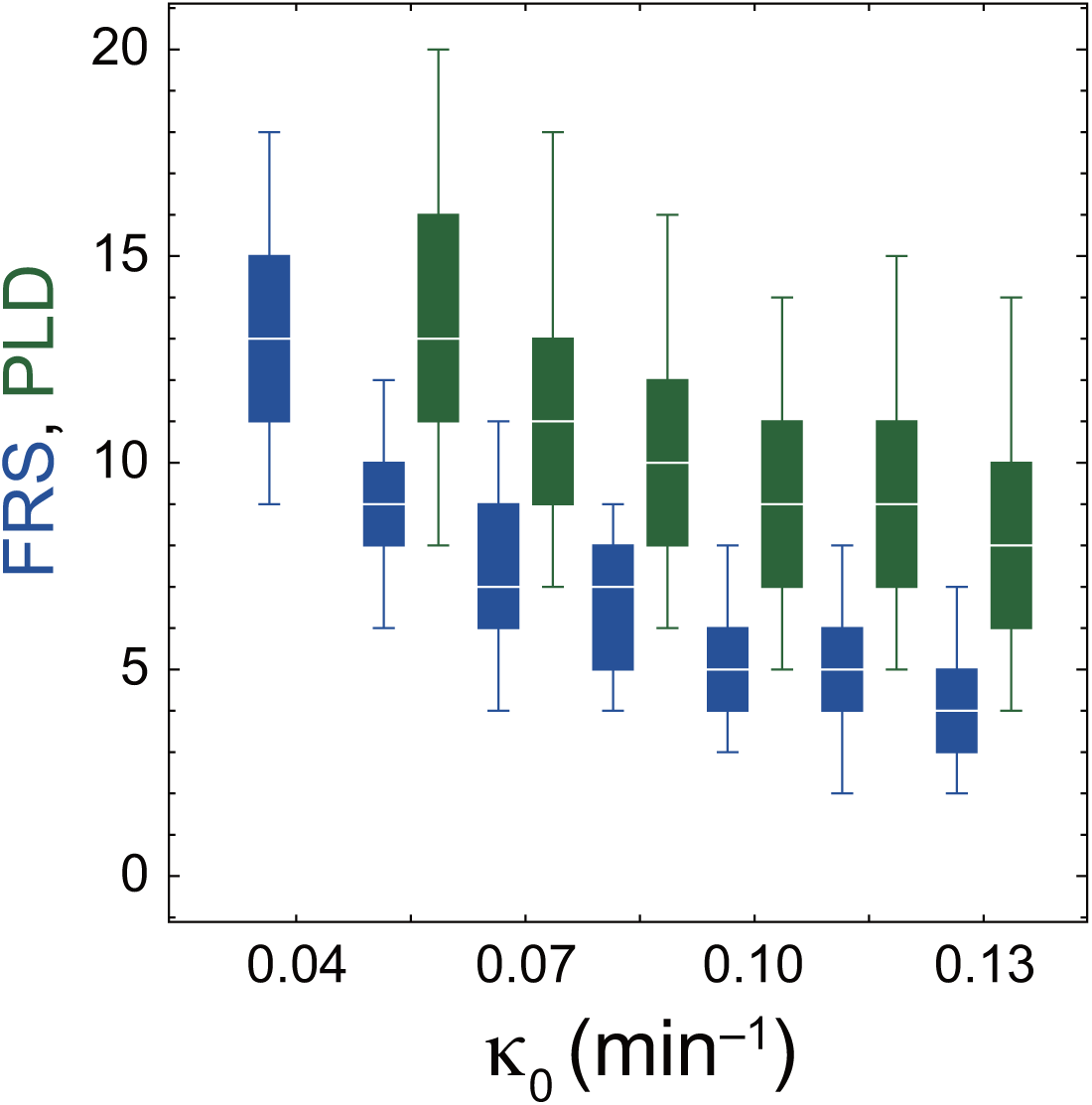
Coupling strength *κ*_*0*_ influences both FRS and PLD. The dependences of FRS (blue) and PLD (green) on *κ*_***0***_ are shown. The box-whisker plot indicates (0.05, 0.25, 0.75, 0.95) quantiles of FRS and PLD for 100 realizations of simulations. The white bars indicate median. Note that PLD for *κ*_***0***_ = 0.04 was not well-defined because a low value of coupling strength could not maintain a higher level of synchrony *Z*(*t*) > *Z*_*c*_ in the presence of cell addition with random phase values. Even when a simulation was started from the completely synchronized state, *Z*(*t*) sometimes became lower than *Z*_*c*_. Therefore, the result of PLD for a such small value of *κ*_***0***_ is not plotted in the figure.

**Figure S10.**
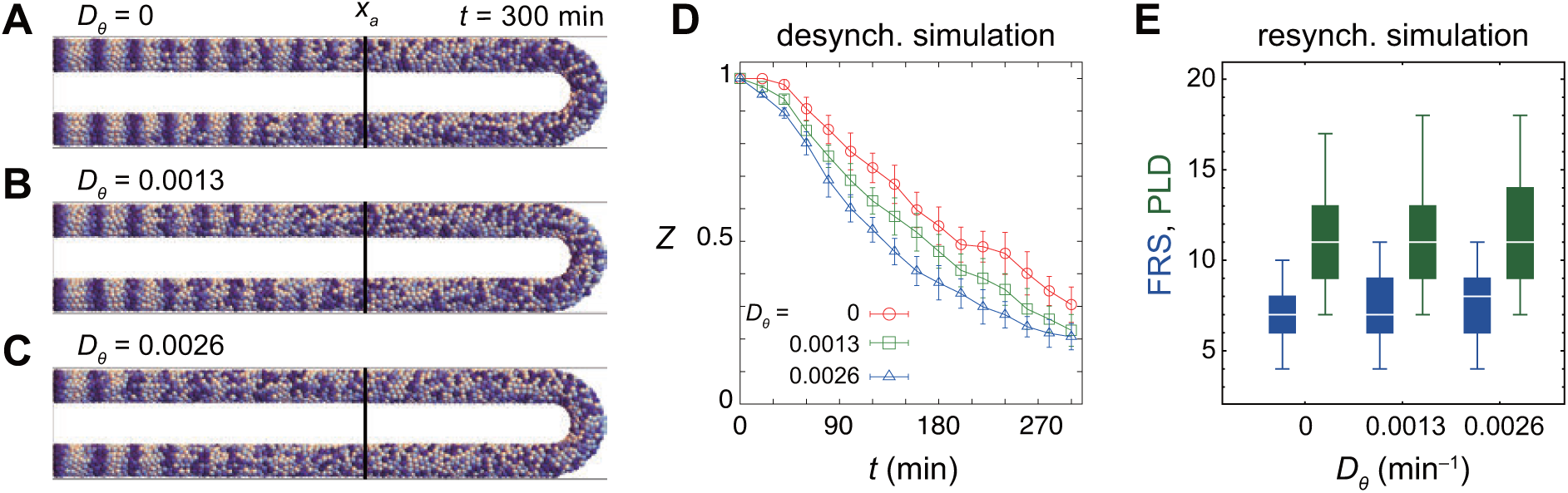
Dependence of desynchronization and resynchronization on phase noise intensity *D*_*θ*_. (A)-(C) Snapshots of spatial phase patterns in desynchronization simulations with (A) the phase noise intensity *D*_*θ*_ = 0 min^−1^, (B) *D*_*θ*_ = 0.0013 min^−1^ and (C) *D*_*θ*_ = 0.0026 min^−1^. In these simulations, coupling strength was set to zero *κ* = 0 in Eq. (10). The black vertical lines indicate the position of the anterior end of the PSM *x*_*a*_. (D) Time evolution of local phase order parameter *Z* at *x*_*a*_ for different values of *D*_*θ*_ in the desynchronization simulations with *κ* = 0. Marks and error bars indicate averages and standard deviations of *Z*, respectively, for 10 realizations of simulations. (E) Dependence of FRS (blue) and PLD (green) on the phase noise intensity *D*_*θ*_ in resynchronization simulations in a constant tissue with *κ*_***0***_ = 0.07. The box-whisker plot indicates (0.05, 0.25, 0.75, 0.95) quantiles of FRS and PLD for 100 realizations of simulations. The white bars indicate median. In all panels, the PSM length and advection pattern were constant over time.

**Figure S11.**
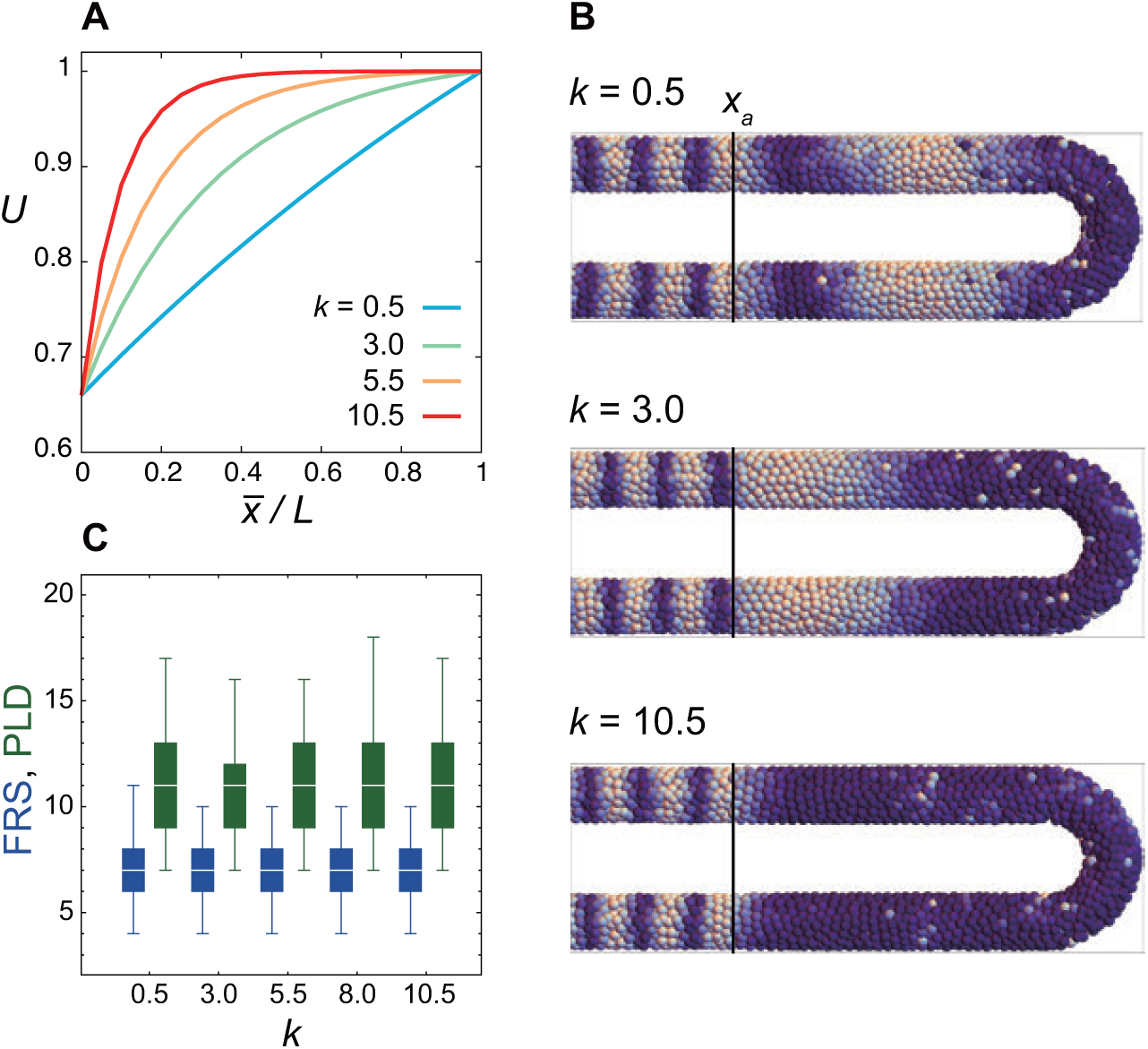
Weak dependence of FRS and PLD on the shape parameter *k* of the frequency profile. (A) Frequency profile *U*(x) for different values of *k* in Eq. (11) in the Materials and Methods. 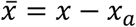 where *x*_*a*_ is the position of the anterior end of the PSM. (B) Snap shots of spatial phase patterns with different values of *k*. Simulations were started from synchronized initial conditions for illustration. *x*_*a*_ is indicated by the black vertical lines. (C) Dependence of FRS (blue) and PLD (green) on *k* in resynchronization simulations from random initial conditions. The box-whisker plot indicates (0.05, 0.25, 0.75, 0.95) quantiles of FRS and PLD for 100 realizations of simulations. The white bars indicate median.

**Figure S12.**
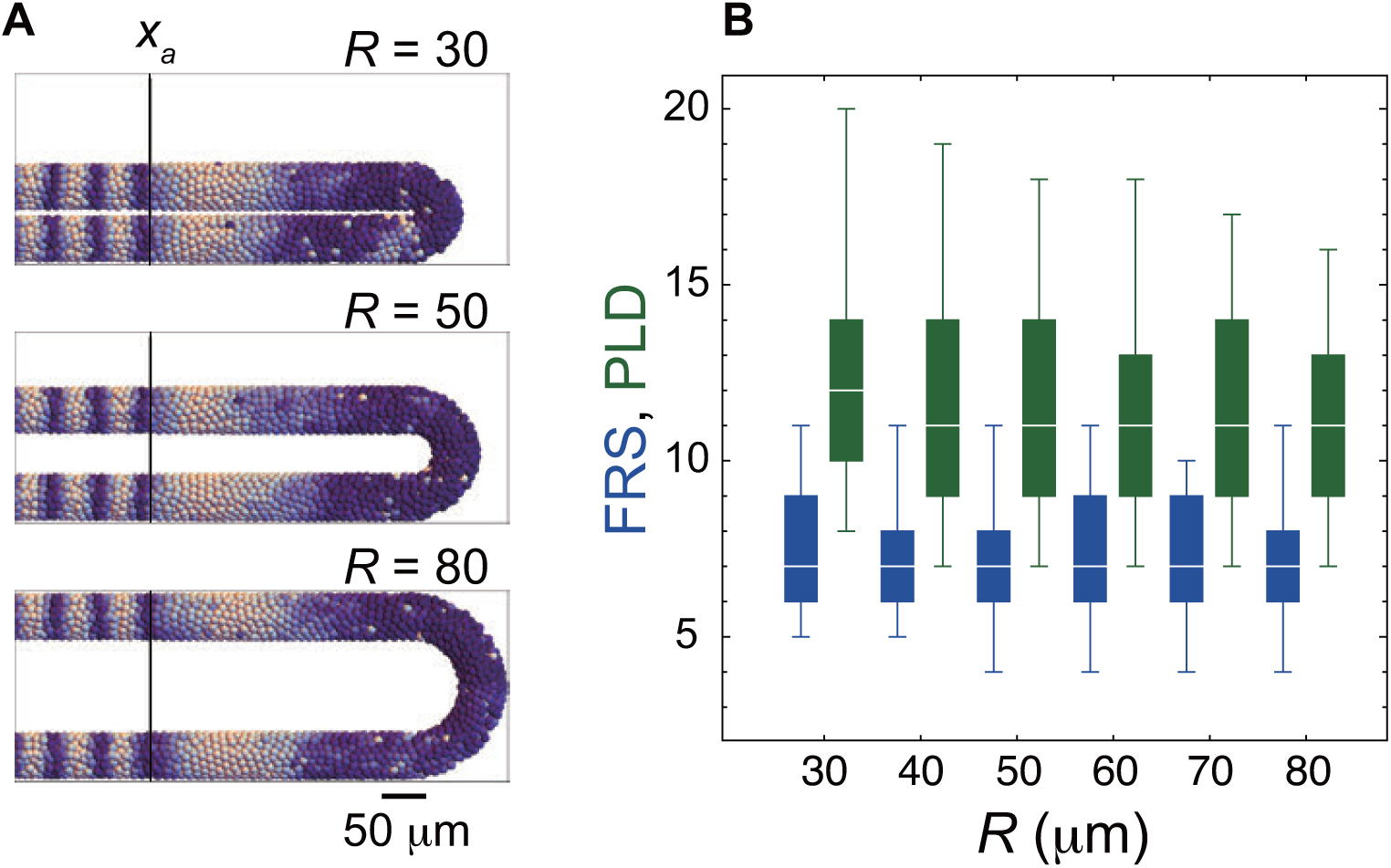
Weak dependence of FRS and PLD on the torus radius for the tailbud *R*. (A) Snap shots of the spatial phase patterns in the PSM with different values of *R*. The black vertical lines indicate the anterior end of the PSM *x*_*a*_. Simulations were started from synchronized initial conditions for illustration. (B) Dependence of FRS (blue) and PLD (green) on *R* in resynchronization simulations. The box-whisker plot indicates (0.05, 0.25, 0.75, 0.95) quantiles of FRS and PLD for 100 realizations of simulations. The white bars indicate median.

**Figure S13.**
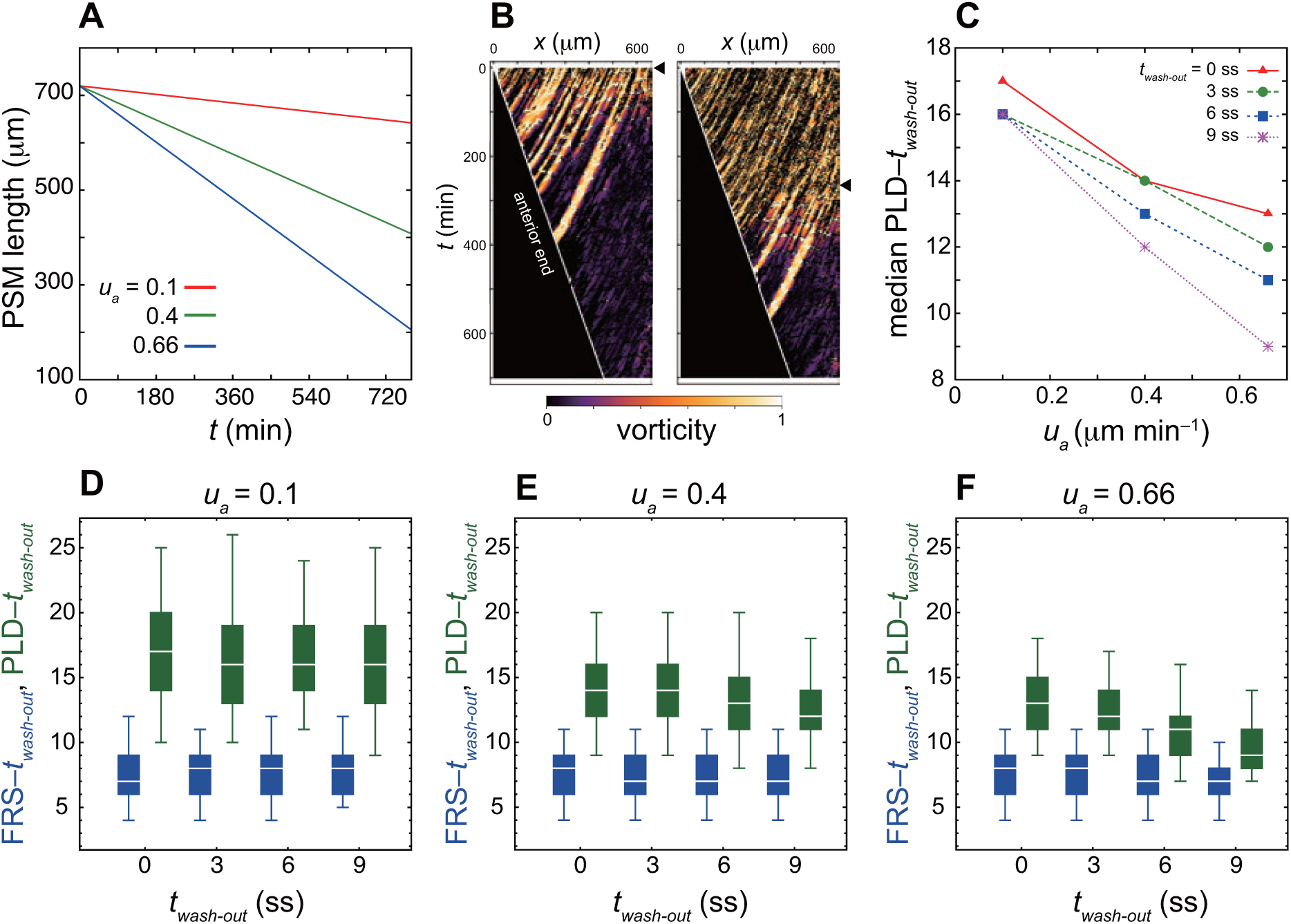
PSM shortening decreases time to PLD whereas it does not affect time to FRS. (A) Time evolution of the PSM length at different speed of PSM shortening *u*_*a*_. (B) Kymographs of vorticity for washout time at *t*_*wash-out*_ = 0 min (0 somite-stage, ss: left) and 270 min (9 ss: right). The black triangles indicate timing of washout. The white solid lines indicate the position of the anterior end of the PSM *x*_*a*_. Vorticity across the right PSM is shown. Trajectories of counter-clockwise vortices for *t*_*wash-out*_ = 0 min and those of clockwise vortices for *t*_*wash-out*_ = 270 min are presented. (C) Dependence of time to PLD, PLD – *t*_*wash-out*_, on *u*_*a*_ for different washout timing *t*_*wash-out*_. Medians for 100 realizations of simulations are plotted. (D)-(F) Dependences of time to FRS, FRS – *t*_*wash-out*_ (blue) and time to PLD, PLD – *t*_*wash-out*_ (green) on *t*_*wash-out*_ for different values of *u*_*a*_. (D) *u*_*a*_ = 0.1, (E) 0.4 and (F) 0.66. The box-whisker plot indicates (0.05, 0.25, 0.75, 0.95) quantiles of times to FRS and PLD for 100 realizations of simulations. The white bars indicate median.

**Figure S14.**
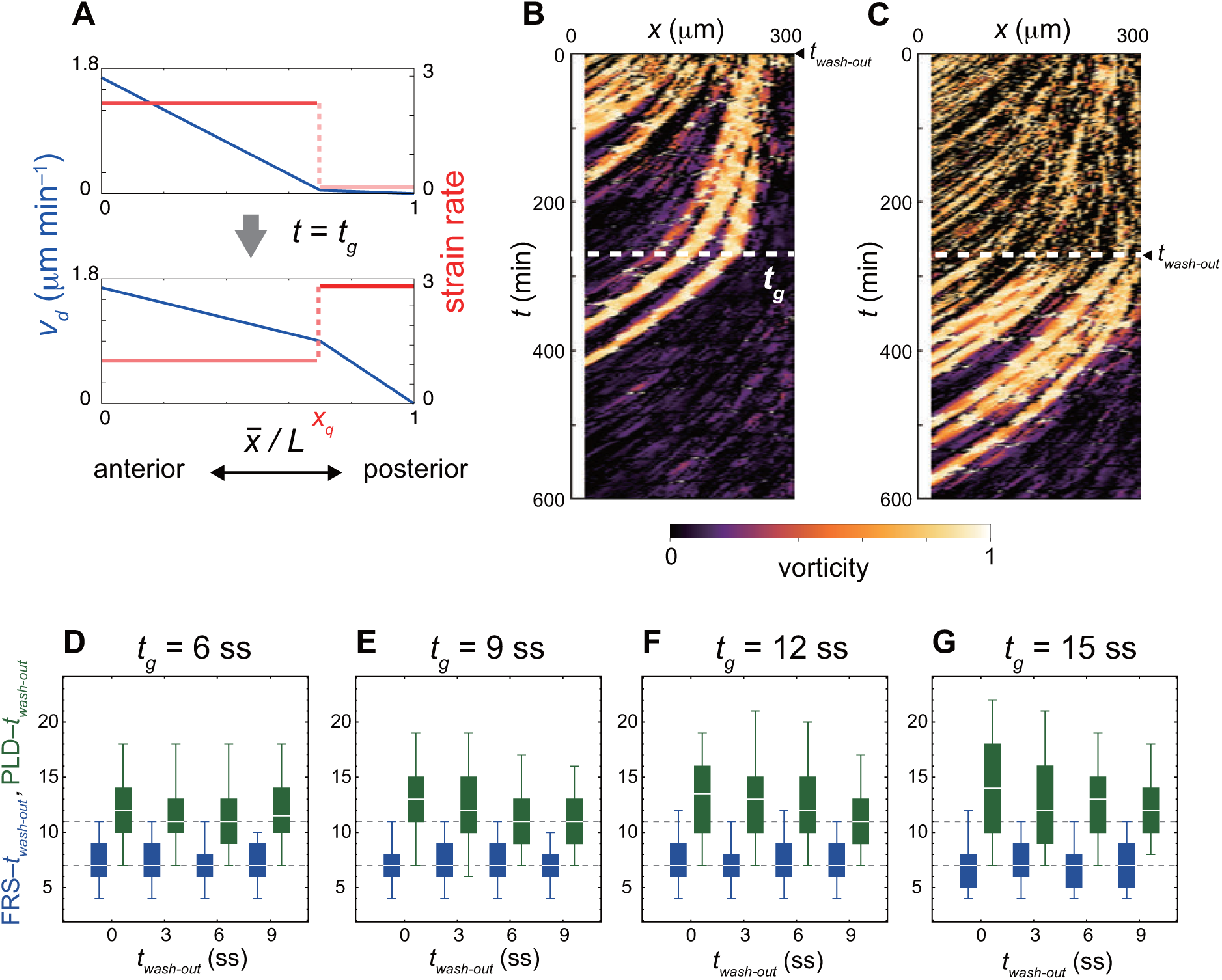
Change in advection pattern increases the time to PLD for earlier DAPT washout. (A) Change in advection pattern at *t* = *t*_*g*_. The blue and red lines indicate the advection speed *v*_*d*_ and strain rate, respectively. 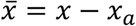 where *x*_*a*_ is the position of the anterior end of the PSM. (B), (C) Kymographs of vorticity for washout at (B) *t*_*wash-out*_ = 0 min (0 somite stage; ss) and (C) 270 min (9 ss). White dotted lines indicate *t*_*g*_ at which the advection pattern changes (*t*_*g*_ = 9 ss). Vorticity across the right PSM is shown. Trajectories of clockwise vortices for *t*_*wash-out*_ = 0 min and those of counter-clockwise vortices for *t*_*wash-out*_ = 270 min are presented. (D)-(G) Dependence of times to FRS (blue) and PLD (green) on washout time *t*_*wash-out*_ with change in advection pattern at (D) *t*_*g*_ = 6 ss, (E) 9 ss, (F) 12 ss, and (G) 15 ss. The gray dotted lines indicate times to FRS and PLD for the constant tissue shown in Fig. 4A in the main text. The box-whisker plot indicates (0.05, 0.25, 0.75, 0.95) quantiles of times to FRS and PLD for 100 realizations of simulations. The white bars indicate median.

**Figure S15.**
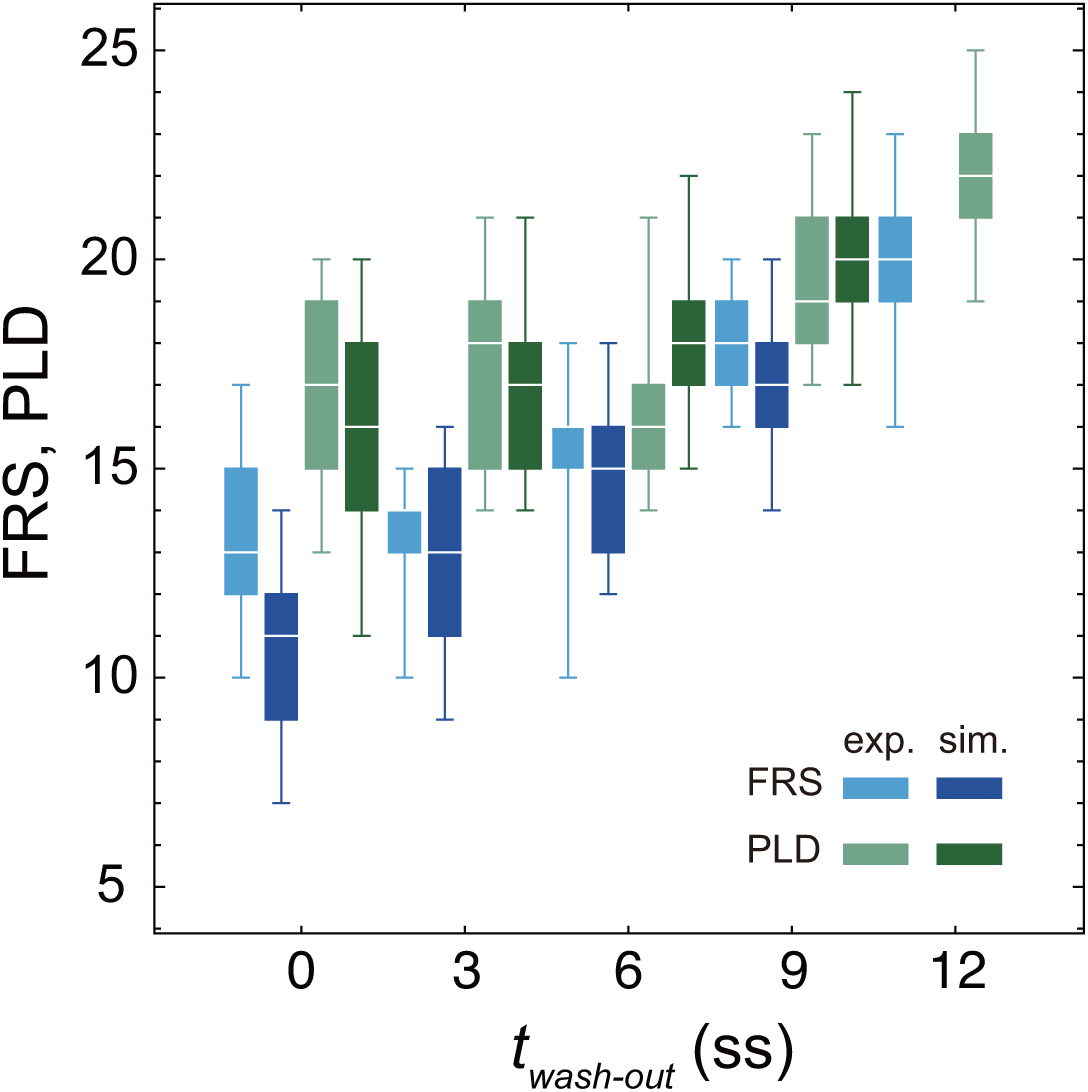
Dependence of embryonic FRS and PLD on DAPT washout time. Experimental FRS (light blue) and PLD (light green) for zebrafish embryos are plotted together with FRS and PLD (darker colors) obtained by 100 realizations of numerical simulations. The physical model for the simulations included PSM shortening, changes in advection pattern and coupling strength. The box-whisker plot indicates (0.05, 0.25, 0.75, 0.95) quantiles of FRS and PLD. The white bars indicate median. Due to the lack of the information about the formation of final segments, we did not perform simulations for the latest washout *t*_*wash-out*_ = 12 somite-stage (ss), see the main text. exp.: experiment. sim.: simulation.

**Figure S16.**
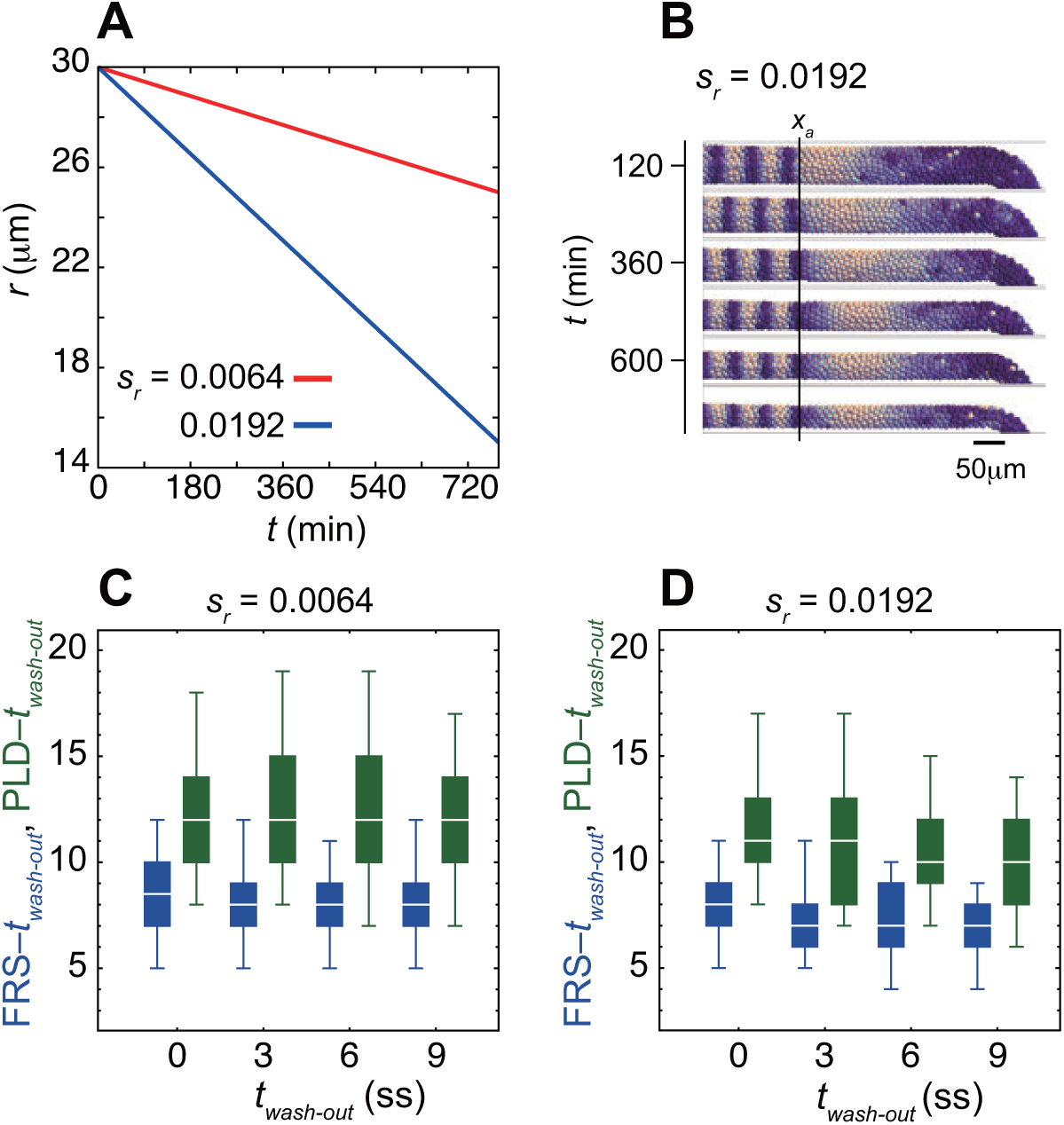
Decrease in the PSM radius *r* over developmental stages reduces both time to FRS and PLD. (A) PSM radius *r* as a linear function of time. *s*_*r*_ represents the absolute value of slope of a linear line. (B) Snapshots of spatial phase patterns of a simulation with decreasing PSM radius. The simulation was started from a synchronized initial condition for illustration. The black vertical line indicates the position of the anterior end of the PSM *x*_*a*_. (C), (D) Dependence of times to FRS (blue) and PLD (green) on *t*_*wash-out*_ with (C) *s*_*r*_ = 0.0064 μm min^− 1^ (red line in A) and (D) *s*_*r*_ = 0.0192 μm min^−1^ (blue line in A). Only the PSM radius changed over time, whereas all the other parameters were kept constant. Simulations were started from random initial conditions. The box-whisker plot indicates (0.05, 0.25, 0.75, 0.95) quantiles of times to FRS and PLD for 100 realizations of simulations. The white bars indicate median. ss: somite stage.

**Figure S17.**
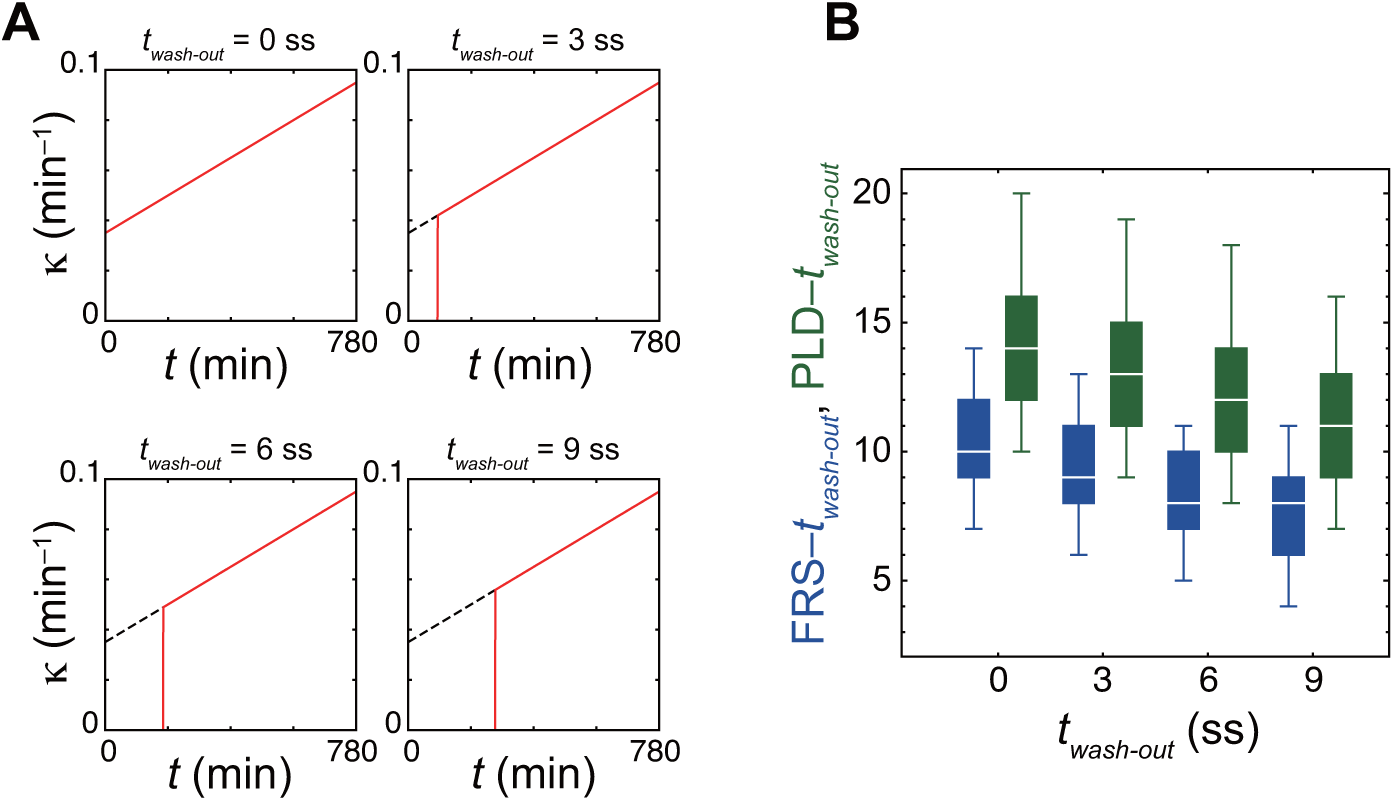
Decrease in time to FRS by an increase in the coupling strength over developmental stages in simulations. (A) Time series of the coupling strength *κ* with different DAPT washout time in the physical model. Black dashed lines indicate the increase of coupling strength over developmental stages in the absence of DAPT treatment. Red solid lines indicate coupling strength before and after DAPT washout. For simplicity, we assume that cells restore coupling to its original value immediately after washout as represented by the vertical red lines. (B) Dependence of times to FRS (blue) and PLD (green) on DAPT washout time in simulations. The box-whisker plot indicates (0.05, 0.25, 0.75, 0.95) quantiles of times to FRS and PLD for 100 realizations of simulations. The white bars indicate median. The effect of increasing coupling strength shown in (A) was examined with the constant PSM length and advection pattern. ss: somite stage.

**Figure S18.**
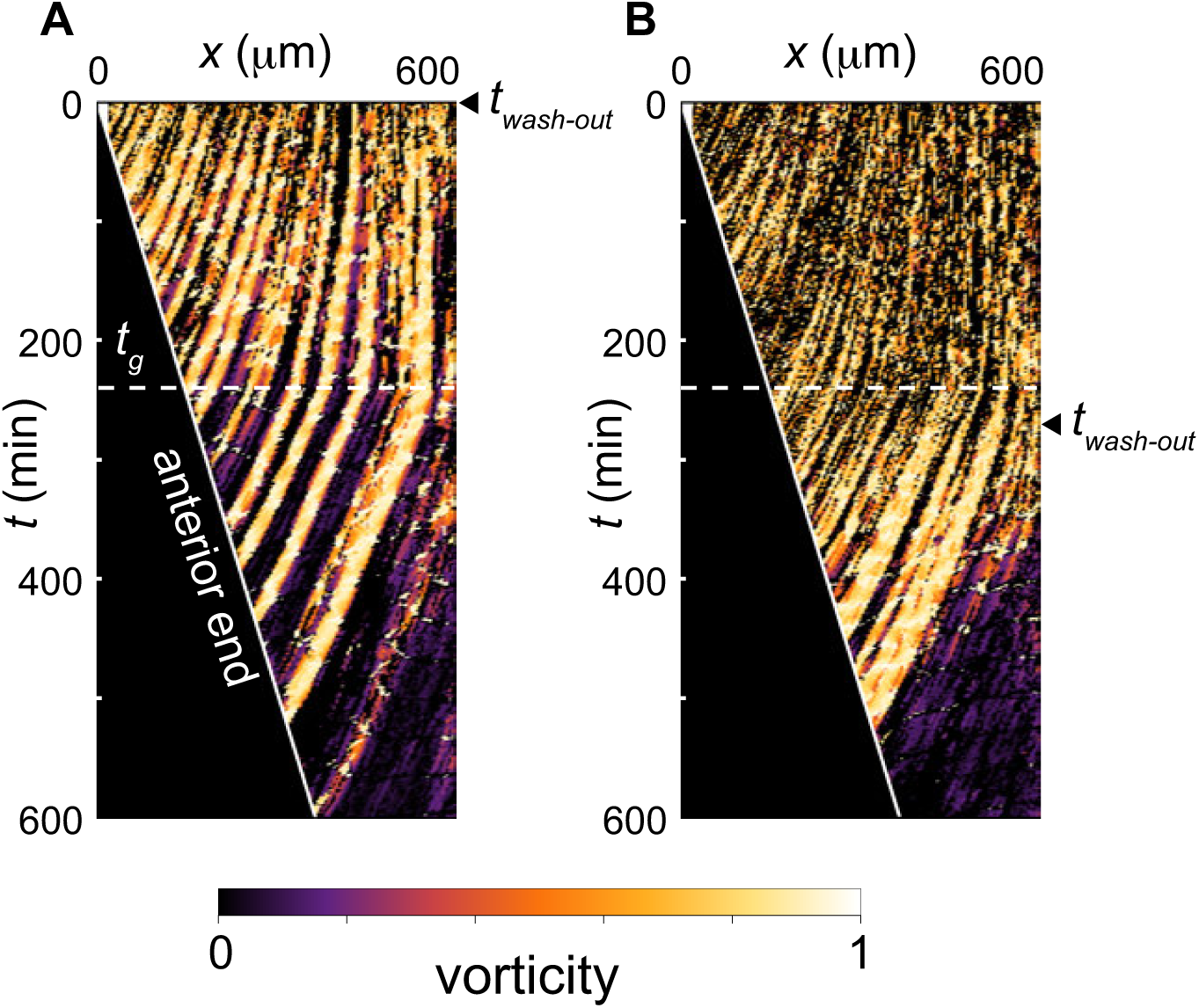
Trajectories of phase vortices in the physical model including PSM shortening, changes in advection pattern and coupling strength. (A), (B) Kymographs of vorticity for (A) *t*_*wash-out*_ = 0 somite stage (ss; 0 min) and (B) *t*_*wash-out*_ = 9 ss (270 min). The white horizontal dotted lines indicate time *t*_*g*_ at which advection pattern changes, *t*_*g*_ = 240 min. The solid white lines indicate the position of the anterior end of the PSM *x*_*a*_. Vorticity across the right PSM is shown. Trajectories of clockwise vortices are presented.

**Figure S19.**
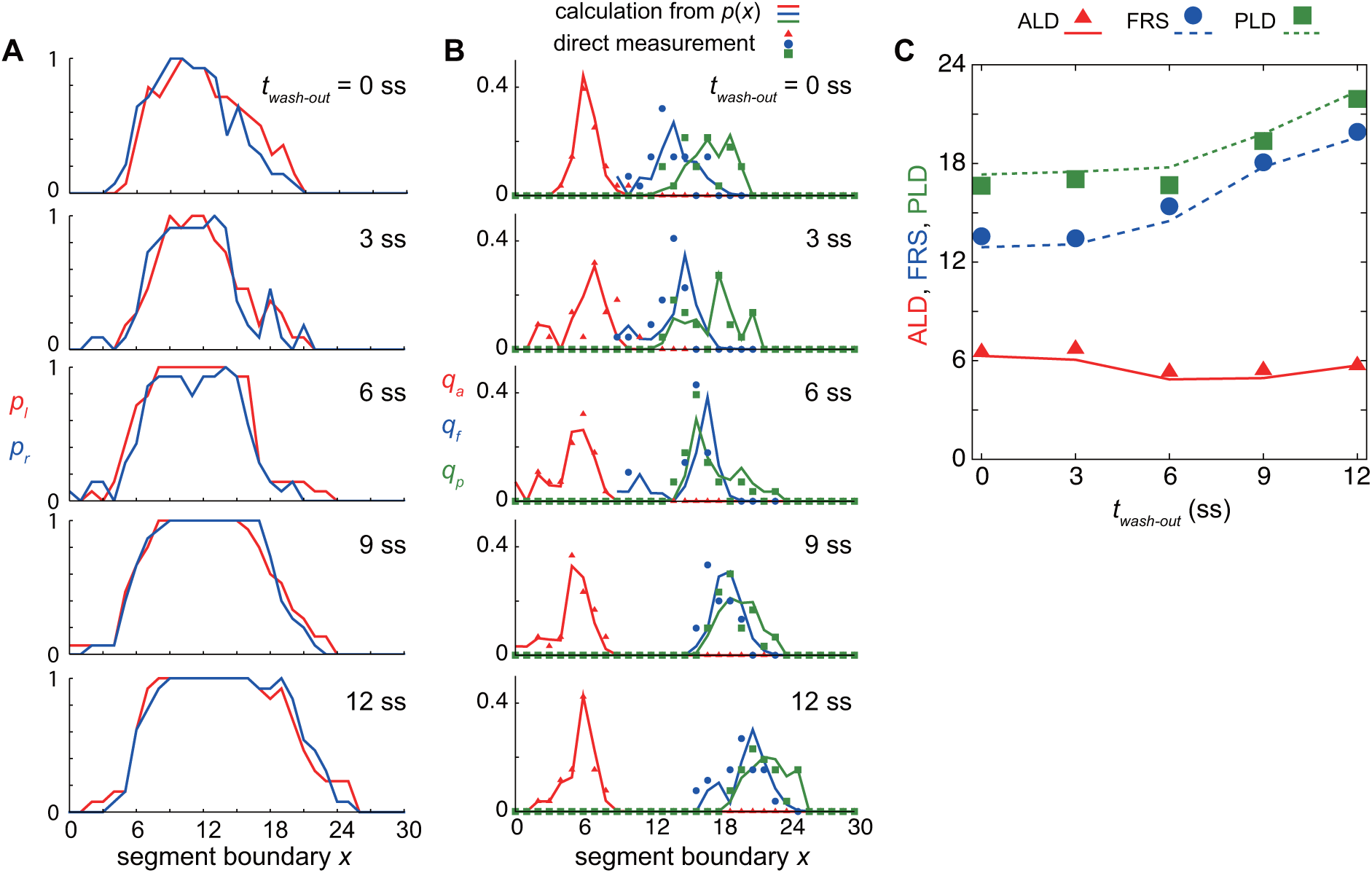
ALD, FRS and PLD calculated with the spatial distribution of defective segments. (A) Spatial distribution of defective segments in left (red lines: *p*_*l*_) and right (blue lines: *p*_*r*_) sides of embryos. (B) Spatial probability distribution of ALD *q*_*a*_ (red lines), FRS *q*_*f*_ (blue lines) and PLD *q*_*p*_ (green lines) for different DAPT washout time calculated from the spatial distribution of defective segments *p*(*x*). Symbols indicate direct measurement of *q*_*a*_ (triangles), *q*_*f*_ (squares) and *q*_*p*_ (circles). See Supporting Information for calculation. (C) ALD, FRS and PLD as a function of DAPT washout timing. Symbols indicate results of direct measurement of these quantities. Lines indicate results of probability calculations described in Supporting Information. ss: somite stage.

**Figure S20.**
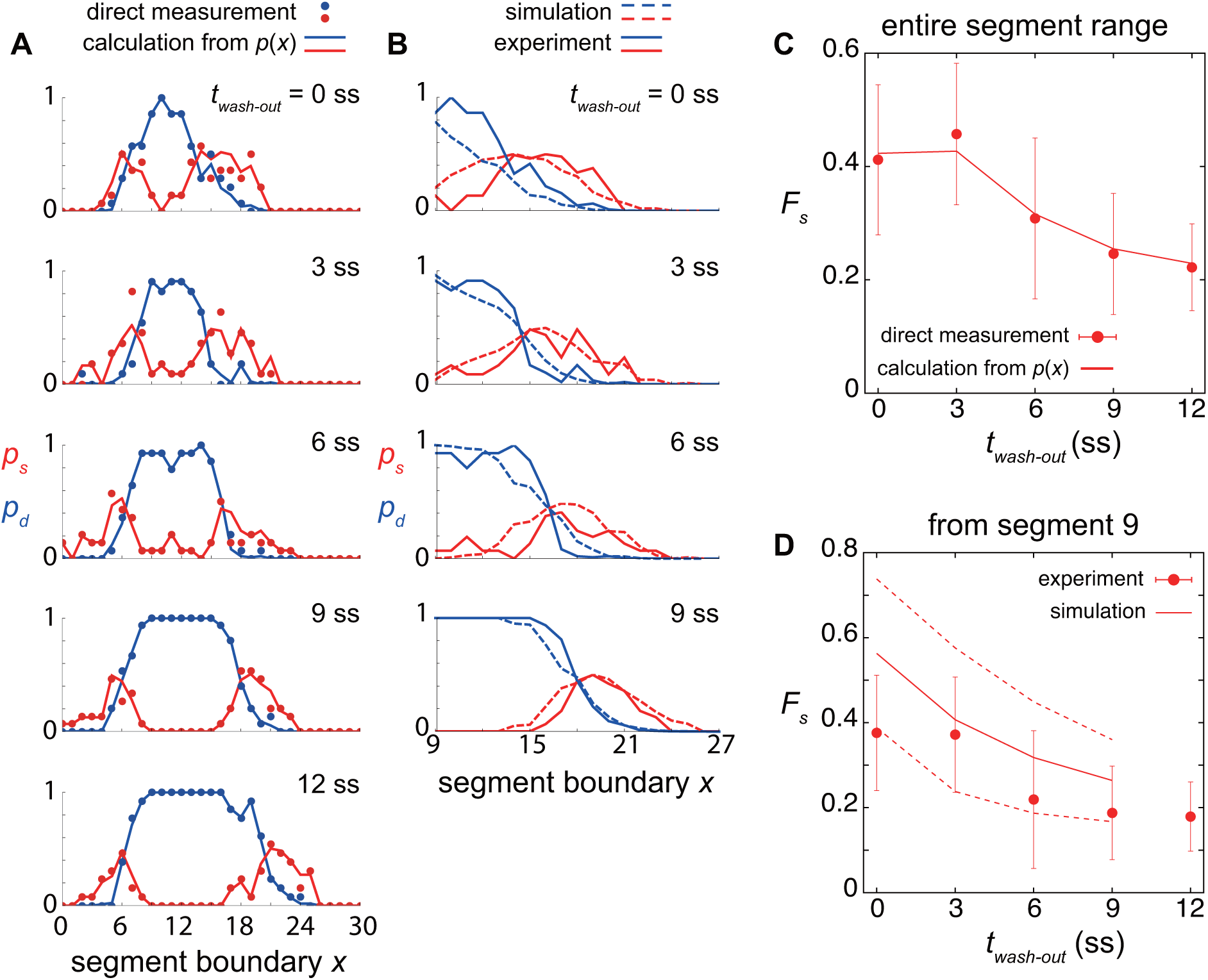
Dependence of single and double defects on DAPT washout timing. (A) Spatial probability distribution of single defects *p*_*s*_ (red lines) and double defects *p*_*d*_ (blue lines) for different DAPT washout time. *p*_*s*_ and *p*_*d*_ were calculated from the experimental spatial distributions of defective segments. Filled circles indicate direct measurement of *p*_*s*_ (red) and *p*_*d*_ (blue). (B) Comparison of *p*_*s*_ (red) and *p*_*d*_ (blue) between simulations (dotted lines) and experiment (solid lines). (C) Dependence of fraction of single defects *F*_*s*_ on DAPT washout timing. Circles indicate *F*_*s*_ obtained by direct measurement. Error bars indicate the standard deviation. The line indicates *F*_*s*_ calculated from the spatial distribution of defective segments. See Supporting Information for the calculation. *F*_*s*_ was computed using entire segment range. (D) Comparison of *F*_*s*_ obtained by the physical model (lines) with that in experiment (circles). The solid and dotted lines indicate mean and standard deviations of simulation data, respectively. *F*_*s*_ was measured from segment 9 for the comparison between simulations and experiment.

**Table S1.** Parameter values used in Figs. 2 and 3.

| tissue geometry |  |  |
| --- | --- | --- |
| parameter | description | value |
| $L_x$ | tip of the tailbud | 385 $\mu\text{m}$ |
| $r$ | PSM radius | 25 $\mu\text{m}$ |
| $R$ | radius of the half torus for the tailbud | 60 $\mu\text{m}$ |
| $X_c$ | $x$ position of the center of the half torus | 300 $\mu\text{m}$ |
| $d_c$ | cell diameter | 11 $\mu\text{m}$ |
| $s_r$ | magnitude of radius change in Eq. (15) | 0 $\mu\text{m min}^{-1}$ |

| cell advection Eq. (2) |  |  |
| --- | --- | --- |
| $v_a$ | advection speed at the anterior end of the PSM | 1.67 $\mu\text{m min}^{-1}$ |
| $v_p$ | strain rate at the posterior domain | 3 $\mu\text{m min}^{-1}$ |
| $x_q$ | $x$ position at which the strain rate changes | 0.3 |
| $t_g$ | time at which the advection pattern changes | - |

| cell movement and intercellular force Eqs. (3)-(9) |  |  |
| --- | --- | --- |
| $v_s$ | maximum cell movement speed at the tip of the tailbud | 1.0 $\mu\text{m min}^{-1}$ |
| $X_v$ | normalized length scale of cell mobility gradient | 0.4 |
| $h$ | steepness of cell mobility gradient | 3 |
| $D_\phi$ | polarity noise intensity | 0.026 $\text{min}^{-1}$ |
| $\mu$ | intercellular force coefficient | 8.71 $\mu\text{m min}^{-1}$ |
| $\mu_b$ | boundary force coefficient | 20 $\mu\text{m min}^{-1}$ |
| $r_b$ | length scale of boundary force | 1 $\mu\text{m}$ |

| phase equation Eqs. (10)-(12) |  |  |
| --- | --- | --- |
| $\omega_0$ | autonomous frequency at the tip of the tailbud | 0.2094 $\text{min}^{-1}$ |
| $\sigma$ | difference in the frequency between anterior and posterior ends of the PSM | 0.66 |
| $k$ | shape parameter for frequency profile | 3.07 |
| $D_\theta$ | phase noise intensity | 0.0013 $\text{min}^{-1}$ |
| $\kappa_s$ | rate of the increase in coupling strength | 0 $\text{min}^{-2}$ |
| $\kappa_0$ | offset of the coupling strength | 0.07 $\text{min}^{-1}$ |
| $T_a$ | period at the anterior end of the PSM $x_a$ | 30 min |

| PSM shortening Eqs. (13), (14) |  |  |
| --- | --- | --- |
| $u_a$ | speed of PSM shortening | $0 \mu\text{m min}^{-1}$ |
| $S$ | segment size | $50 \mu\text{m}$ |
| $c$ | sum of $u_a$ and $v_a$ | $1.67 \mu\text{m min}^{-1}$ |
| $\rho_0$ | cell density | $0.0015 \mu\text{m}^{-3}$ |
| $\zeta$ | anterior limit of cell addition | $100 \mu\text{m}$ |

**Table S2.** Parameter values used in Fig. 4 and Figs. S5-S18, S20.

|  |  |
| --- | --- |
| Fig. 4 | (A) All the parameter values are the same as in Table S1.<br>(B) $L_x = 720$, $X_c = 635$, $u_a = 0.66$, $v_a = 1.01$, and $\omega_0 = 0.2001$. Values of all the other parameters are the same as in Table S1.<br>(C) $t_g = 270$ min, $v_p = 0.16$ for $t < t_g$ and $v_p = 3$ for $t \geq t_g$. Values of all the other parameters are the same as in Table S1.<br>(E), (F) $L_x = 720$, $X_c = 635$. $u_a = 0.66$, $v_a = 1.01$ and $\omega_0 = 0.2001$. $t_g = 240$ min, $v_p = 0.16$ for $t < t_g$ and $v_p = 3$ for $t \geq t_g$. $\kappa_s = 0.06/780$, and $\kappa_0 = 0.035$. Values of all the other parameters are the same as in Table S1. |
| Fig. S5 | (A)-(C) $v_s = 0.2$-1.6. Values of all the other parameters are the same as in Table S1. |
| Fig. S6 | (A)-(C) $L_x = 300, 400, 500, 600$. $X_c = 215, 315, 415, 515$. All the other parameters are the same as in Table S1. |
| Fig. S7 | (B) $v_p = 0.16$ for the left panel, $v_p = 3$ for right panel. (C) $v_p = 0.16 - 4.42$. All the other parameters are the same as in Table S1. |
| Fig. S8 | $r = 15 - 30$. All the other parameters are the same as in Table S1. |
| Fig. S9 | $\kappa_s = 0$ and $\kappa_0 = 0.04 - 0.13$. All the other parameters are the same as in Table S1. |
| Fig. S10 | (A)-(D) $\kappa_s = \kappa_0 = 0$. $D_\theta = 0, 1.3 \times 10^{-3}, 2.6 \times 10^{-3}$. All the other parameters are the same as in Table S1. (E) $\kappa_s = 0$, $\kappa_0 = 0.07$. $D_\theta = 0, 1.3 \times 10^{-3}, 2.6 \times 10^{-3}$. All the other parameters are the same as in Table S1. |
| Fig. S11 | $k = 0.5, 3.0, 5.5, 8.0, 10.5$. All the other parameters are the same as in Table S1. |
| Fig. S12 | $R = 30 - 80$. All the other parameters are the same as in Table S1. |
| Fig. S13 | $L_x = 720$, $X_c = 635$. For $u_a = 0.66$, $v_a = 1.01$ and $\omega_0 = 0.2001$ in (A), (B), (C), and (F). For $u_a = 0.1$, $v_a = 1.57$ and $\omega_0 = 0.2082$ in (A), (C) and (D). For $u_a = 0.4$, $v_a = 1.27$ and $\omega_0 = 0.2042$ in (A), (C) and (E). All the other parameters are the same as in Table S1. |
| Fig. S14 | In (B)-(G), $v_p = 0.16$ for $t < t_g$ and $v_p = 3$ for $t \geq t_g$. (B), (C) and (E) $t_g = 270$ min. (D) $t_g = 180$ min. (F) $t_g = 360$ min. (G) $t_g = 450$ min. All the other parameters are the same as in Table S1. |
| Fig. S15 | Parameter values are same as in Fig. 4 (E), (F). |
| Fig. S16 | (C) $s_r = 0.0064$ and (D) $s_r = 0.0192$. In both (C) and (D), $r_0 = 30 \mu\text{m}$. All the other parameters are the same as in Table S1. |
| Fig. S17 | $\kappa_s = 0.06/780$, and $\kappa_0 = 0.035$. All the other parameters are the same as in Table S1. |
| Fig. S18 | Parameter values are same as in Fig. 4 (E), (F). |
| Fig. S20 | (B) and (D) Parameter values are same as in Fig. 4 (E), (F). |

## Movie legends

**Movie 1** Traveling phase waves in a constant tissue. The simulation was started from a completely synchronized state. The color indicates (1 + sin *θ*_*i*_)/2. The value of the coupling strength is constant *κ*_0_ = 0.07 for *t* > 0. The black vertical line indicates the position of the anterior end of the PSM, *x*_*a*_. Values of parameters are the same in Fig. 2 in the main text and listed in Table S1.

**Movie 2** Resynchronization simulation in a constant PSM tissue. The simulation was started from random initial phases. The color indicates (1 + sin *θ*_*i*_)/2. The value of coupling strength is constant *κ*_0_ = 0.07 for *t* > 0. The black vertical line indicates the position of the anterior end of the PSM, *x*_*a*_. Values of parameters are the same in Fig. 3 in the main text and listed in Table S1.

**Movie 3** Resynchronization simulation with the changes in PSM length. DAPT washout time is *t*_*wash-out*_ = 0 somite stage (0 min). The simulation was started from random initial phases. The color indicates (1 + sin *θ*_*i*_)/2. The black vertical line indicates the position of the anterior end of the PSM, *x*_*a*_. Values of parameters are the same in Fig. 4B in the main text and listed in Tables S1, S2.

**Movie 4** Resynchronization simulation with the changes in PSM advection pattern. The advection pattern changes at *t*_*a*_ = 9 somite stage (ss; 270 min). DAPT washout time is *t*_*wash-out*_ = 0 ss (0 min). The simulation was started from random initial phases. The color indicates (1 + sin *θ*_*i*_)/2. The black vertical line indicates the position of the anterior end of the PSM, *x*_*a*_. Values of parameters are the same in Fig. 4C in the main text and listed in Tables S1, S2

**Movie 5** Resynchronization simulation with the changes in PSM length, advection pattern and value of coupling strength. DAPT washout time is *t*_*wash-out*_ = 0 somite stage (0 min). The simulation was started from random initial phases. The color indicates (1 + sin *θ*_*i*_)/2. The black vertical line indicates the position of the anterior end of the PSM, *x*_*a*_. Values of parameters are the same in Fig. 4E in the main text and listed in Tables S1, S2.

**Movie 6** Resynchronization simulation with the changes in PSM length, advection pattern and value of coupling strength. DAPT washout time is *t*_*wash-out*_ = 9 somite stage (270 min). The simulation was started from random initial phases. The color indicates (1 + sin *θ*_*i*_)/2. The black vertical line indicates the position of the anterior end of the PSM, *x*_*a*_. Values of parameters are the same in Fig. 4E in the main text and listed in Tables S1, S2.

**Movie 7** Formation of phase vortices in a resynchronization simulation in a cuboid domain, 110×110×55 μm^3^. The color indicates (1 + sin *θ*_*i*_)/2. The number of oscillators is *N* = 998. Frequencies of all the oscillators are identical, ω = 0.2094 min^−1^. Values of the other relevant parameters in Eq. (1) in the supporting information are: *κ*_0_ = 0.07 min^−1^, *κ*_s_ = 0 min^−1^, *D*_θ_ = 0.0013 min^−1^, μ = 8.71 μm min^− ^1^^, *d*_*c*_ = 11 μm, μ_*b*_ = 20 μm min^− ^1^^, and *r*_*b*_ = 1 μm.

## Supporting Information

### From local resynchronization to global pattern recovery in the zebrafish segmentation clock

### MATERIALS AND METHODS

#### Animals and Embryos

Zebrafish (*Danio rerio*) adult stocks were kept in 28°C fresh water under a 14:10 hour light:dark photoperiod. Embryos were collected within 30 minutes following fertilization and incubated in petri dishes with E3 media. Until the desired developmental stages (1), embryos were incubated at 28.5 °C. For whole mount *in situ* hybridization, PTU (1-phenyl 2-thiourea) at a final concentration of 0.003% was added before 12 hours post fertilization (hpf) to prevent melanogenesis. All wildtypes were *AB* strain.

#### DAPT treatment and washout

DAPT treatment was carried out as previously described (2). 50 mM DAPT stock solution (Merck) was prepared in 100% DMSO (Sigma) and stored in a small volume at −20 °C. Embryos in their chorions were transferred to 12-well plates at 2 hpf in 1.4 ml E3 medium with 20 embryos per well. 50 μM DAPT in E3 medium was prepared immediately before the treatment. To prevent precipitation, the DAPT stock solution was added into E3 medium while vortexing, and then filtered by 0.22 μm PES syringe filter (Millipore). DAPT treatment was initiated by replacing E3 medium with E3/DAPT medium at desired stage. At 9.5 (0 somite stage: ss), 11 (3 ss), 12.5 (6 ss), 14 (9 ss), or 15.5 hpf (12 ss), DAPT was washed out at least twice with fresh E3 medium + 0.03% PTU. Embryos were dechorionated and fixed at 36 hpf. All experimental steps were incubated at 28.5 °C, except for short operations, e.g., washing out, which were at room temperature.

#### Whole-mount *in situ* hybridization and segmental defect scoring

*In situ* hybridization was performed according to previously published protocols (3). Digoxigenin-labelled *xirp2a* (clone: *cb1045*) riboprobe was as previously described (2). Stained embryos were visually scored under an Olympus SZX-12 stereomicroscope and images were acquired with a QImaging Micropublisher 5.0 RTV camera. Defective segment boundaries were scored as previously described (2), with the addition that left and right (LR) sides of the embryo were scored and assessed separately. Since boundary formation in unperturbed embryos is extremely reliable, with errors occurring in less than 1 in 1000 embryos, any interruption or fragmentation to the boundary, and/or alterations in spacing, or alignment was recorded as a defect. The following observables were collected for each LR side: an anterior limit of defects (ALD), i.e., the position of the first defective boundary; a posterior limit of defects (PLD), i.e., the position of the last defective boundary; and the first recovered segment (FRS), i.e., the position of the first normal segment after the segment 9, as described below.

Each segment has anterior and posterior boundaries. For the segment *i* (*i* = 1, 2, 3, ..,), the anterior and posterior boundaries were numbered as *i* – 1 and *i*, respectively, Fig. S1A. Both ALD and PLD were numbered using the posterior boundary of the defective segment. For example, if the *j*th segment boundary was the last defective boundary, PLD was numbered as *j*. FRS was numbered using the anterior boundary of the recovered segment. For example, if the *k*th segment boundary was the first normal boundary after washout, the first normal segment was segment *k*, and FRS was numbered as *k* – 1, Fig. S1A. With this definition of FRS, if the first normal segment boundary after the washout was located immediately after the last defective boundary, the difference between PLD and FRS, PLD – FRS, was 0, Fig. S1C-G. Definitions of FRS and PLD for simulation date were the same as those for experimental data written here, and described in a later section in this supporting information.

#### Physical model of PSM and tailbud cells

##### 3D tissue geometry

We model the PSM and tailbud as a U-shaped domain in a 3D space, Fig. S2A. The left and right PSMs are represented as two tubular domains Ω_*l*_ and Ω_*r*_, respectively with radius *r*. The tailbud is described as a half toroidal domain Ω_*t*_ with a larger radius *R* and smaller radius *r*.

We implement this U-shaped domain in the spatial coordinate system as follows. The *x*-axis is along the anterior-posterior axis of the PSM. We set the initial position of the anterior end of the PSM at *x* = 0 and the posterior tip of the tailbud at *x* = *L*_*x*_. We denote the position of the anterior end of the PSM at time *t* as *x*_*a*_(*t*), so *x*_*a*_(0) = 0. The length of the tissue is *L* = *L*_*x*_ − *x*_*a*_. **X**_c_ denotes the position of the center of torus core curve. The position of posterior tip of the tailbud can then be written as *L*_*x*_ = *X*_*c*_ + *R* + *r*. The *y*-axis points along the left-right axis and *z*-axis along the dorsal-ventral axis of an embryo.

##### Reference frames: Lab reference frame

To describe cell movements and tissue deformations it is important to define a reference frame. A natural choice may be a reference frame which is at rest in the Lab, termed a Lab reference frame. For instance, the origin of *x*-axis can be set at the initial position of the anterior end of the PSM at *t* = 0. We write the position of the posterior tip of the tailbud in this Lab reference frame as 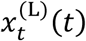, where superscript (L) indicates variables in the Lab reference frame. Because an embryo elongates posteriorly, 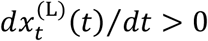. The position of anterior end of the PSM in the Lab reference frame 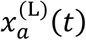 also changes over time due to the formation of new segments. In this reference frame, PSM length is 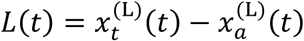. The rate of change of PSM length is 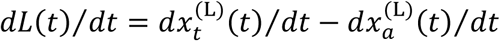. PSM length is constant if the velocity of the tailbud is the same as that of the anterior end of the PSM, that is 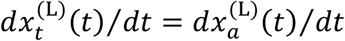, but it will be changing over time whenever 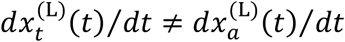. In this Lab reference frame, the position of a cell 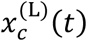 may change both due to tissue elongation and due to tissue deformations such as local tissue stretch.

##### Reference frames: Tailbud reference frame

In this study however, we mostly use the tailbud reference frame (t) unless noted otherwise. In this reference frame we measure the position of cells and tissue landmarks from the tailbud. The position of the tailbud is fixed at 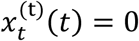 for all time *t*, so 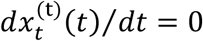. The position of a tissue landmark *l* is 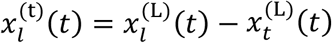. In this tailbud frame, the velocity of a landmark is zero if it moves with the same velocity as the tailbud in the Lab frame. In other words, a tissue landmark moves relative to the tailbud when their velocities in the Lab reference frame are different. The positions of cells in the tailbud reference frame may change over time, and this will enter as cell advection in the cell equations of motion as we describe below. In the following we omit the superscript (t) for the tailbud reference frame to alleviate the notation.

##### Reference frames: Computational implementation

For the implementation of the tailbud frame for numerical simulations, we fix the position of the posterior tip of the tailbud at *x* = *L*_*x*_. In contrast, cells positions and other tissue landmarks such as the anterior end of the PSM change over time. We model cell displacement relative to the tailbud as cell advection in the anterior direction as explained below.

##### Cell mechanics and equation of motion

We model PSM and tailbud cells as spheres with diameter *d*_*c*_ in the 3D domain. We denote the number of cells in the PSM and tailbud at time *t* as *N*(*t*). State variables for these cells are their position **x** in the domain, the phase *θ* of their genetic oscillators and the unit vector **n** representing cell polarity for intrinsic cell movement. Position of cell *i* at time *t* is denoted as **x**_*i*_(*t*) = (*x*_*i*_(*t*), *y*_*i*_(*t*), *z*_*i*_(*t*)) (*i* = 1, 2, 3, …, *N*(*t*)). We describe cellular motion by the following over-damped equation based on the previous study (4):

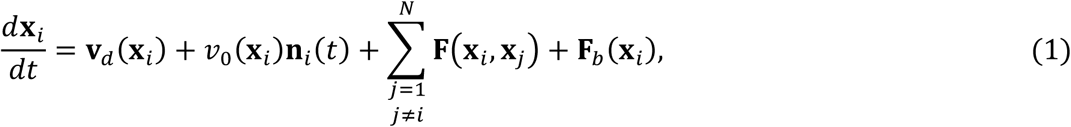

where **v**_***d***_(**x**_*i*_) is cell advection from posterior to anterior, *v*_0_(**x**_*i*_) represents the speed of intrinsic cell movement, **n**_*i*_(*t*) is the cell polarity pointing the direction of intrinsic movement, **F**(**x**_*i*_, **x**_*j*_) is a physical contact force between cells *i* and *j*, and **F**_*b*_(**x**_*i*_) is a boundary force that confines cells within the U-shaped domain. We explain each of these terms below.

##### Elongation, strain rate and cell advection

To introduce tissue elongation and deformations we first discuss the movement of tissue landmarks and cells in the Lab reference frame. Then, we switch to the tailbud reference frame where the tissue elongation results in cell advection from the posterior to the anterior directions.

In the Lab reference frame, segments and cells within do not move. The tailbud, together with cells at that point, moves away from segments with a velocity 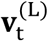. We call *elongation* to this outgrowth of the tissue. As cells differentiate into a segment, the anterior end of the PSM also moves after the tailbud, with a velocity 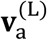. If this velocity matches that of the tailbud 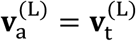, the length of the PSM remains constant. If this velocity differs from that of the tailbud 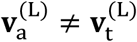, the length of the PSM changes over time, causing a *global tissue deformation*. At these PSM ends we have boundary conditions for the cells velocities: (i) a cell at the tailbud moves with the same velocity as the tailbud, and (ii) a cell at the anterior end of the PSM is at rest, like neighboring cells within a segment, even though the anterior end of the PSM itself is moving in the Lab frame. Within the PSM, cells are subject to an advection velocity field **v**^(*L*)^(**x**) that satisfies these boundary conditions. This velocity field producing *internal local deformations* is caused by internal strains described below. The shape of this velocity field determines the kind of local deformations. For example, a linear velocity field produces uniform deformations, while a non-linear velocity field produces non-uniform deformations, as the piecewise linear function described below in the implementation.

To see the relation between the velocity field and underlying internal strains, let *x*_1_(*t*) and *x*_2_(*t*) be the *x* positions of cells 1 and 2 at time *t* in the Lab reference frame, where we drop the (L) superscript for notational convenience. We assume that their positions are close to each other, so the distance between them Δ*x*(*t*) = *x*_2_(*t*) − *x*_1_(*t*) is small. Due to the presence of an advective velocity field, the velocities of neighboring cells may be different, that is *v*_1_ ≠ *v*_2_, with *v*_1_ = *dx*_1_⁄*dt* = *v*(*x*_1_(*t*)) and *v*_2_ = *dx*_2_⁄*dt* = *v*(*x*_2_(*t*)) = *v*(*x*_1_(*t*) + Δ*x*(*t*)). Thus, during a short time interval Δ*t* these cells change their relative position due to this velocity difference, Δ*x*(*t* + Δ*t*) = *x*_2_(*t* + Δ*t*) − *x*_1_(*t* + Δ*t*) ≈ Δ*x*(*t*) + (*v*_2_ − *v*_1_)Δ*t*. Thus, there is a local strain (Δ*x*(*t* + Δ*t*) − Δ*x*(*t*))⁄Δ*x*(*t*) ≈ (*∂v*⁄*∂x*)Δ*t*, where the velocity field gradient (*∂v*⁄*∂x*) is a local strain rate. Thus, the advection velocity field effectively describes the presence of local strains and determines the local tissue deformation. Although such local deformations make cells move apart from each other along the *x*-axis, intercellular and boundary forces in Eq. (1) constrain cell distances and we do not observe large density fluctuations due to the velocity field gradient, as can be seen in simulations. Furthermore, where local cell density fluctuations do happen, cell addition described below will bring back the density to its average value.

##### Cell advection patterns relative to the tailbud reference frame

Back to the tailbud reference frame, the velocity field is 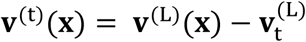, which we refer to as the cell advection pattern **v**_***d***_(**x**) in Eq. (1) and in the main text. We model different advection patterns of the PSM effectively by changing the spatial derivative of the advection speed in subdomains of the tissue. The advection pattern of the PSM may change at a certain developmental stage as we discussed in the main text. Below, we model the spatial profile of cell advection speed and its temporal change. For simplicity, we divide the PSM into two subdomains, namely the anterior PSM ((*x* − *x*_*a*_)/*L* < *x*_*q*_) and the posterior PSM ((*x* − *x*_*a*_)/*L* > *x*_*q*_), and consider different strain rates *∂***v**_***d***_⁄*∂x* for each domain, Fig. 2C. The advection field **v**_*d*_(**x**) in Eq. (1) depends on spatial position along the anterior-posterior axis *x* (*x*_*a*_ ≤ *x* ≤ *L*) and is described as (Fig. 2C):

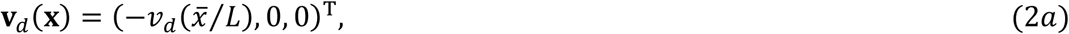

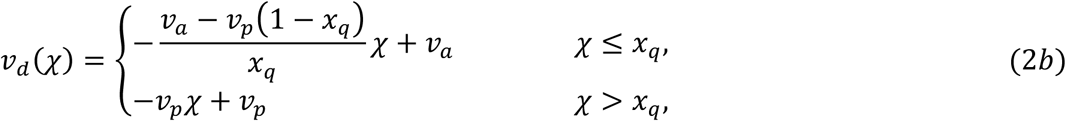

Where 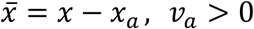 and *v*_*p*_ > 0 are the parameters that determine the advection speed, the superscript T denotes transposition of the vector, and *L* is the length of the PSM *L* = *L*_*x*_ − *x*_*a*_. The slope changes at the position *x*_*q*_ (0 < *x*_*q*_ < 1). Thus, *x*_*q*_ divides the PSM into anterior and posterior subdomains. Within each domain, the strain rate is uniform, whereas it may be different between these two domains. The strain rate, given by the magnitude of the spatial gradient of advection speed, is *v*_*p*_ in the posterior PSM domain and (*v*_*a*_ – *v*_*p*_(1 − *x*_q_))/*x*_q_ in the anterior PSM domain.

##### Temporal change in advection pattern

The advection pattern of the PSM in embryos may change at a certain developmental stage. To model the change in the advection pattern of the PSM, we change the value of *v*_*p*_ in Eq. (2) at time *t* = *t*_*g*_. We assume that for *t* < *t*_*g*_, advection occurs mostly at the anterior part, *v*_*p*_ < *v*_*a*_. For *t* > *t*_*g*_, we assume that advection occurs at the posterior part of the tissue *v*_*p*_ > *v*_*a*_.

##### Gradient of intrinsic cell movement speed

A cell mixing gradient is observed along the anterior-posterior axis of the PSM (4-7). The degree of cell mixing is higher in the tailbud and posterior region of the PSM than anterior region. To model the cell mixing gradient, we assume that the speed of intrinsic cell movement *v*_0_(**x**) in Eq. (1) depends on the spatial position 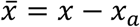 along the anterior-posterior axis (Fig. 2B):

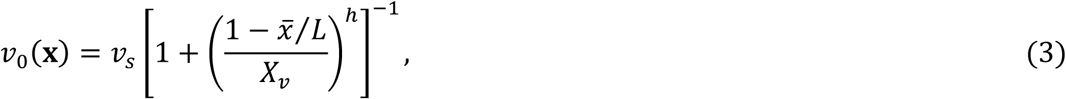

where *v*_*s*_ is the maximum speed at the posterior tip of the tailbud, *X*_*v*_ is the lengthscale of the mobility gradient, and the coefficient *h* determines the steepness of the mobility gradient, Fig. S5A.

##### Cell polarity

The unit vector **n**_*i*_(*t*) in Eq. (1) represents the polarity of cell *i* and determines the direction of intrinsic cell movement. In spherical coordinates, **n**_*i*_(*t*) = (sin*ϕ*_*i*_(*t*)cos*φ*_*i*_(*t*), sin*ϕ*_*i*_(*t*)sin*φ*_*i*_(*t*), cos*ϕ*_*i*_(*t*))^T^ with 0 ≤ *ϕ*_*i*_(*t*) ≤ π and 0≤ *φ*_*i*_(*t*) <2π. We assume random change of cell polarity by letting the polarity angles *ϕ*_*i*_ and *φ*_*i*_ perform a random walk. Specifically, the time evolution of **n**_*i*_ is described by the following Langevin equations for these two polarity angles (4):

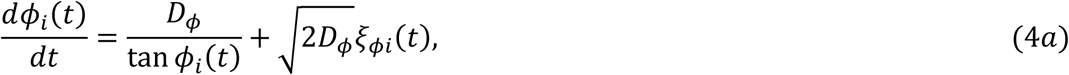

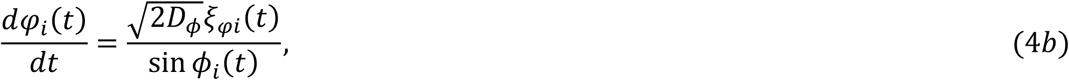

where *D*_*ϕ*_ is the polarity noise intensity. White Gaussian noise *ξ*_*ϕi*_(*t*) and *ξ*_*φi*_(*t*) satisfy ⟨*ξ*_*ϕi*_(*t*)⟩ = 0, ⟨*ξ*_*ϕi*_(*t*)*ξ*_*ϕj*_(*t*′)⟩ = *δ*_*ij*_*δ*(*t* − *t*′), ⟨*ξ*_*φi*_(*t*)⟩ = 0, ⟨*ξ*_*φi*_(*t*)*ξ*_*φj*_(*t*′)⟩ = *δ*_*ij*_*δ*(*t* − *t*′) and ⟨*ξ*_*ϕi*_(*t*)*ξ*_*φj*_(*t*′)⟩ = 0. With the Langevin equations, the polarity vector **n**_*i*_ performs a uniform random walk on the surface of a unit sphere.

##### Intercellular force

For simplicity, we consider a linear elastic force **F**(**x**_*i*_, **x**_*j*_) to model volume exclusion of cells in Eq. (1). If the distance between two cells becomes shorter than cell diameter *d*_*c*_, they repel each other (4, 8):

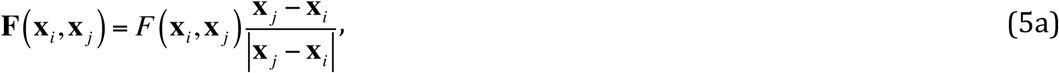

where *F*(**x**_*i*_, **x**_*j*_) is the modulus of intercellular force

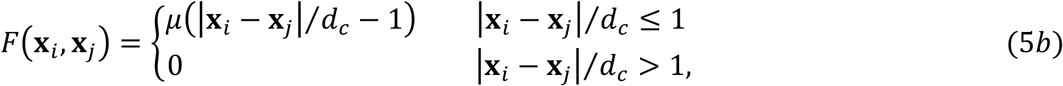

where *μ* > 0 is the coefficient for intercellular force.

##### Boundary force

**F**_*b*_(**x**_*i*_) in Eq. (1) is a confinement force that a cell receives from the boundaries of the domains. Below, we specify this confinement force depending on which tissue domains cells are located in, Fig. S2B.

*In the PSM tubes*: In the PSM regions Ω_*l*_ and Ω_*r*_, we consider boundary force in *y* and *z* directions. The position of cell *i* in the right PSM region Ω_*r*_ can be written as

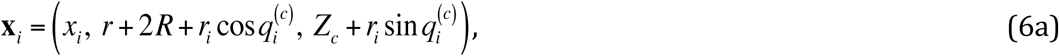

see Fig. S2B. If cell *i* is in the left PSM region Ω_*l*_, its position can be described as

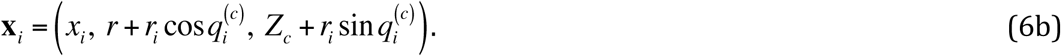

Then, we define frictionless boundary force in the columns as

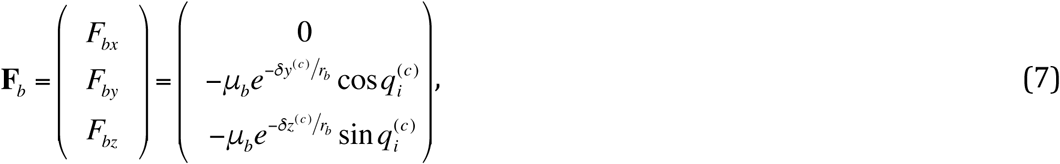

where 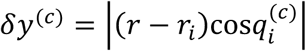 and 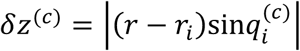.

*In the tailbud half-toroid*: Position of cell *i* within the half-toroidal domain Ω_*t*_ can be expressed as:

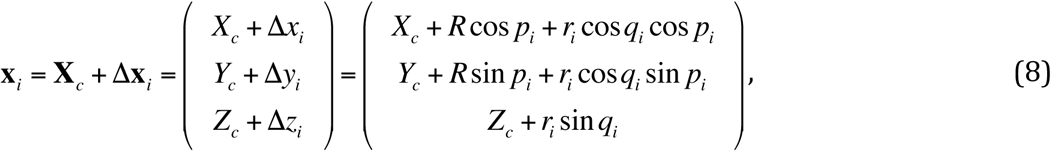

where *p*_*i*_ = tan^−1^(Δ*y*_*i*_/Δ*x*_*i*_), *q*_*i*_ = tan^−1^(Δ*z*_*i*_cos*p*_*i*_/(Δ*x*_*i*_–*R*cos*p*_*i*_)) and *r*_*i*_ = Δ*z*_*i*_/sin*q*_*i*_ (Fig. S2B). We then define the boundary force in the half-toroidal domain as:

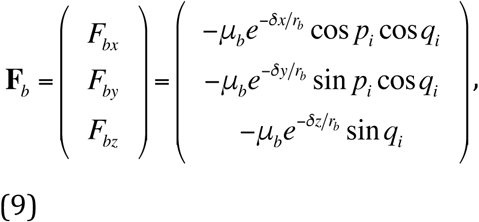

where *μ*_*b*_ is the coefficient and *r*_*b*_ is the length scale of the boundary force, *δx* = |*R*cos*p*_*i*_ + *r*cos*p*_*i*_cos*q*_*i*_ − Δ*x*_*i*_|, *δy* = |*R*sin*p*_*i*_ + *r*sin*p*_*i*_cos*q*_*i*_ − Δ*y*_*i*_|, and *δz* = |*r*sin*q*_*i*_ − Δ*z*_*i*_|.

##### Phase equation

The phase dynamics of genetic oscillators in single PSM cells is described by a phase oscillator model (2, 4, 9, 10):

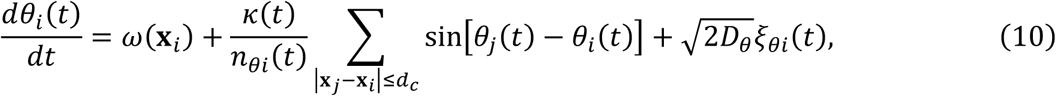

where *θ*_*i*_ is the phase of cell *i, ω*(**x**_*i*_) is the autonomous frequency at position **x**_*i*_, *κ* is the coupling strength, *n*_*θi*_ is the number of neighboring cells interacting with cell *i* and *D*_*θ*_ is the phase noise intensity. Interactions occur between touching cells |**x**_*j*_ − **x**_*i*_| ≤ *d*_*c*_. The coupling strength *κ* may be time-dependent as described below. *ξ*_*θi*_(*t*) is a white Gaussian noise satisfying ⟨*ξ*_*θi*_(*t*)⟩ = 0 and ⟨*ξ*_*θi*_(*t*)*ξ*_*θj*_(*t*′)⟩ = *δ*_*ij*_*δ*(*t* − *t*′).

We assume the frequency profile *ω*(**x**) along the anterior-posterior axis of the PSM to generate traveling phase waves (10-12). It is described as *ω*(**x**) = *ω*_0_*U*(**x**) where *ω*_0_ is the frequency at the posterior tip of the tailbud. For simplicity, we scale the frequency profile with the PSM length *L*. The function *U*(**x**) reads, Fig. 2D (12):

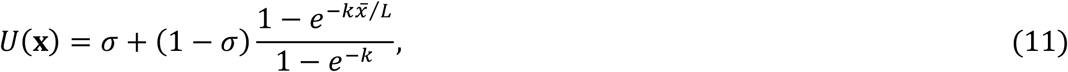

where 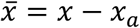, *σ* denotes the difference in the frequency between anterior and posterior ends of the tissue, and *k* determines the shape of the frequency profile, Figs. 2D, S11A.

In Eq. (10), we introduced some simplifications. For example, time delays in intercellular interactions with Delta-Notch signaling play important roles in setting the period of collective rhythms and synchronization (13, 14). However, Eq. (10) does not include it to reduce computational time. In addition, a recent study suggested the presence of a spatial gradient of noise intensity along the PSM (15), but we assume that *D*_*θ*_ is constant across the tissue. Wildtype embryos with Delta-Notch signaling do not spontaneously form defective segments, suggesting that coupling strength is sufficiently strong to overcome phase noise. Therefore, we assume that the phase noise intensity is sufficiently lower than coupling strength and approximate its spatial gradient by the zeroth order term.

The phase of cells anterior to the PSM *x* < *x*_*a*_ is arrested (*i*.*e. dθ*_*i*_ /*dt* = 0 for *x*_*i*_ < *x*_*a*_). Then, we obtain advecting stripes for normal traveling waves in the PSM, Fig. 2E. We consider that the stripes represent segment boundaries and the interval between two consecutive stripes represents segment length as described below.

##### DAPT washout at different time points

In this study, we allow the coupling strength in Eq. (10) to be a time dependent function *κ*(*t*). The value of the coupling strength is varied in the presence or absence of DAPT in embryos. Besides the influence of DAPT, the coupling strength in embryonic cells may change intrinsically with developmental stages due to, for example, gradual changes in Delta and Notch protein levels on cell membrane, and/or changes in contact surface areas between neighboring cells. To model such changes in the coupling strength, we assume the following time dependence:

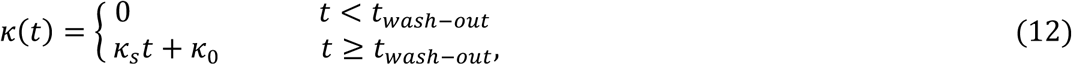

where *t*_*wash-out*_ represents the time at which DAPT is washed out in simulations. In the presence of DAPT *t* < *t*_*wash-out*_, there is no interaction between cells. After DAPT washout at *t* = *t*_*wash-out*_, cells restore coupling immediately and interact with each other at finite rates. For simplicity, we assume that the coupling strength changes linearly with time in Eq. (12) to consider its intrinsic dependence on developmental stages, Fig. S17A. Setting *κ*_*s*_ = 0 in Eq. (12) describes a constant coupling strength *κ*_0_ after DAPT washout. We first analyze resynchronization dynamics with the fixed value of the coupling strength *κ*_0_ by setting *κ*_*s*_ = 0, Figs. 3, 4, S5-S14, S16. Then, we consider a positive value of *κ*_*s*_ > 0 and let the value of the coupling strength gradually increase to study its effect on time to FRS, Figs. 4E, S15, S17.

##### PSM shortening

When we model PSM shortening, we consider that the position of the anterior end of the PSM *x*_*a*_(*t*) changes over time. For simplicity, we assume that the anterior end of the PSM moves in the posterior direction (*x* > 0) at a constant velocity *u*_*a*_ (*u*_*a*_ > 0), Fig. S13A:

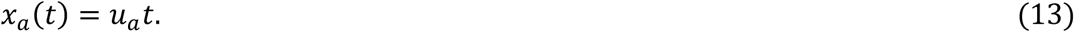

Hence, the length of the PSM becomes shorter as *L*(*t*) = *L*_*x*_ − *u*_*a*_*t*. The speed of the anterior end of the PSM may influence the segment length. For better comparison, we fix the segment length *S* constant for different values of *u*_*a*_ in simulations. For this, we impose the following condition:

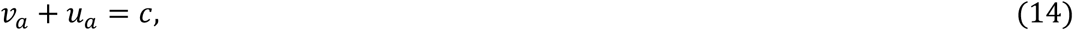

where *v*_*a*_ is the cell advection speed at the anterior end used in Eq. (2) and *c* is a constant. Hence, *v*_*a*_ can be expressed as *v*_*a*_ = *c* − *u*_*a*_. With Eq. (14), the segment length *S* reads *S* = *c* × *T*_*a*_ where *T*_*a*_ is the period of oscillation at the position *x*_*a*_.

##### Cell influx and outflux

Due to cell advection, cells reach the anterior end of the PSM *x* = *x*_*a*_. These cells exit from the domains Ω_*l*_ and Ω_*r*_ at the advection speed |**v**_*d*_(*x*_*a*_)| = *v*_*a*_. We make the simplifying assumption of constant cell density *ρ* ≈ *ρ*_0_. We implement this assumption in simulations by local density dependent cell addition to the PSM and tailbud, as described below:

1. We measure cell density of the left and right PSM *ρ*_*l*_ and *ρ*_*r*_ (*x*_*a*_(*t*) < *x* < *X*_*c*_), respectively, and that of the tailbud *ρ*_*t*_ (*X*_*c*_ < *x*). Then, we compute the difference between the measured density and the target density *ρ*_0_, δ_*l*_ = *ρ*_*l*_ −*ρ*_0_, δ_*r*_ = *ρ*_*r*_ −*ρ*_0_ and δ_*t*_ = *ρ*_*t*_ −*ρ*_0_.
2. If all δ_*l*_, δ_*r*_, and δ_*t*_ are positive, we do not add a cell to the PSM and tailbud when a cell leaves the PSM from its anterior end *x*_*a*_(*t*) due to advection.
3. In contrast, if some of δ_*l*_, δ_*r*_, and δ_*t*_ are negative, we add one cell to the region of which density is smallest. For example, if δ_*l*_ is negative and the smallest, we add a cell to the left PSM. The phase *θ* of the added cell is assigned randomly from the uniform distribution between 0 and 2π. The position of the added cell is randomly assigned within the chosen domain. In the anterior end of the embryonic PSM, cells divide less frequently (unpublished observations). For this reason, we do not add cells in the region *x* ≤ *x*_*a*_ + *ζ* for the left and right PSM. In this study, we set *ζ* = 100 μm. We note that adding cells with random phases in this region would be detrimental for the segmented pattern, given that phase disturbances would not have time to synchronize to their neighbors. The cell polarity angles of the added cell are assigned randomly from the uniform distribution between 0 and 2π for *φ*, and between 0 and π for *ϕ*. The autonomous frequency, speed of intrinsic cell movement and advection speed for the added cell are determined depending on its added position.

##### Change in the PSM radius

We examine how change in the PSM radius over time influences resynchronization, Fig. S16. For simplicity, we assume that the PSM radius *r* decreases uniformly in the U-shaped domain and linearly over time:

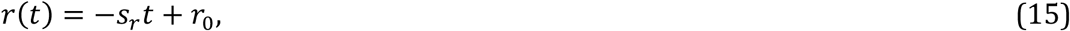

where *s*_*r*_ (*s*_*r*_ > 0) is the magnitude of change in the PSM radius. Because we fix cell density as described in the subsection “*cell influx and outflux*”, the number of cells in the U-shaped domain decreases as the PSM radius becomes narrower. Note that although the thinning of PSM in one direction may cause a length extension in other directions to preserve a volume, we do not consider such extension for simplicity.

To simplify the implementation of the model, we separately update cell positions **x**_*i*_ and the radius size *r*(*t*). When updating *r*(*t*) into *r*(*t*+Δ*t*) by Eq. (15) above, some cells may be left outside the U-shaped domain, *r*_*i*_(*t*+Δ*t*) > *r*(*t*+Δ*t*) where *r*_*i*_(*t*+Δ*t*) is the position of cell *i* in the radial direction defined in Eq. (6) or (8) at time *t*+Δ*t*. For such cells, we correct their positions after the update of PSM radius as: *r*_*i*_(*t*+Δ*t*) → *r*(*t*+Δ*t*) −2*r*_*b*_ where *r*_*b*_ is the lengthscale of boundary force used in Eqs. (7) and (9).

##### Initial condition of simulations

We set the initial position of the anterior end of the PSM *x*_*a*_(0) as *x*_*a*_(0) = 0. We choose the cell number *N*(0) to satisfy the cell density *ρ* = *ρ*_0_ with a given tissue geometry. Initial positions of cells **x**_*i*_(0) (*i* = 1, 2, …, *N*) are within the PSM and tailbud domains. For simulations of embryos without DAPT treatment (*i*.*e*. synchronized initial condition), we use a synchronized initial condition θ_*i*_(0) = 3π/2 for *i* = 1, …, *N*. For resynchronization simulations, the initial phase value of cell *i* θ_*i*_(0) is chosen randomly from a uniform distribution between 0 and 2π. The values of initial cell polarity angles in Eq. (4) are also chosen randomly from the uniform distributions between 0 and π for *ϕ*_*j*_(0), and between 0 and 2π for *φ*_*j*_(0). The autonomous frequency, speed of intrinsic cell movement and advection speed are determined by the initial position **x**_*i*_(0).

##### Parameter values

The values of parameters used in this study are listed in Tables S1 and S2. Cell density and diameter of cells are based on the estimation by ref. (4). The values of parameters that determine intrinsic cell movement and physical forces between cells are set to reproduce experimental data, Fig. S5B (also refer to ref. (4) for details). We use the PSM length within the range reported in ref. (16). The parameters for the frequency profile in Eq. (10) are based on refs. (12, 16). Also, the value of the coupling strength *κ* in Eq. (10) is in the range estimated by refs. (2, 13).

Previous studies estimated noise intensity in the segmentation clock employing different theoretical formalisms (2, 15, 17). The present physical model has two noise sources for phase of oscillation. One is the white Gaussian noise with intensity *D*_*θ*_ in the phase equation (10). The other is cell addition with a random phase value described in the section “*Cell influx and outflux*”. In desynchronization simulations with *κ*(*t*) = 0 in Eq. (10), noise by cell addition alone (*D*_*θ*_ = 0) ruins kinematic phase waves and stripe patterns, Fig. S10A, D. First five stripes are typically recognizable with *D*_*θ*_ = 0, Fig. S10A. With the increase in *D*_*θ*_, the local phase order decays more quickly and stripes are lost earlier, Fig. S10B-D. Because ALD with a saturated dose of DAPT is around five in current experiments, we set *D*_*θ*_ = 0.0013 throughout this study, Fig. S10B. In resynchronization simulations with *κ*(*t*) = *κ*_0_ = 0.07, both FRS and PLD do not depend on *D*_*θ*_ within its examined rage, Fig. S10E. Hence, even if a different value of *D*_*θ*_ is used in resynchronization simulations, only a slight modification of the parameter values would be enough to better fit experimental FRS and PLD. Qualitative aspects of FRS and PLD in simulations do not depend on the value of *D*_*θ*_.

#### Calculation of local phase order

To measure the level of synchrony at local domains along the anterior-posterior axis of the PSM, we introduce a local phase order parameter (18). It is based on the Kuramoto phase order parameter (9) with some modifications in computing average over cells to capture the presence of spatial phase waves in the PSM. For the left PSM Ω_*l*_, we first compute phase order for cells within a thin slice domain Ω_*m*_(*x*) = [*x* + (*m* − 1)Δ*x, x* + *m*Δ*x*] × [0, 2*r*] × [0, 2*r*] for *m* = 1, 2, .., *M* by:

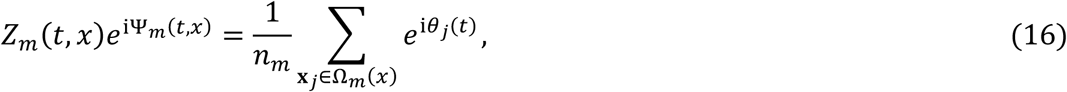

where *n*_*m*_ is the number of cells within the domain Ω_*m*_(*x*), Fig. S3A. We set Δ*x* equal to the cell diameter Δ*x* = *d*_*c*_ so that Eq. (16) measures the phase order for cells in same position in the anterior-posterior axis. *Z*_*m*_ indicates the level of local synchrony of this slice domain. If the phases of cells in the domain are synchronized, *Z*_*m*_ is close to one. In contrast, if they are not synchronized, *Z*_*m*_ is close to zero. Ψ_*m*_ indicates the mean phase of the cells in the slice domain.

This definition of *Z*_*m*_ measures local order with a high spatial resolution along *x*-axis. However, only a few cells are present in each slice, introducing finite size fluctuations. Thus, we define the local phase order parameter at position *x* by taking the average of *Z*_*m*_ over a number *M* of these domains, to smooth out small finite size fluctuations:

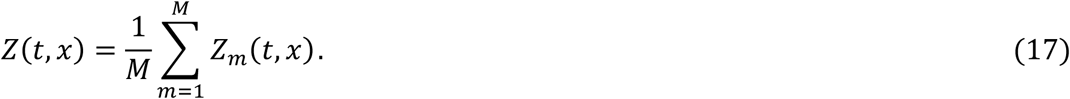

The segment length in the anterior-posterior direction in our parameter sets is ∼5Δ*x*. Therefore, we set *M* = 5 in the calculation of the local phase order Eq. (17). We calculate the local phase order parameter in a similar manner for the right PSM Ω_*r*_. In this case, the domain Ω_*m*_(*x*) in Eq. (16) is Ω_*m*_(*x*) = [*x* + (*m* − 1)Δ*x, x* + *m*Δ*x*] × [2*R*, 2(*R* + *r*)] × [0, 2*r*].

Note that computing local order in thin slices first as in Eq. (16), and then averaging as in Eq. (17), we can capture high local order values even in the presence of a phase gradient. Computing a local order parameter directly in a thicker slab domain of width 5Δ*x* would result in low values of the order parameter in the presence of a phase gradient as in the anterior PSM.

#### Definition of a normal segment boundary in simulations

In computational simulations, we use the phase order parameter Eq. (17) to define normal and defective segment boundaries. We first define a segment boundary by considering simulations for wildtype embryos untreated with DAPT. Such simulations are started from a synchronized initial condition. Then, we describe how to detect normal segment boundaries in resynchronization simulations started from random initial conditions.

In a untreated embryo where cells in the PSM are locally synchronized, kinematic phase waves can be observed across the tissue. In such embryos, the position of a new segment boundary is specified when a wave of gene expression arrives in the anterior end of the PSM. Namely, a position of a segment boundary is determined when the phase of cells near the anterior end of the PSM attains a certain value *ϑ*, Fig. S3B, C.

Based on this, we monitor the mean phase at the anterior end of the PSM *x*_*a*_ to detect when a segment boundary position is set in simulations, Fig. S3A-C. We compute the mean phase Ψ_1_(*t, x*_*a*_) at position *x*_*a*_ at time *t* for Ω_1_(*x*_*a*_) = [*x*_*a*_, *x*_*a*_ + Δ*x*] × [0, 2*r*] × [0, 2*r*] for the left PSM and Ω_1_(*x*_*a*_) = [*x*_*a*_, *x*_*a*_ + Δ*x*] × [2*R*, 2(*R* + *r*)] × [0, 2*r*] for the right PSM by using Eq. (16) (*m* = 1). As noted in Eq. (16), we set Δ*x* as the cell diameter Δ*x* = *d*_*c*_.

We then detect a time *τ*_*i*_ (*i* = 1, 2, …) that satisfies:

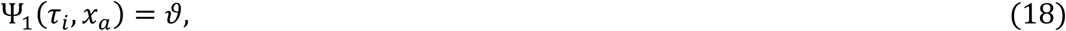

where *ϑ* is a constant that we set *ϑ* = 3*π*/2 without loss of generality in this study, Fig. S3C. For simulations for control embryos where DAPT is not added and, therefore, the level of synchrony is high, *τ* should be the time when the position of the segment boundary *i* is determined, Fig. S3B, C.

##### Identification and numbering of defective segment boundaries

For resynchronization simulations starting from random initial conditions, we modify the above procedure as follows. After detecting the time *τ* when the mean phase of anterior cells becomes *ϑ*, we check the local phase order p_*i*_arameter *Z*(*t, x*_*a*_) defined in Eq. (17) to determine whether these cells can form a normal boundary, Fig. S3D. We define that the anterior cells can form a normal boundary at time *τ* if:

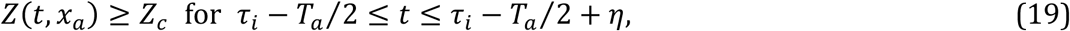

where *T*_*a*_ is the period o_*i*_f oscillation at the anterior end of the PSM *x*_*a*_. When the PSM length does not change during simulation, the anterior period *T*_*a*_ is equal to the period at the tailbud, *T*_*a*_ = 2π/ω_0_. When the PSM becomes shorter over time, we set *T*_*a*_ = 30 min by tuning ω_0_ considering the doppler effect (12, 16). Since *Z*(τ_*i*_−*T*_*a*_/2, *x*_*a*_) is the average local phase order across nearly one segment length at τ_*i*_−*T*_*a*_/2, it evaluates the integrity of the segment boundary and its neighboring inter-boundary regions. To suppress a false detection of a normal segment boundary caused by the fluctuation of *Z*(*t, x*_*a*_), we monitor *Z*(*t, x*_*a*_) in a short time interval with a window size *η* in Eq. (19). We set *η* = 4 min in Eq. (19). By visual inspection of stripe patterns in simulations, we set *Z*_*c*_ = 0.85 for the recovery simulations throughout this paper, see Figs. 3F, S3D, E. Note that this threshold value is simply for detecting a normal segment boundary in simulations. It may be different from the critical value of the order parameter for normal segment boundary formation in actual embryonic tissues.

If the condition (19) is satisfied for *τ*_*i*_, we then specify the segment boundary number. Note that the subindex *i* of *τ*_*i*_ does not specify the segment number in resynchronization simulations due to the fluctuation of the average phase Ψ_1_ for earlier time when cells are not synchronized yet, Fig. S3D. If the previous anterior cell population that satisfied Ψ_1_(*τ*_*i*−1_, *x*_*a*_) = *ϑ* at time *τ*_*i*−1_ (*τ*_*i*−1_ < *τ*_*i*_) was also satisfied the condition (19) and numbered as segment boundary *j*, the current one is numbered as *j*+1. If not, we infer the segment boundary number based on *τ*_*i*_ and anterior period *T*_*a*_. We assign the current segment boundary with the expected segment number:

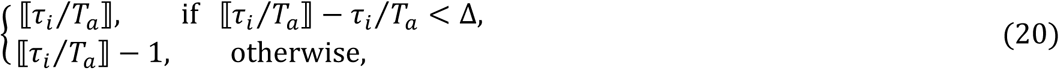

where ⟦*χ*⟧ represents rounding of real number *χ* to the closest integer value. Note that we assign the number ⟦*τ*_*i*_/*T*_*a*_⟧ when the phases of anterior cells become *ϑ* slightly earlier, *τ*_*i*_ > *T*_*a*_⟦*τ*_*i*_⁄*T*_*a*_⟧ − Δ*T*_*a*_ considering the fluctuation after resynchronization, Fig. S3F. In this study, we set Δ = 0.3 in Eq. (20). After detecting the normal segment boundaries and identifying their numbers in this way, we assign all the remaining segment boundaries to be defective.

#### Definition of FRS and PLD in simulations

FRS is defined as *j*_*f*_ − 1 where *j*_*f*_ is the smallest segment boundary number that was determined to be normal. The subtraction of 1 from *j*_*f*_ is to match the definition of FRS for experimental data, see the section “*Whole-mount in situ hybridization and segmental defect scoring*”. PLD is defined as follows. We find the minimum segment boundary number *j*_*p*_ above which all the segment boundaries, including *j*_*p*_, are normal. Then, PLD is *j*_*p*_ − 1. In the example simulation shown in Fig. S3D, E, FRS is 9 and PLD is 13.

#### Calculation of phase vorticity

Vortices are regions in space where the phase values circulate from 0 to 2π around some point. Vorticity could be detected taking a closed path around cells and computing the accumulating change in the phase of neighboring cells around it. However, it is challenging to detect vorticity in a tissue where phase is not continuous in space, but only defined at points where cells are. Besides, there are phase fluctuations that can introduce local variations of phase change. Therefore, here we discretize the closed path in angular steps and average the phase over the resulting domains, Fig. S4. The phase within these domains may grow linearly from 0 to 2π when one turns around a position close to the center of a vortex. Below, we describe the definition of vorticity that is shown in Figs. 3C, S6, S7, S13, S14, S18.

Vortex axis can have different spatial orientations. To detect vortices with different rotating axis, we set several planes in the space and compute phase vorticity at each plane. Then, we project phase vorticity on *x*-axis to obtain its trajectory along the anterior-posterior axis of the PSM. We first choose either the left or right PSM for the calculation of vorticity. Subsequently, we consider the four slices in the PSM, Fig. S4A, B. These slices are two *z*-slices located at *z* = 0 μm and *z* = 2*r*–20 μm (slice 1 and 2, Fig. S4A), and two *y*-slices located at *y* = *y*_0_ μm and *y* = *y*_0_ + 2*r* – 20 μm, (slice 3 and 4, Fig. S4B), where *r* is the radius of the PSM. For the left PSM, *y*_0_ = 0 μm, while for the right PSM *y*_0_ = 2*R* μm where *R* is the tailbud torus radius. The thickness of these four slices is 20 μm (∼2 cell diameters, compare to the 50 μm of PSM diameter). We then project the phase values of cells in each slice to 2D planes, П^(*α*)^ (*α* = 1, 2, 3, 4). The two *x*-*y* planes П^(1)^ and П^(2)^ obtained by the projection of the two *z*-slices (slice 1 and 2) are used to detect vortices with a rotating axis parallel to *z*-axis (Fig. S4A, C). For instance, the *x*-*y* plain П^(1)^ for the right PSM contains cells within the *z*-slice [0, *L*_*x*_] × [2*R*, 2*R* + 2*r*] × [0, 20]. The two *x*-*z* planes П^(3)^ and П^(4)^ obtained by projection of the *y*-slices (slice 3 and 4) are used to detect vortices with a rotating axis parallel to *y*-axis, Fig. S4B. We hardly observed phase vortices with the rotating axis parallel to *x*-axis in simulations. Therefore, we do not consider *y*-*z* planes in the calculation of vorticity.

Next, we set grids (*x*_*s*_, *y*_*u*_) in the planes П^(1)^ and П^(2)^ where *x*_*s*_ = *x*_*a*_ + *s*Δ*x* (*s* = 0, 1, 2, …) and *y*_*u*_ = *y*_0_ + *u*Δ*y* (*u* = 0, 1, 2, …) with the grid size Δ*x* and Δ*y*. Similarly, we set grids (*x*_*s*_, *z*_*u*_) to the planes П^(3)^ and П^(4)^ where *x*_*s*_ = *x*_*a*_ + *s*Δ*x* and *z*_*u*_ = *u*Δ*z*. We chose Δ*x* = 5 μm and Δ*y* = Δ*z* = 2 μm. For each grid point in the plane П^(*α*)^, we compute vorticity *ψ* as follows. Below, we explain the case of П^(2)^ with the grid (*x*_*s*_, *y*_*u*_), Fig. S4A, C. Same calculations were performed in the other planes as well.

We set a circular ring for the grid point (*x*_*s*_, *y*_*u*_) in the plane:

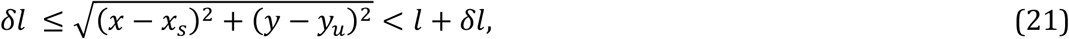

as shown in Fig. S4D, E. We set *δl* = 5.5 μm and *l* = 14 μm. The circular ring is subdivided into 6 domains *V*_*i*_ (*i* = 0, 1, 2, …, 5) by angles π/3 measured counterclockwise from the *x*-axis. We then compute average phase 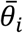 over cells within each subdomain 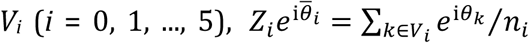 where *n*_*i*_ is the number of cells in *V*_*i*_. If there is no cell within one of the subdomains *V*_*i*_ (*n*_*i*_ = 0), we do not compute the vorticity for the grid point (*x*_*s*_, *y*_*u*_) and set *ψ* = 0.

To detect a vortex with clock-wise rotation, we permutate 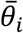 based on their values (Fig. S4D, E): 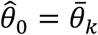 where 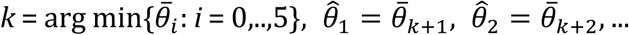, and 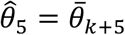, (mod 6). For a vortex with counter clock-wise rotation, we permutate 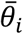 as: 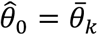 where 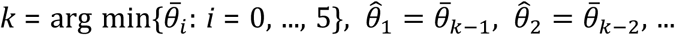, and 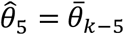, where a negative value of *k* − *j* (*j* =) 1, 2, …, 5) should be replaced as −1 → 5, −2→ 4, …, and −5 → 1.

We assume that when a phase vortex is present near the grid point (*x*_*s*_, *y*_*u*_), 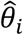 for the grid point increases linearly with *i*, Fig. S4D, F. To detect this linear increase of 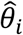, we compute the correlation coefficient *α* defined as:

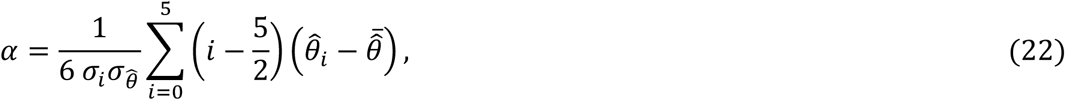

where 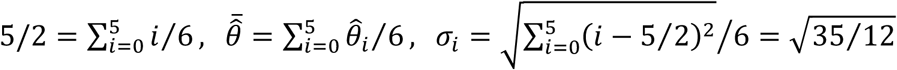, and 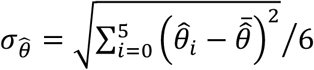. A value of the correlation coefficient *α* close to one means that the phase increases linearly along a perimeter of a circle, indicating the existence of a phase vortex, Fig. S4D, F. If the correlation coefficient is larger than a threshold *α* ≥ *α*_0_, we consider that the phase value consistently increases along the circumference of the ring and rotates along the *z*-axis. In this case we define vorticity for the grid point as 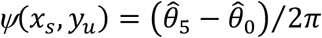, Fig. S4F. If *α* < *α*_0_, we set *ψ*(*x*_*s*_, *y*_*u*_) = 0 to exclude false positive detection of a vortex by fluctuation of phase values. We used *α*_0_ = 0.75 throughout the article. After calculating vorticity for each grid point, we project *ψ*(*x*_*s*_, *y*_*u*_) to *x*-axis. We use maximum projection, 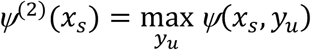.

By performing same calculations for the remaining three planes, we obtain {*ψ*^(1)^(*x*_*s*_), *ψ*^(2)^(*x*_*s*_) *ψ*^(3)^(*x*_*s*_), *ψ*^(4)^(*x*_*s*_)} for position *x*_*s*_, top panel of Fig. S4I. Finally, we take their maximum value and adopt it as the vorticity at the position 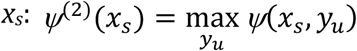, bottom panel of Fig. S4I. For visualization, we made a density plot as a kymograph by using the data (*x*_*s*_, *t, ψ*^(max)^).

#### Quantification of single and double defects

In both experiments and simulations, we sometimes observe that a defective segment boundary appears either only left or right side of an embryo, which we refer to as a single defect. We also observe another instance where left and right boundaries are both defective, which we call double defect. To examine whether the theory can account for the emergence of single defects in embryonic experiments, we compute the fraction *F*_*s*_ of single defects defined below and compare it between simulations and experiment.

##### Experimental data

We first counted the total number *N*_*t*_ of defective segment boundary loci for an embryo. We then counted the number of single defects *N*_*s*_ and computed the fraction *F*_*s*_ = *N*_*s*_/*N*_*t*_. When we compare the experimental data with simulation data, Fig. S20D, we measure *N*_*t*_ and *N*_*s*_ after segment 9 that marks the onset of resynchronization, Fig. S1B.

##### Simulation data

We defined normal and defective segment boundaries based on the local phase order at the anterior end of the PSM *Z*(*t, x*_*a*_) as described in the previous section “*Definition of a normal segment boundary in simulations*”. For single realizations of simulation, we counted the total number of defective segment boundary loci *N*_*t*_ and the number of single defects *N*_*s*_ appeared posterior to the segment 9 as in experimental data. Then, we computed the fraction, *F*_*s*_ = *N*_*s*_/*N*_*t*_, Fig. S20D.

#### Implementation for numerical simulations

We solved Eqs. (1) and (10) with the Euler-Maruyama method with the time step for integration δ*t* = 0.01 min. Custom simulation codes were written in C language. Movies of numerical simulations, calculations of local phase order and vorticity, and analysis of left-right segment boundary defects were done with custom Mathematica (Wolfram) codes.

#### Segment statistics from the spatial distribution of defective segments

The distribution of defects along the embryonic axis can be parametrized in different ways. Here, we introduce complementary pictures, one relies on the fraction of left and right defects, and the other on fractions of single and double defects. Using a random defect hypothesis we show how to relate these pictures with probability theory. Taking into account the spatial distribution of defects we compute ALD, PLD and FRS from these statistics, and we predict and test the values of the fraction of single defects per embryo.

##### Segment state variables

We introduce a state variable that accounts for the presence or absence of a defective segment *S*_*ik*_(*x*), where *i* labels the embryo, *k* = {*l, r*} labels the side of the embryo and *x* is the segment locus along the axis. We consider *N* embryos with *M* + 1 segment boundaries, so *i* = 1, …, *N* and *x* = 0, …, *M*. The state variable takes the values zero and one depending on whether the segment is normal or defective.

##### Segment defect distribution on both sides of the embryo

When a segment locus is defective on the left (right) side of the embryo we call it a left (right) defect independently of the state of the other side. We define left and right defect distributions taking the population average of segment state variables,

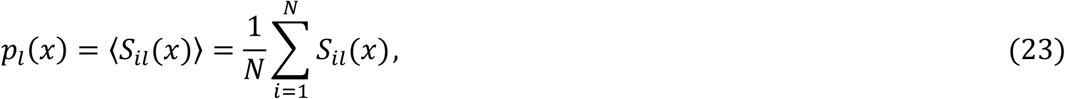

and

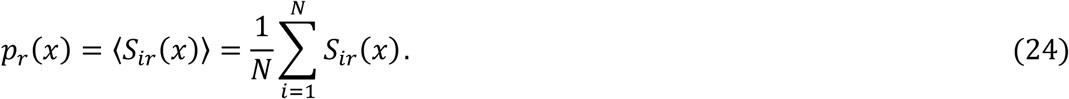

These spatial distributions of defective segments on the left and right sides of embryos are very similar, for different DAPT washout timing, Fig. S19A. This agreement between left and right distributions supports the assumption *p*_*l*_(*x*) = *p*_*r*_(*x*) = *p*(*x*). In the following sections, we use probability theory to compute ALD, PLD and FRS from this spatial distribution of defective segments.

##### Probabilistic calculation of ALD

Let *q*_*a*_(*x*_*a*_) be the probability to find an ALD at position *x*_*a*_. *q*_*a*_(*x*_*a*_) can be expressed in terms of *p*(*x*) as:

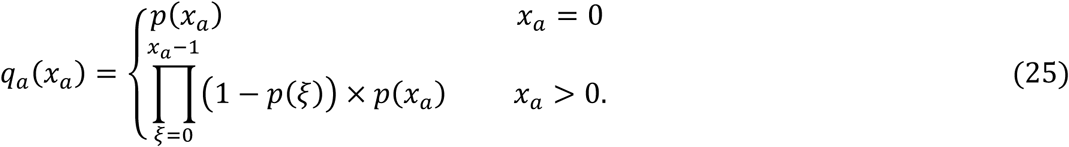

The first factor 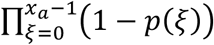 in the second line represents the probability of normal segment boundaries from position *x* = 0 to *x* = *x*_*a*_ – 1. The second factor *p*(*x*_*a*_) is the probability of a defective segment boundary at position *x* = *x*_*a*_. The resulting ALD distribution *q*_*a*_(*x*_*a*_) presents a clear peak at the onset of the defective region, Fig. S19B. The ALD is then calculated as the mean value for this probability distribution,

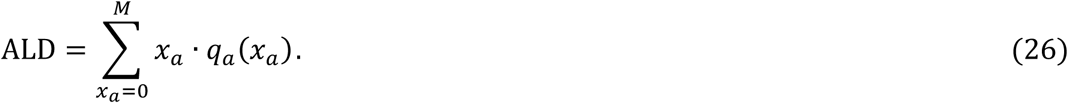

##### Probabilistic calculation of PLD

Let *q*_*p*_(*x*_*p*_) be the probability of PLD at position *x*_*p*_. It can be written as:

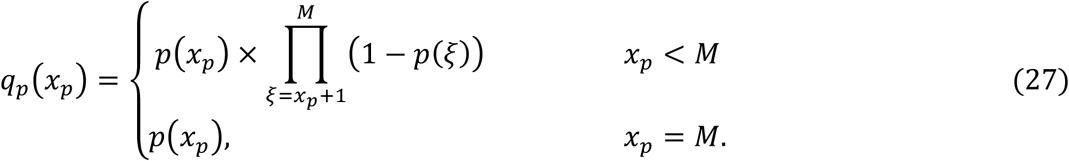

The first factor *p*(*x*_*p*_) in the first line is the probability of a defective boundary at *x*_*p*_. The second factor 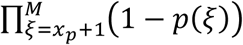 represents the probability that all the remaining segment boundaries posterior to *x*_*p*_ are normal. The resulting PLD distribution *q*_*p*_(*x*_*p*_) peaks at the end of the defective region, Fig. S19B. The PLD can be written as the mean value for this distribution,

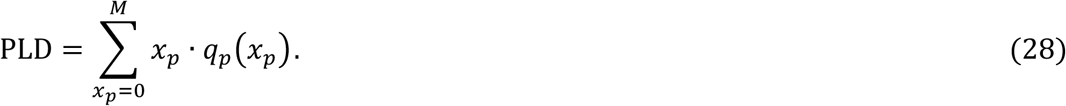

##### Probabilistic calculation of FRS

Occurrence of a recovered segment is conditioned to the previous occurrence of defective segments. In this study, we measure the FRS after the desynchronization phase. The desynchronization phase is determined based on the distribution of defective segments, Fig. S1B. Suppose that the desynchronization phase ends by the formation of segment *s*_*d*_. We define FRS as the first normal segment after *s*_***d***_ − 1. This is the definition we use to measure the embryonic FRS. With this definition of FRS, the probability *q*_*f*_(*x*_*f*_) of the first normal segment boundary at locus *x*_*f*_ is:

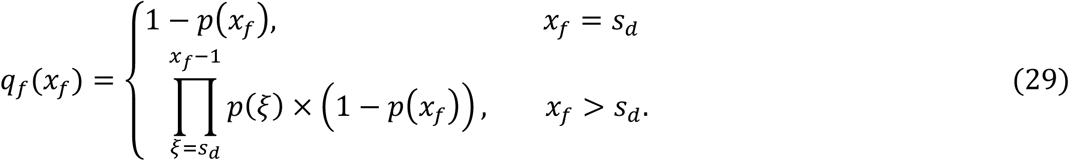

The first factor of the second line represents the probability for all the segment boundaries between *s*_***d***_ and *x*_*f*_ − 1 to be defective. The second factor is the probability for a normal segment boundary to form at *x*_*f*_.

To compute *q*_*f*_(*x*_*f*_), we set *s*_***d***_ = 9 as in the main text, Fig. S1B. The resulting distribution *q*_*f*_(*x*_*f*_) has a peak that precedes that of PLD and partly overlaps with it, Fig. S19B. From these results, the FRS then can be expressed as

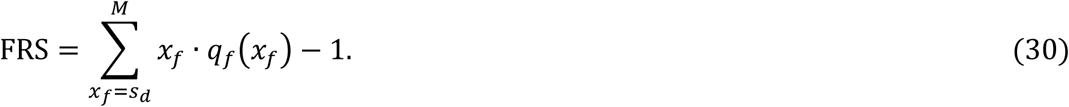

In Eq. (30), we subtract 1 because FRS is defined with the anterior boundary of the first normal segment after *s*_***d***_ − 1, see definition of FRS for experimental data. *q*_*a*_, *q*_*p*_ and *q*_*f*_ calculated with Eqs. (25), (27) and (29) agree well with the direct measurement of these distributions, Fig. S19B. Furthermore, expressions obtained for ALD, PLD and FRS from the spatial distribution of defects *p*(*x*), Eqs. (26), (28) and (30), are in very good agreement with direct measurements of these quantities, Fig. S19C.

##### The spatial distribution of single and double defects

In a complementary framework, we introduce double defects, single defects and normal segments. Double defects occur when both sides of the embryo at a given locus *x* are defective. Single defects occur when at a given locus there is a defect either left or right, but not on the other side. Normal segments have no defects on either side. Taking population averages, we can define the double defect distribution

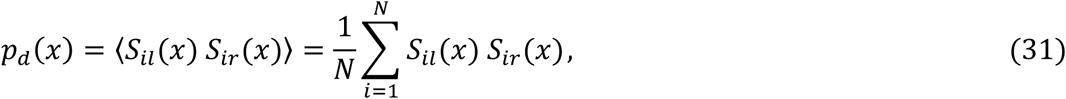

the single defects distribution

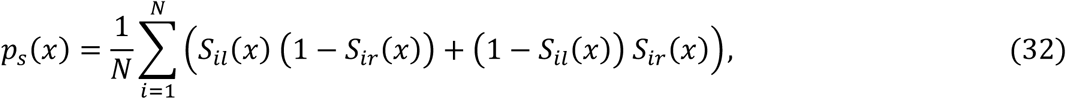

and the normal segments distribution

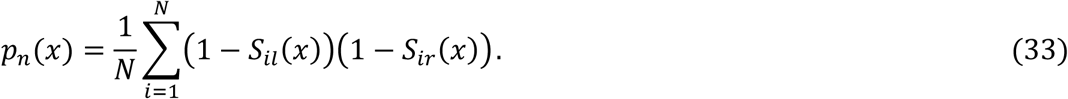

These distributions can be related to the left/right framework introduced above. Key to this is that the two sides of the anterior PSM are physically unconnected, separated by another tissue –the notochord. It has been shown that interfering with Retinoic Acid, which controls somitogenesis bilateral symmetry by shielding asymmetric cues, results in asymmetric left/right segmentation and clock waves (19). This indicates that segmentation clock oscillations are independent in the left and right sides of the PSM. In the physical model, we tacitly assume this independence since there is no coupling between oscillators on one side and the other. A consequence of this left/right independence should be a vanishing covariance of the segment defect variables at opposite sides of the embryo (20),

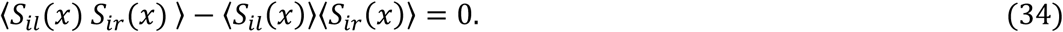

The two terms in this covariance can be written using the segment defect distributions of the two frameworks,

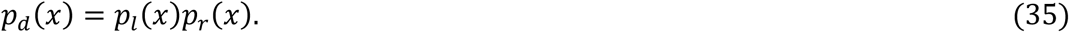

Similarly, we obtain for single defects

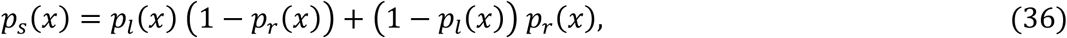

and for normal segments

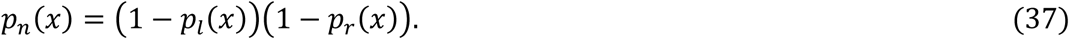

We can further simplify these expressions with the assumption *p*_*l*_(*x*) = *p*_*r*_(*x*) = *p*(*x*). The axial distribution of double defects *p*_***d***_(*x*) gradually grows to a plateau at the onset of the defective region and then decays at its end, while the distribution of single defects *p*_*s*_(*x*) peaks both at the onset of the defective region and at its end, Fig. S20A. The good agreement observed between direct measurement of *p*_***d***_(*x*) and *p*_*s*_(*x*) and results obtained from *p*(*x*) together with probabilistic arguments, provides a test for the vanishing of the covariance and left/right independence, Fig. S20A.

##### Fraction of single defects from the axial distribution of defects

As a further test of the reach of the defect distribution *p*(*x*), we use it to compute the fraction of single defects in a population of embryos. We first count the number of double and single defects in a given embryo *i*

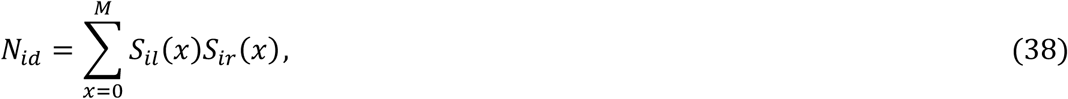

and

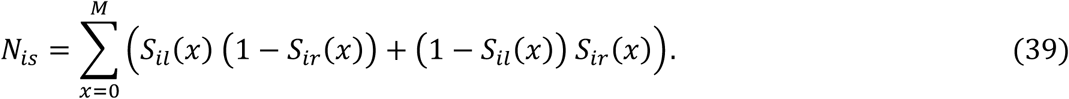

The total number of defects in embryo *i* is *N*_*it*_ = *N*_*is*_ + *N*_*i****d***_. Then, we take the population average of these quantities. The average number of double defects in the population is

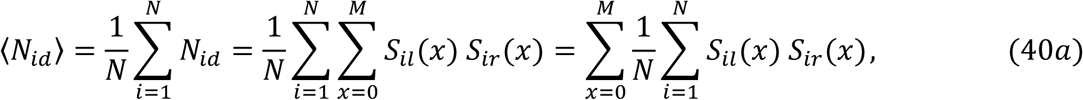

So

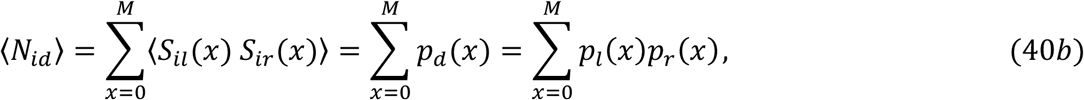

using left/right independence. Similarly, the average number of single defects ⟨*N*_*is*_⟩ is

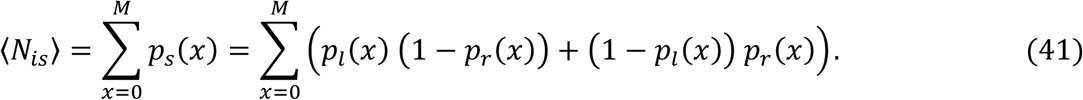

We define the fraction of single to total defects in an embryo

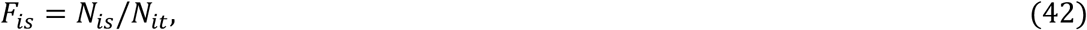

and its population average

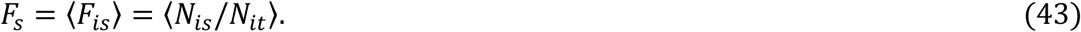

If the coefficient of variation of the total number of defects in the population is small, we can approximate

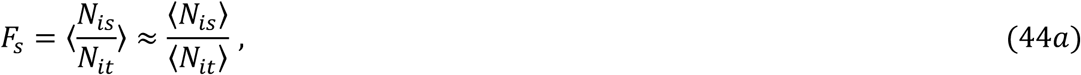

so

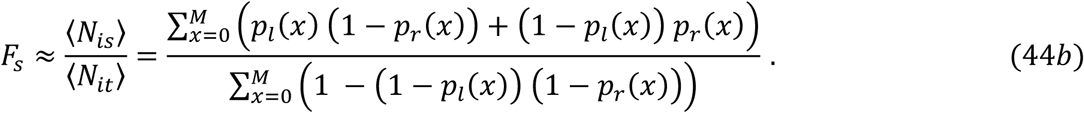

Using the left/right symmetry of defect distributions *p*_*l*_(*x*) = *p*_*r*_(*x*) = *p*(*x*), we can further simplify this expression

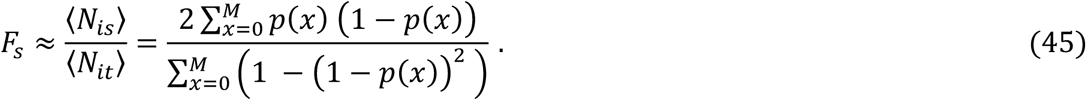

The good agreement observed between this probabilistic calculation and direct measurement of *F*_*s*_ counting single defects in individual embryos, Fig. S20C, suggests that the recovery of segments after DAPT washout occurs independently between left and right sides of the PSM.

Finally, the physical model of the PSM reproduces both the single and double defect distributions, Fig. S20B, and the dependence of *F*_*s*_ on DAPT washout timing, Fig. S20D.

##### Conclusion

Taken together, these results suggest that knowledge of *p*(*x*) is enough to compute some of the key observables that we use to quantify segment recovery, such as ALD, PLD, FRS and *F*_*s*_. Together with the result that the physical model of the PSM reproduces *p*(*x*) at different washout timings, Fig. 4F, this is evidence for the breadth of the physical model results and predictions.

